# Tertiary folds of the SL5 RNA from the 5′ proximal region of SARS-CoV-2 and related coronaviruses

**DOI:** 10.1101/2023.11.22.567964

**Authors:** Rachael C. Kretsch, Lily Xu, Ivan N. Zheludev, Xueting Zhou, Rui Huang, Grace Nye, Shanshan Li, Kaiming Zhang, Wah Chiu, Rhiju Das

## Abstract

Coronavirus genomes sequester their start codons within stem-loop 5 (SL5), a structured, 5′ genomic RNA element. In most alpha- and betacoronaviruses, the secondary structure of SL5 is predicted to contain a four-way junction of helical stems, some of which are capped with UUYYGU hexaloops. Here, using cryogenic electron microscopy (cryo-EM) and computational modeling with biochemically-determined secondary structures, we present three-dimensional structures of SL5 from six coronaviruses. The SL5 domain of betacoronavirus SARS-CoV-2, resolved at 4.7 Å resolution, exhibits a T-shaped structure, with its UUYYGU hexaloops at opposing ends of a coaxial stack, the T’s “arms.” Further analysis of SL5 domains from SARS-CoV-1 and MERS (7.1 and 6.4-6.9 Å resolution, respectively) indicate that the junction geometry and inter-hexaloop distances are conserved features across the studied human-infecting betacoronaviruses. The MERS SL5 domain displays an additional tertiary interaction, which is also observed in the non-human-infecting betacoronavirus BtCoV-HKU5 (5.9-8.0 Å resolution). SL5s from human-infecting alphacoronaviruses, HCoV-229E and HCoV-NL63 (6.5 and 8.4-9.0 Å resolution, respectively), exhibit the same coaxial stacks, including the UUYYGU-capped arms, but with a phylogenetically distinct crossing angle, an X-shape. As such, all SL5 domains studied herein fold into stable tertiary structures with cross-genus similarities, with implications for potential protein-binding modes and therapeutic targets.

**Significance:** The three-dimensional structures of viral RNAs are of interest to the study of viral pathogenesis and therapeutic design, but the three-dimensional structures of viral RNAs remain poorly characterized. Here, we provide the first 3D structures of the SL5 domain (124-160 nt, 40.0-51.4 kDa) from the majority of human-infecting coronaviruses. All studied SL5s exhibit a similar 4-way junction, with their crossing angles grouped along phylogenetic boundaries. Further, across all species studied, conserved UUYYGU hexaloop pairs are located at opposing ends of a coaxial stack, suggesting that their three-dimensional arrangement is important for their as-of-yet defined function. These conserved tertiary features support the relevance of SL5 for pan-coronavirus fitness and highlight new routes in understanding its molecular and virological roles and in developing SL5-based antivirals.

**Classification:** Biological Sciences, Biophysics and Computational Biology

## Introduction

In the *Coronaviridae* family, only seven species, SARS-CoV-2, SARS-CoV-1, MERS, HCoV-HKU1, HCoV-OC43, HCoV-229E, and HCoV-NL63, are known to infect humans, and all are derived from the alpha- and betacoronavirus genera (1). While effective vaccines are available against SARS-CoV-2, the likelihood of future coronavirus pandemics motivates efforts to understand the conserved features of coronavirus machinery. The SARS-CoV-2 proteome and its interactions with host proteins have been well studied structurally, enabling the design of therapeutics targeting host and viral proteins (2–4), but the RNA genome has been relatively understudied. On one hand, the monumental scientific response to COVID-19 has resulted in extensive mapping of SARS-CoV-2 RNA secondary structure (5–10). On the other hand, to date, only a few small fragments of the RNA genome — stem-loop 1 in complex with non-structural protein 1, stem-loop 2, and stem-loop 4 from the 5′UTR; the frameshift stimulation element; and the stem-loop 2 motif from the 3′UTR — have been characterized in three dimensions (11–18). While current designs of RNA-targeting therapeutics are often based on the RNA’s two-dimensional base-pairing pattern, known as RNA secondary structure (12, 19–21), three-dimensional (3D) structures are necessary for structure-guided design of many classes of therapeutics (5, 22, 23).

The 5′ proximal region of the SARS-CoV-2 genome is a highly structured and functionally important genomic locus (24, 25). This region is divided into secondary structure domains termed “stem-loops,” with multiple stem-loops predicted to be present in the 5′ proximal region across the coronavirus family (25–27). Stem-loop 5 (SL5, residues 150-294) contains the start codon of open reading frame 1a/b (ORF1a/b, residues 266-268). Additionally, phylogenetic covariance analysis and chemical probing experiments have shown that SL5′s secondary structure forms a four-way junction that sequesters the genome’s start codon within one of its helical stems (5–8, 10). Beyond SARS-CoV-2, the multi-way junction and the sequestration of the start codon in a stem are conserved features across most coronaviruses (6, 27, 28), despite large sequence divergence from SARS-CoV-2 (average sequence identity of 51% for the NCBI Reference Sequences for alpha- and betacoronaviruses; **Supplemental Figure S1**). While conserved tertiary structural motifs would be attractive targets for antiviral targeting and help pinpoint SL5 function, it is unknown whether SL5 forms a stable 3D structure in solution. Computational algorithms for RNA tertiary structure predict a wide range of 3D conformations (9, 29). Furthermore, SL5 stem lengths, internal loops, and sequences at the four-way junction are not conserved across coronaviruses (**Supplemental Tables S1, S2**), leading to further uncertainty as to whether SL5 robustly forms a tertiary fold.

SL5 has been proposed to play a role in protein-RNA or RNA-RNA binding because of its conserved hexaloops, characterized by 5′-UUYYGU-3′ (Y=C,U) repeat loop motifs (27). This motif is conserved across most alpha- and betacoronaviruses, with the exception of betacoronaviruses in the *Embecovirus* subgenus, including the human-infecting HCoV-HKU1 and HCoV-OC43, which instead harbor a repeat loop motif elsewhere in their genomes (27). While the function of the UUYYGU hexaloop is unknown, it has been proposed to serve as a packaging signal in coronaviruses (27), and the 5′ proximal region has been confirmed to contain a packaging signal in one alphacoronavirus, the pig-infecting transmissible gastroenteritis coronavirus (30). The start codon’s occlusion in SL5′s secondary structure suggests that the folded form of SL5 does not enhance translation, and the deletion of SL5 or sub-structures of SL5 reduces its translation efficiency (31). However, the full role of SL5 structure in viral packaging or viral translation remains unclear and its role in other viral functions, such as viral genome replication, has not been ruled out.

To set a foundation for structure-function relationships, we set forth to characterize conserved structural elements among homologous constructs to distinguish species-specific and genus-specific features of SL5. Comparative structural biology studies have been widely pursued to assess the structures of multiple coronavirus spike proteins (32, 33). Such tertiary structural comparisons in RNA-only structures, however, have been limited. Among viral RNA structures, previous comparisons, conducted by NMR or X-ray crystallography, were limited to the comparison of two homologs (34–36). Structural comparisons for coronavirus RNA genomes have been limited to RNA secondary structure, including analysis of the 5′ UTR (26, 27) and the frameshift stimulation element (37), rather than comparisons of tertiary structure. Cryogenic electron microscopy (cryo-EM) offers the possibility of more routinely characterizing RNA elements across multiple homologs, particularly when integrated with biochemical secondary structure determination and automated computer modeling, such as in the Ribosolve pipeline (38).

Herein, we conduct a comparative study of SL5′s tertiary fold across six coronaviruses. First, we report a tertiary structure of a 124 nt portion (40.0 kDa) of SARS-CoV-2 SL5, obtaining a 4.7 Å map of the domain. Contrary to the many possible tertiary conformations observed in *de novo* computational modeling (9, 29), we observe a well resolved T-shaped conformation, where the larger stem-loops, SL5a and SL5b, base stack perpendicularly to the SL5-stem, with the short SL5c stem-loop jutting out from the T-shape. We then investigate 3D structural homology by resolving the structures of the SL5 domain from five additional coronaviruses. We find that all studied alpha- and betacoronaviruses share the same base stacking geometry, but there are genus-specific inter-helical angles. Additionally, within betacoronaviruses, the merbecovirus orthologs show tertiary features not observed in sarbecoviruses. Even though half of the SL5 domains characterized exhibit structural heterogeneity, every SL5 domain examined here populated a conformation in which the UUYYGU hexaloop sequences were displayed at opposing ends of a coaxial stack of conserved length. These structures and the analysis of 3D structural feature conservation suggest hypotheses for the function of the SL5 RNA element and may aid the rational design of therapeutics.

## Results

### SARS-CoV-2 SL5 domain secondary structure

Our approach to SL5 structural characterization followed an established protocol (38), which integrates biochemical determination of secondary structure, tertiary fold determination with cryo-EM, and automated coordinate building in Rosetta. The secondary structure of SL5 was determined using multidimensional mutate-and-map chemical mapping as read out by sequencing (M2-seq) (39) with an updated “scarless” procedure such that, at the time of chemical modification, the sequence of interest did not contain any appendages. The secondary structure was found to contain a four-way junction without unpaired nucleotides, two UUYYGU hexaloops, and a GAAA tetraloop (**Figures 1A-B**). This secondary structure was also recovered from a “large-library” M2-seq approach, which used a synthesized library of mutants instead of error-prone PCR, and also matches previous *in vivo* and *in vitro* experimental secondary structure determinations (**Supplemental Figure S2**) (5–8, 10). The secondary structures are consistent, with only minor differences in base-pairing at the terminal stem (residues 159-165 and 277-282) where our construct was excised out of the SARS-CoV-2 genome.

**Figure 1:**
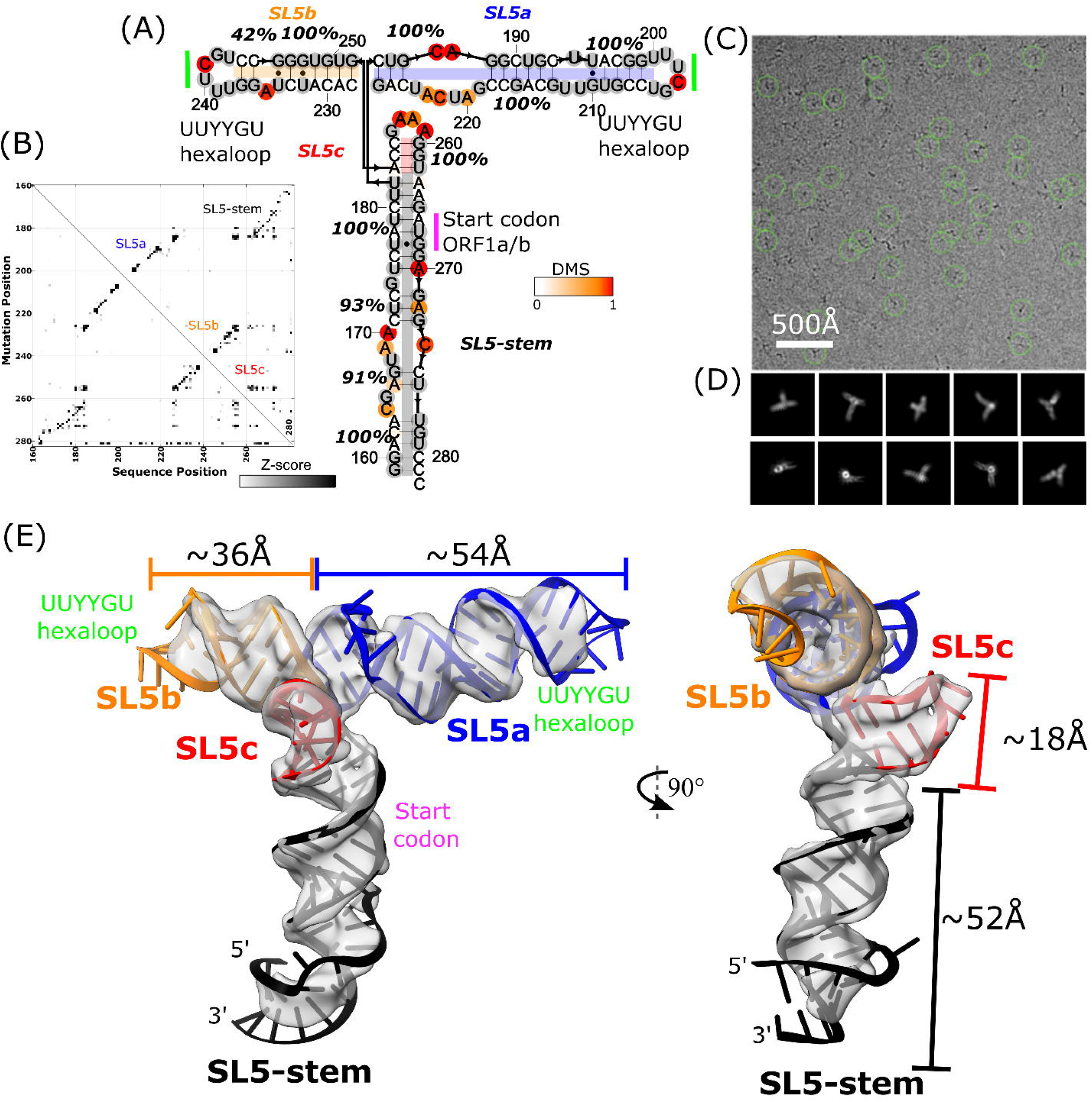
The 3D global fold of SARS-CoV-2 SL5. **(A)** M2-seq-derived secondary structure of SARS-CoV-2 SL5. Stem confidence estimates (100 bootstraps) are given as percentage values and nucleotides are colored by DMS reactivity. **(B)** M2-seq Z-score plot, where the increases in reactivity across the molecules (x-axis) upon mutations (y-axis) are displayed in black. **(C)** Representative micrograph and **(D)** 2D class averages for the cryo-EM dataset of SARS-CoV-2 SL5. **(E)** The 4.7 Å cryo-EM map, displayed in gray, with a representative model. The model was obtained by using the M2-seq derived secondary structure and auto-DRRAFTER followed by refinement with ERRASER2. SL5 helices are colored in black (SL5-stem), blue (SL5a), orange (SL5b), and red (SL5c). The locations of the start codon (magenta) and UUYYGU hexaloops (lime) are also labeled.

### SARS-CoV-2 SL5 exhibits a stable 3D tertiary fold

Cryo-EM image reconstruction of the SARS-CoV-2 SL5 domain (residues 159-282, 124 nt, 40.0 kDa) shows a single, well defined 3D structure resolved to 4.7 Å resolution (**Figures 1C-E, Supplemental Figure S3**). Four helices extend from one junction, all with clear major and minor grooves. The approximate lengths of these helices align with the expected lengths of the stems, as implied from the M2-seq secondary structure, enabling the unambiguous identification of SL5c as the shortest stem and SL5b as the medium length stem (**Figure 1E, Supplemental Table S3**). The junction is well resolved, revealing two pairs of coaxially stacked helices and a hole at the junction that clearly demarcates the backbone connectivity between helical pairs (**Figure 3A**). This connectivity also enables the unambiguous assignment of SL5a and the SL5-stem into the map (**Figure 1E**).

### SARS-CoV-2 SL5 domain 3D structure

Guided by the 4.7 Å map and M2-seq secondary structure, integrative modeling was conducted using auto-DRRAFTER, which is specifically suited for medium to low resolution RNA-only cryo-EM maps (38). Auto-DRRAFTER enumerates helical placements in the cryo-EM density and scores models by a combined biophysical and fit-to-map score. Acknowledging that, in cryo-EM maps lower than 3.5 Å resolution, nucleotide bases cannot be precisely placed, we use the top ten models, ranked by the auto-DRRAFTER scoring, as a representation of experimental uncertainty. Additionally, in this study, we identified multiple secondary structures from M2-seq, the literature (6), and predictions from EternaFold (40). All 3D modeling that converged across multiple computational runs and that fit well into the map was collected into a coordinate ensemble. This resulted in an ensemble of 30 models with a mean pairwise root-mean-squared error (r.m.s.d) convergence of 2.5 Å; slight differences in secondary structure are reflected in the ensemble but did not significantly affect the tertiary fold (**Supplemental Figure S4, Supplemental Table S4**). Auto-DRRAFTER converged on a T-shaped conformation wherein the SL5-stem forms the “leg” of the T-shape, SL5a and SL5b stack end-to-end to form the perpendicular “arms’’ of the T-shape, and SL5c juts out of the plane of the T-shape at the junction. This automated modeling is in agreement with the global fold that we visually inferred from the map features alone. The UUYYGU hexaloops of SL5a and SL5b are positioned at the “hands’’ of the T-shape, positioned 84.2±0.8 Å away from each other (N=30, **Figure 1E**, **Table 1**, **Supplemental Figure S5**). The two pairs of coaxially stacked helices, SL5a:SL5b and SL5-stem:SL5c have an inter-helical angle of 84.3±0.5° but do not form any significant tertiary interactions (N=30, **Figure 1E**, **Table 1**, **Supplemental Figure S5**).

**Table 1:** 3D features of the SL5 domain of coronaviruses.

| Genus | Subgenus | Species | Distance between UUYYGU hexaloops (Å)<sup>1</sup> |  | Inter-helical angle (°)<sup>2</sup> |
| --- | --- | --- | --- | --- | --- |
| *Betacoronaviridae* | *Sarbecovirus* | SARS-CoV-2 | 84.2±0.8 |  | 84.3±0.5 |
|  |  | SARS-CoV-1 | 82±1 |  | 86.8±0.5 |
|  | *Merbecovirus* | MERS | 80±2 |  | 82±2 |
|  |  | BtCoV-HKU5 | 80±4 |  | 86±2 |
| *Alphacoronaviridae* | *Duvinacovirus* | HCoV-229E | 92.4±0.9 |  | -121.3±0.2 |
|  |  |  | SL5b ↔ SL5c | 94.3±0.8 |  |
|  |  |  | SL5a↔SL5c | 49.1±0.8 |  |
|  | *Setracovirus* | HCoV-NL63 | 95, 90<sup>3</sup> |  | -120<sup>3,4</sup> |
|  |  |  | SL5b ↔ SL5c | 85, 65<sup>3</sup> |  |
|  |  |  | SL5a↔SL5c | 45, 50<sup>3</sup> |  |

<sup>1</sup> Distance is between the apical loops of SL5a and SL5b unless otherwise specified. Distance is defined as the distance between the centroid of the C1' atoms of each loop. Error is standard deviation of the distance across the refined auto-DRRAFTER models.
<sup>2</sup> The angle is the angle between the SL5-stem:SL5c coaxial stack and SL5a:SL5b coaxial stack. The parallel configuration is defined as 0° and the anti-parallel as 180°. The positive rotation is defined as rotating the SL5a:SL5b stack, positioned in the background, clockwise as in the orientations in **Figure 6**. See **Supplemental Figure S5** for a diagram explaining the angles. Error is the standard deviation of the angle across the refined auto-DRRAFTER models.
<sup>3</sup> Cryo-EM maps were not modeled, so distance and angle were estimated from the map alone.
<sup>4</sup> Only conformation 1 is considered, as conformation 2 does not have the listed coaxial stacks.

The auto-DRRAFTER models were further refined using ERRASER2 (version 2, available in Rosetta 3.10) (41) and evaluated for physical outliers and model-to-map fit (**Supplemental Table S5**). The non-base-paired regions of the RNA converged the least during modeling, reflecting uncertainty in their tertiary structure. Such heterogeneity is supported by low map resolvability and Q-score (42) in these regions of the map (**Supplemental Figure S4**). The 4-way junction is well converged and also has atomic resolvability above what is expected at 4.7 Å resolution (0.41 Q-score for junction atoms on average, with an expected 0.35 Q-score at that resolution). Refinement with ERRASER2 significantly improved the stereochemical quality of the models, while only marginally changing model-to-map fit scores, indicating ERRASER2 was able to successfully refine these models (**Supplemental Figure S6, Supplemental Table S5**). Additionally, after refinement with ERRASER2, the models remained divergent in the regions with poor map resolvability, showing that the set of models continues to reflect experimental uncertainty post-refinement (**Supplemental Figure S4, Supplemental Table S4**). These trends hold true for modeling of all constructs in this study (**Supplemental Figure S6**).

### Substantiation of the SARS-CoV-2 SL5 domain 3D structure

Next, we investigated an RNA segment, SL5-6 (residues 148-343, 196 nt, 63.1 kDa), which contains the full SARS-CoV-2 SL5 domain (residues 150-294) and its nearest neighboring domain, the SL6 domain (residues 302-343) (**Figure 2A**). M2-seq experiments revealed that the SL5 secondary structure folds independently of the SL6 domain with a 7 nt linker region (residues 295-301) (**Supplemental Figure S2**). In the cryo-EM map of SL5-6 (7.8 Å resolution, modeling convergence 4.4 Å **Supplemental Figures S4, S7**), we resolved SL5, which retains the previously observed T-shaped 3D fold (**Figure 1E**), but did not observe density corresponding to SL6 (**Figures 2B, E, Supplemental Figure S7**). To test the stem assignments in the SL5-6 structure and verify the absence of SL6, we designed extensions in the SL5b and SL5c stems (both 204 nt, 65.7 kDa) so that observed changes in the cryo-EM map would tag the corresponding stems, as previously done (12, 43–45). The tertiary structures of the extension constructs (SL5b extension: 7.4 Å resolution, modeling convergence: 2.7 Å; SL5c extension: 9.0 Å resolution, modeling convergence: 2.8 Å **Supplemental Figures S4, S7**) exhibited new densities in the extended stems, substantiating the stem assignments in our SL5-6 and SL5 models (**Figures 2C, D, F, G**). Though these maps were modeled successfully by auto-DRRAFTER, these models were not deposited to the PDB due to the partial resolvability of the construct; the maps only resolve part of the RNA construct, which may have led to the distortions at the apical ends of stems in the model (**Supplemental Figure S4**).

**Figure 2:**
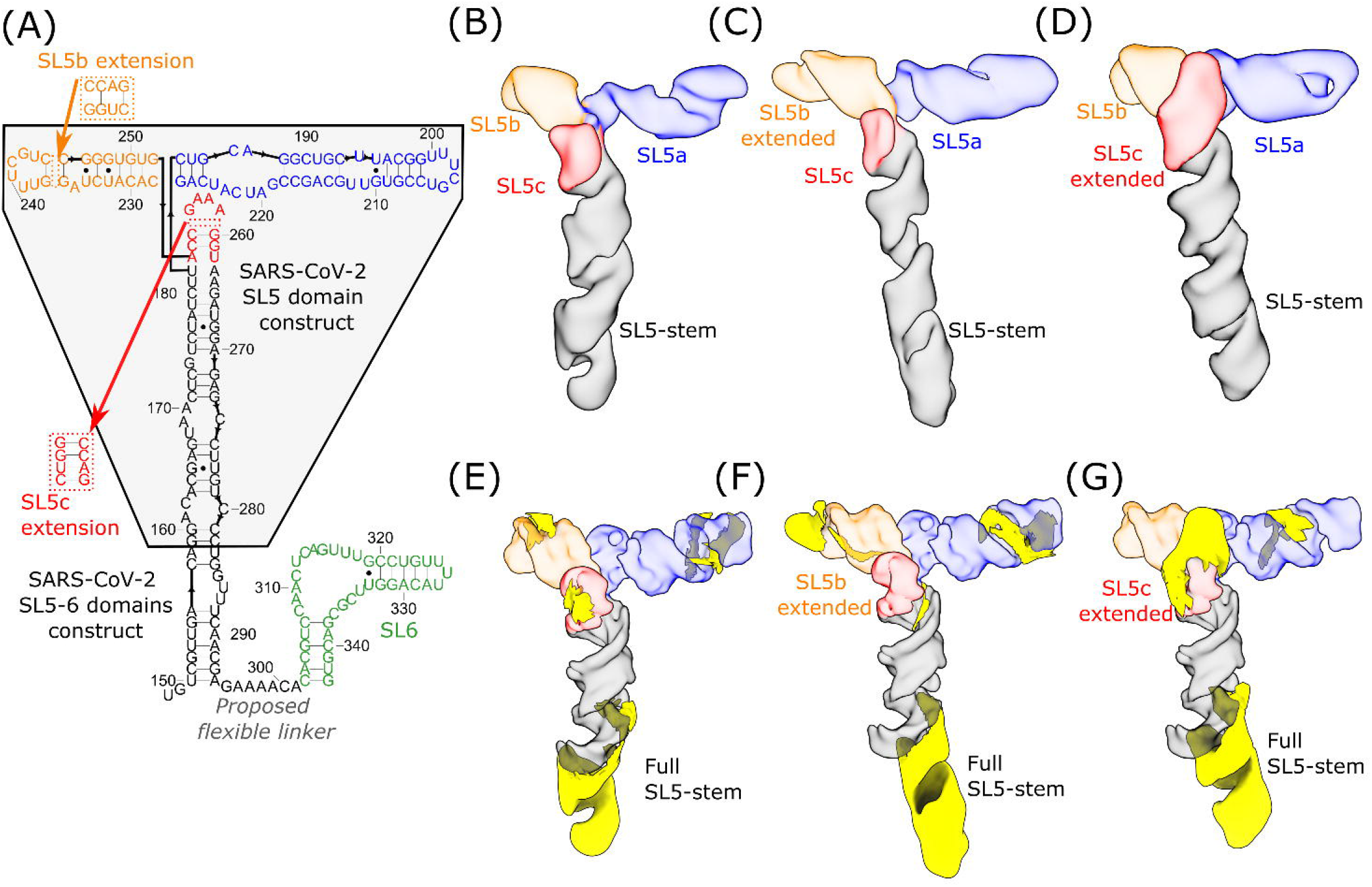
Substantiation of SARS-CoV-2 SL5 domain 3D structure using extension constructs. **(A)** The secondary structure, derived from M2-seq, of the SL5-6 domains of SARS-CoV-2 is depicted and colored in black (SL5-stem), blue (SL5a), orange (SL5b), red (SL5c), and green (SL6). The original construct used in this study for SARS-CoV-2 SL5 (Figure 1) is highlighted in the gray box. Relative to this construct, all SL5-6 constructs are extended to include the full SL5-stem and an additional stem-loop, SL6. In addition, the location of the four base-pair extensions to SL5b and SL5c are depicted. The cryo-EM maps of **(B)** SL5-6, **(C)** SL5-6 with SL5b extended, and **(D)** SL5-6 with SL5c extended are displayed, colored, and labeled by stem. Extensions are highlighted in yellow after masking out the density of the original SL5 construct for **(E)** SL5-6, **(F)** SL5-6 with SL5b extended, and **(G)** SL5-6 with SL5c extended.

We hypothesized that SL6 was not well resolved due to flexibility in the natural linker sequence connecting SL5 and SL6. This hypothesis is supported by an additional density consistent with SL6 appearing adjacent to the SL5-stem when SL5a, SL5b, and SL5c are removed, either by subtracting their density from particle images or from imaging an RNA construct (129 nt, 41.4 kDa) without these stems (**Supplemental Figure S8**). Furthermore, unlike in the map containing the full SL5, in the map containing the SL5-stem with SL6, the helical grooves of the SL5-stem are not resolved, consistent with averaging that would result from SL6 moving around the SL5-stem (**Supplemental Figure S8**). The natural extension of SL5 by SL6, along with the designed extensions, preserve SL5′s distinct T-shaped fold, which suggests that the tertiary structures of SL5 and SL6 fold independently of each other.

### 3D structure of the SL5 domain in betacoronaviruses

Based on our cryo-EM analysis, the SL5 domain of SARS-CoV-2 has a defined tertiary fold, as opposed to a highly flexible ensemble. The coronavirus genome may have evolved this sequence to have a specific arrangement of SL5′s four stems to serve a functional role. However, this arrangement could also be a biophysical coincidence and not be a result of natural selection. While the secondary structure of the SL5 domain is conserved in most coronaviruses, it is unknown whether its tertiary structure would also be conserved, given the region’s low sequence conservation (pairwise sequence identity mean of 54.3% and minimum of 46.9% for the six sequences studied here, **Supplemental Note S1**) and variation in stem lengths. 3D conservation of the arrangement of these stems across coronaviruses would further support the importance of this feature for SL5 function.

We therefore carried out cryo-EM to resolve the SL5 domain of other coronaviruses, with a focus on human-infecting coronaviruses, for structural comparison with the SL5 domain of SARS-CoV-2. Betacoronaviruses contain five human-infecting coronaviruses, SARS-CoV-2, SARS-CoV-1, MERS, HCoV-OC43, and HCoV-HKU1. Amongst these viruses, HCoV-OC43, and HCoV-HKU1 are members of the *Embecovirus* subgenus, which was previously found to have replaced UUYYGU hexaloops in SL5 with repetitive loop motifs elsewhere in the genome (27), and so our studies focused on SARS-CoV-1 and MERS.

First, we examined the SL5 domain from SARS-CoV-1 (residues 151-291, 143 nt, 46.1 kDa), which has high sequence similarity with SARS-CoV-2 (85.9% sequence identity, **Supplemental Note S1**) and also belongs to the *Sarbecovirus* subgenus. We found that SL5 of SARS-CoV-1 (7.0 Å resolution, 2.6 Å modeling convergence, **Supplemental Figures S4, S9**) adopts the same T-shaped fold and junction geometry as that of SARS-CoV-2, with an inter-helical angle of 86.8±0.5° and distance between UUYYGU hexaloops of 82±1 Å (N=20, **Figure 3B**, **Table 1**, **Supplemental Figure S4**).

**Figure 3:**
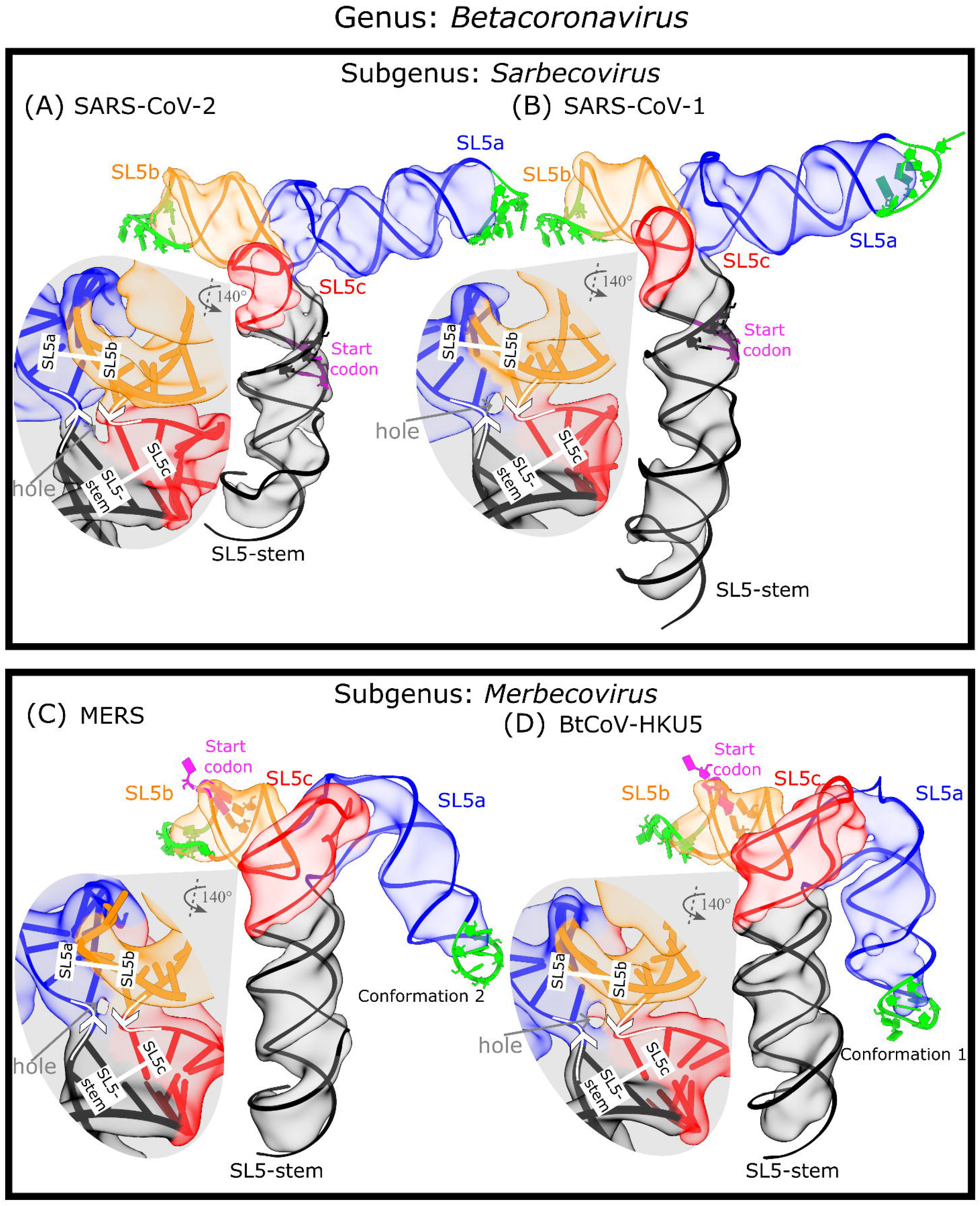
The 3D global fold of SL5 in betacoronaviruses. The cryo-EM maps and representative model of the SL5 domain of two sarbecoviruses, **(A)** SARS-CoV-2 and **(B)** SARS-CoV-1, and two merbecoviruses, **(C)** MERS and **(D)** BtCoV-HKU5, are colored by and labeled by stem. The nucleotides in the start codon are sequestered in a stem, and these base-paired nucleotides are displayed with the start codon in magenta. The nucleotides in the UUYYGU hexaloops are displayed in lime green. For each map, a zoom-in of the four-way junction cryo-EM map and model is displayed, showing the 5′-to-3′ direction with white arrows, the coaxial stacking with white bars, and the junction hole with a gray arrow.

Expanding the cryo-EM analysis to more distant coronavirus relatives, we examined the orthologous SL5 domain from MERS (residues 206-338, 135 nt, 43.7 kDa), which belongs to the different *Merbecovirus* subgenus of betacoronaviruses. From the secondary structures obtained from “large-library” M2-seq and the literature (6), we already noted a difference: while the UUYYGU hexaloops are still found on SL5a and SL5b, the SL5c stem from MERS is significantly longer than SL5c from sarbecoviruses (**Supplemental Figure S2**). Interestingly, our MERS SL5 cryo-EM analysis showed three conformation (6.9, 6.4, 6.4 Å resolution, 3.5, 3.2, 3.4 Å modeling convergence, **Supplemental Figures S9-S10**) and had the same conformation seen in sarbecovirus orthologs: helical stacking, junction geometry, and inter-helical angle matching within experimental error (**Figure 3C**, **Table 1**, **Supplemental Figure S10).**

To investigate the conservation of the SL5 fold further, we examined an additional merbecovirus ortholog, the SL5 domain of BtCoV-HKU5 (residues 188-320, 135 nt, 43.7 kDa). BtCoV-HKU5 SL5 has a 77.0% sequence identity to MERS SL5 (**Supplemental Note S1**) and “large-library” M2-seq and the literature (6) show the secondary structure is very similar to MERS SL5 (**Supplemental Figure S2**). The cryo-EM structure analysis of BtCoV-HKU5 SL5 resolved in four conformations (5.9, 6.4, 8.0, 7.3 Å resolution, 3.0, 3.0, 5.2, 3.0 Å modeling convergence, **Supplemental Figures S10-S11**) and again shares the same helical stacking and junction geometry with the other betacoronaviruses studied, with an inter-helical angle of 86±2° (**Figure 3D**, **Table 1**, **Supplemental Figure S5**). While auto-DRRAFTER was able to model all maps consistently, we did not deposit the coordinates for conformation 3 in the PDB because the helical grooves were insufficiently resolved (**Supplemental Figure S10**). As with the other three betacoronavirus domains imaged, the four-way junction is well resolved in the BtCoV-HKU5 maps, revealing a clear hole between the helical strands that separates perpendicular, coaxially stacked stems (**Figure 3D**).

While the four-way junction geometry is conserved between the sarbecovirus and merbecovirus orthologs, the merbecoviruses have a distinct ensemble of 3D folds that is conserved between MERS SL5 and BtCoV-HKU5 SL5. In particular, a flexible bend is observed emanating from the SL5a internal loop that allows the SL5a arm to swing in a hinge-like motion (**Figures 4A-B**). In order to model this conformational heterogeneity, the particles were classified into discrete classes to resolve cryo-EM maps that could individually be modeled (**Figures 4C-D**). Among these conformations, the junction geometry was conserved, with inter-helical angles occupying a narrow range of 81-84° and 84-88° for MERS and BtCoV-HKU5 SL5, respectively (**Supplemental Figure S5**). Despite the junction geometry conservation, the relative locations of the hands of the SL5a and SL5b arms are variable, increasing the range of distance between UUYYGU hexaloops to 74-85 Å and 71-85 Å for MERS and BtCoV-HKU5 SL5, respectively (**Supplemental Figure S5**).

**Figure 4:**
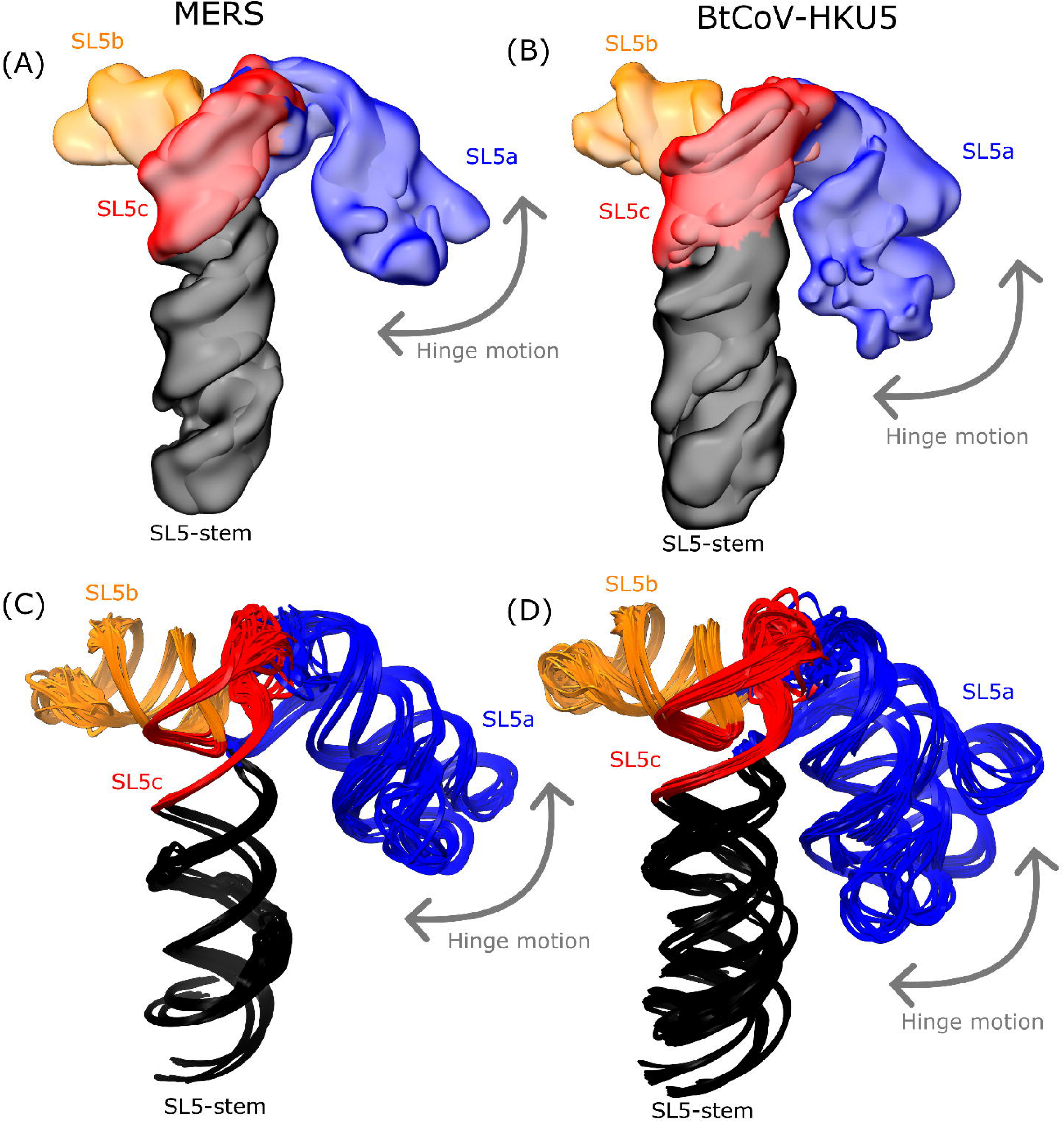
The hinge motion in the SL5a stem of merbecoviruses. The cryo-EM maps obtained after discrete classification of particles are overlaid for the **(A)** MERS and **(B)** BtCoV-HKU5 SL5 domains. The models derived from these maps using auto-DRRAFTER and ERRASER2 are overlaid for the **(C)** MERS and **(D)** BtCoV-HKU5 SL5 domains. All constructs are colored by stem, and the hinge motion is labeled with an arrow.

In addition, the merbecovirus orthologs display an unexpected tertiary interaction between the SL5a internal loop and SL5c apical loop (**Figures 5C, F**), whereas the sarbecovirus SL5c stem is too short to form this tertiary interaction (**Figure 5A**). Due to resolution limitations, the SL5a-SL5c interaction of merbecoviruses cannot be modeled with atomic precision. This uncertainty in the SL5a-SL5c interaction is reflected in the ensemble of models produced by auto-DRRAFTER (**Figures 5D, G**). The models, across all conformations and both merbecovirus orthologs, do converge in identifying the same interacting regions, namely the the asymmetric internal loop of SL5a, 5′-AAUU-3′ and the apical loop of SL5c, 5′-AAGGUGC-3′ (MERS: residues 264-267 and 397-313, respectively; BtCoV-HKU5: residues 246-249 and 289-295, respectively **Figures 5B, E**).

**Figure 5:**
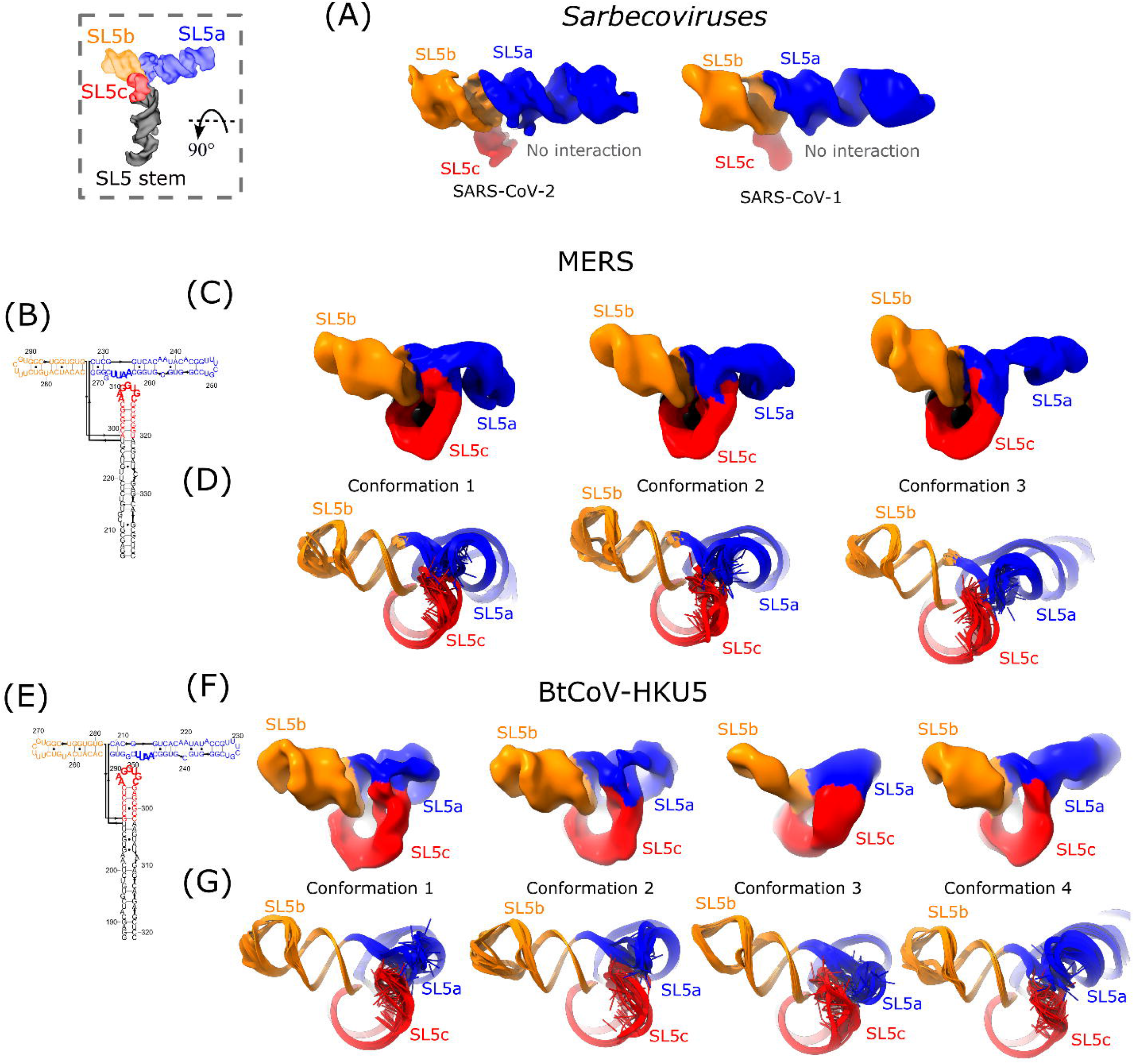
The tertiary interaction in the SL5 domains of merbecoviruses. The proposed tertiary interaction is displayed by viewing the top of the molecules with the SL5-stem at the back, relative rotation is shown in the top left of the figure. **(A)** The SL5 domains from members of the *Sarbecovirus* subgenus do not have densities connecting SL5c (red) and SL5a (blue), indicating there is no tertiary interaction between these stem-loops. In SL5 domains from the *Merbecovirus* subgenus, **(C)** MERS and **(F)** BtCoV-HKU5, all cryo-EM maps are displayed and colored by stem, showing that a density connecting the apical loop of SL5c to the first internal loop of SL5a was resolved. The models derived from the cryo-EM map using auto-DRRAFTER and ERRASER2 are displayed for the SL5 domains of **(D)** MERS and **(G)** BtCoV-HKU5. The cryo-EM maps were insufficiently resolved for modeling to converge on the atomic level details of this junction, but the same set of residues are consistently interacting. These interacting nucleotides are bolded in the secondary structures of **(B)** MERS and **(E)** BtCoV-HKU5.

### 3D structural comparison of the SL5 domain across alpha- and betacoronaviruses

We next looked to alphacoronaviruses to explore the 3D structure of SL5 in a different genus. We selected the SL5 domains from the remaining two human-infecting coronaviruses, HCoV-229E (residues 153-292, 140 nt, 45.1 kDa) and HCoV-NL63 (residues 138-295, 160 nt, 51.4 kDa) from the *Duvinocovirus* and *Setracovirus* subgenera, respectively, for investigation. While an experimental secondary structure for the SL5 domain in HCoV-NL63 was previously identified (6), the secondary structure for HCoV-229E was deduced by exhaustively modeling a set of published (26, 27), predicted (40, 46), and manually curated secondary structures into the cryo-EM map (**Supplemental Figure S12**). Only one secondary structure for HCoV-229E resulted in converged auto-DRRAFTER modeling that agreed with the cryo-EM map (**Supplemental Figure S13**). Beyond containing four helical stems, the secondary structures of these alphacoronaviruses differ from those of the previously examined betacoronaviruses. These alphacoronaviruses have three UUYYGU hexaloops, as opposed to two, and the four-way junctions contain unpaired nucleotides. Hence, we sought to investigate which 3D structural features, if any, were conserved within human-infecting alphacoronaviruses and between alpha- and betacoronaviruses.

HCoV-229E SL5 (6.5 Å resolution, 2.3 Å modeling convergence, **Supplemental Figures S9, S13**) and HCoV-NL63 SL5 (8.0, 8.4 Å resolution, not modeled, **Supplemental Figure S14**) form X-shaped folds. Both alphacoronavirus SL5s adopt the same helical stacking as the betacoronavirus domains, with the SL5-stem:SL5c and SL5a:SL5b coaxially stacking (**Figure 6**). The junction is well resolved in HCoV-229E, with a visible hole separating the stems (**Supplemental Figure S13**). For HCoV-NL63, however, the cryo-EM data was classified into two maps with distinct conformations (**Supplemental Figures S13-S14**). These maps did not achieve sufficient resolution to view the major or minor grooves of helices and thus, coordinates were not modeled into the maps. Nevertheless, the disparate lengths of each stem in HCoV-NL63 SL5 allow for unambiguous stem assignment to assess junction geometry (**Supplemental Figure S13**). One conformation reveals a distinct stacking pattern at the junction, in which the SL5-stem stacks coaxially with SL5a while SL5b stacks with SL5c. The other conformation is homologous with the HCoV-229E global fold that coaxially stacks SL5a and SL5b, as seen with all the other coronaviruses studied here.

**Figure 6:**
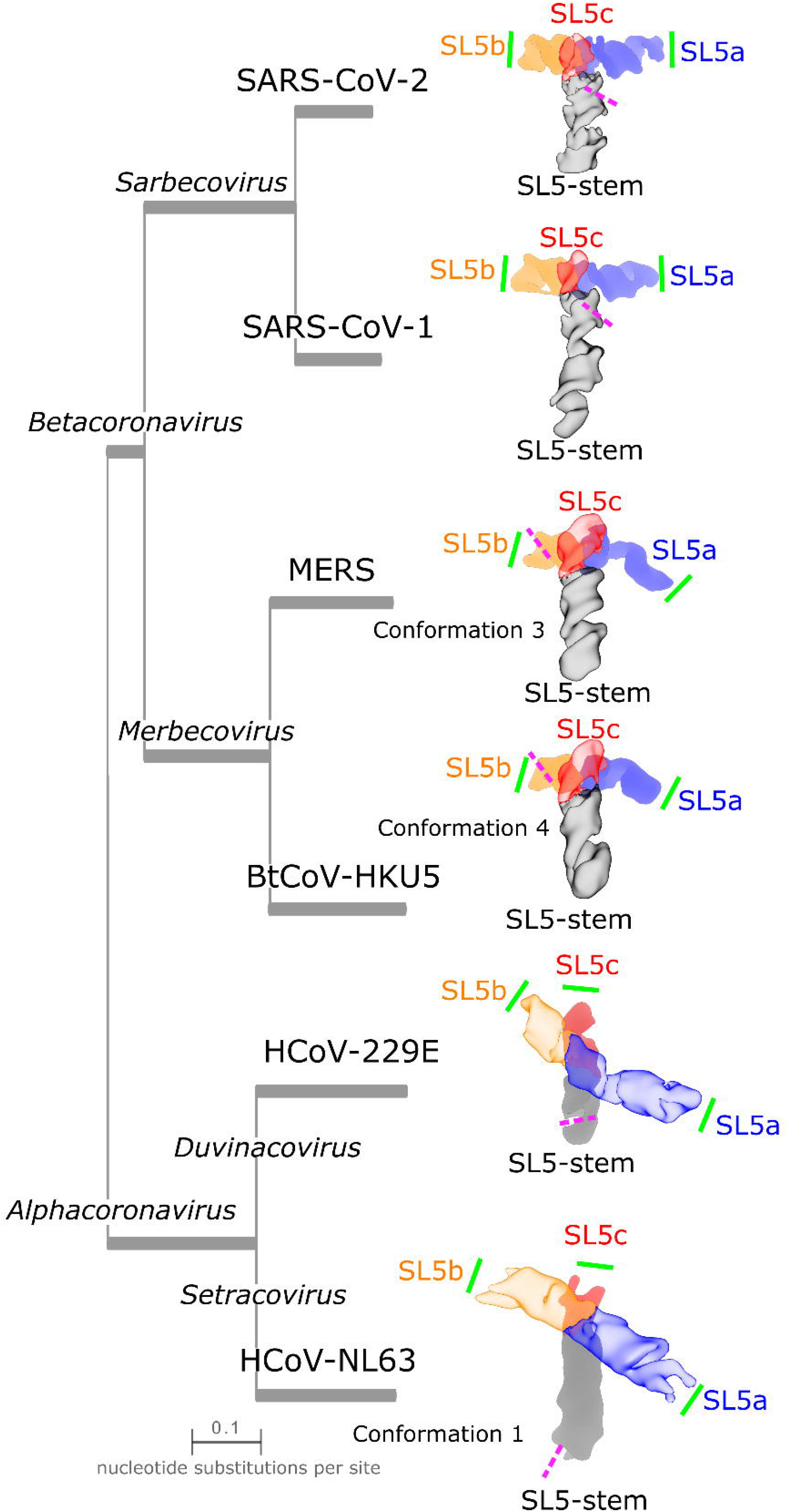
Similarities and differences in the tertiary folds of SL5 in alpha- and betacoronaviruses. To compare junction geometries between the SL5 domain of betacoronaviruses and alphacoronaviruses, the cryo-EM maps of the SL5 domains are displayed with the junction perpendicular to the text and with the stem pointed downwards as a reference. Additionally, SL5b is pointed left and SL5a right. The maps are colored by domain with the foreground, SL5-stem:SL5c for betacornoviruses SL5ba:SL5b for alphacoronaviruses, made transparent to enable the view of the stems below. The various species are positioned on a phylogenetic tree with branch length proportional to evolutionary distance. For BtCoV-HKU5, MERS, and HCoV-NL63 SL5, the conformation that is closest to other SL5 domains is displayed. The pink dashed lines indicate the position of the start codon; note that for HCoV-NL63, the SL5 construct imaged was truncated on the 3′ end directly before the start codon. The green solid line indicates the position of a UUYYGU hexaloop.

The inter-helical angle is similar among the alphacoronavirus SL5s, -121.3±0.2° (N=10) and -120° (estimated) for HCoV-229E and HCoV-NL63 respectively, despite being quite different in sequence (59.1% sequence identity, **Supplemental Note S1,** **Figure 6**, **Table 1**). This inter-helical angle for X-shaped alphacoronavirus SL5s is distinct from the near-perpendicular angle formed by the T-shaped betacoronaviruses SL5s (81-88°) (**Figure 6**). Despite this difference, the UUYYGU hexaloops are positioned a similar distance apart of 92 and 95 Å for HCoV-229E SL5 and HCoV-NL63 SL5 conformation 1, respectively (**Table 1**). Surprisingly, despite a different coaxial stacking pattern, HCoV-NL63 conformation 2 has a similar estimated SL5a:SL5b UUYYGU hexaloop distance of 90 Å. These distances are ∼10 Å longer than the distances, 82-84 Å, observed in the betacoronavirus SL5 domains. While not identical distances, the observed range is narrower than the ranges of distance predicted by *de novo* 3D structure prediction algorithms in CASP15 (predictions were for SARS-CoV-2 and BtCoV-HKU5 SL5) (29) (**Supplemental Figure S5**).

## Discussion

We have presented structural characterization of the SL5 domain across six coronaviruses. This study was enabled by the increasing throughput of 3D RNA structural characterization, made possible by single particle cryo-EM integrated with biochemical secondary structure mapping, automated computer modeling and structure validation (38, 42). Cryo-EM offers the opportunity to increase the knowledge base of RNA 3D global folds, particularly through the ability to study RNA homologs, as carried out here, revealing similarities and differences that may be relevant to function. While cryo-EM of these small RNA samples (40.0-65.7 kDa in size) was limited in resolution, for all but one sample, we were able to achieve sufficient resolution to resolve major and minor grooves and even resolve the hole in the four-way junction. This resolution enabled the unambiguous identification of stem positions and hence the RNA’s global fold (**Figures 1, 3, 4, 6**). This demonstrates the utility of cryo-EM to resolve the 3D folds of some viral RNA elements that may exhibit flexibility (**Figure 5**). We further tested our structural models, demonstrating that the SL5 domain folds independently of the closest downstream stem-loop, SL6, and confirmed our stem placements using extension constructs (**Figure 2**). Although atomic level detail cannot be ascertained from the structures, we have deposited multiple models in the Protein DataBank to characterize this uncertainty estimation.

The SL5 domains of all human-infecting coronaviruses that contain the UUYYGU hexaloops, along with the bat-infecting BtCoV-HKU5 SL5, fold into a limited number of stable conformations, which we have resolved using cryo-EM. We found that the SL5 domain of SARS-CoV-2 folds into a T-shaped structure, modeled using a 4.7 Å cryo-EM map, in which the SL5-stem and SL5c form a continuous, long, coaxial stack that lies perpendicular to a second continuous, long, coaxial stack formed by SL5a and SL5b. This stacking pattern is conserved across all of the imaged SL5 domains while the junction geometry and inter-helical angles of 81-88° for betacoronaviruses and approximately -120° for alphacoronaviruses are conserved within each genus. Although the junction angle is genus-specific, all coronaviruses studied display an experimentally resolved conformation that places a pair of UUYYGU hexaloops a distance of 82-92 Å apart at opposing ends of an SL5a:SL5b coaxial stack.

The *Merbecovirus* subgenus of betacoronaviruses has an additional subgenus-specific structural feature: an interaction between the SL5a internal loop and SL5c apical loop (**Figure 5**). Signatures for this interaction have not been observed in chemical mapping studies, which are frequently not sensitive to tertiary interactions (47, 48). This interaction could help stabilize the SL5 stacking pattern and junction orientation, relative to other global stacking patterns and inter-helical rotations that would position the SL5a internal loop and SL5c apical loop apart.

The structurally conserved features of SL5 across coronavirus genera suggest potential functional roles for SL5. First, the SL5 domain sequesters the start codon in a stem, but this sequence must be exposed by unfolding SL5 to initiate translation. Thus, the SL5 element may act as a switch, enforcing exclusivity between viral translation and an as yet unknown function corresponding to SL5′s folded structure, such as viral replication or viral packaging, that should not occur at the same time as viral translation. Work on the structure of SL5 bound to translation initiation machinery may elucidate the nature of the conformational change required for translation.

Second, SL5 contains two or three UUYYGU hexaloops in the selected betacoronavirus and alphacoronavirus domains, respectively. These cryo-EM structures reveal that the distances between hexaloops are within a narrow range, 82-92 Å, and are placed at opposing ends of a long, continuous coaxial stack. This conservation suggests SL5 may position the hexaloops for a functional reason. As one possibility, the viral genome must be selectively packaged, compared to host RNA, in virions. While UUYYGU motifs will recur throughout host RNAs by chance, the stereotyped placement of two such sequences as apical loops on opposing sides of a coaxial stack is less likely to occur in host RNAs. The two loops could therefore be selectively recognized by oligomers of viral or host proteins with RNA-binding domains (49). While, in principle, structures other than four-way junctions can produce similar coaxial stacks, the SL5 four-way junction provides a natural solution. For example, three-way junctions can provide similar positionings of UUYYGU motifs at opposite ends of a co-axial stack, but they can form a larger number of alternative stacking patterns than four-way junctions, which would give rise to competing, potentially non-functional geometries (50).

In addition to resolving dominant structures of the SL5 elements, since we imaged RNA in vitreous ice, we were able to resolve a “cryo-ensemble” of structures. We observed a flexible hinge of SL5a in both merbecovirus SL5 domains and an alternative stacking pattern at the four-way junction of the HCoV-NL63 SL5 domain. Additionally, we observed indications of other modes of flexibility that we did not model — for example, the SL5-stem in SARS-CoV-1 (**Supplemental Figure S9**). Nevertheless, we observed some limitations in current cryo-EM data analysis procedures. For example, we hypothesize that the 7 nt flexible linker between SL5 and SL6 left SL6 (13.3 kDa) unresolved. Also heterogeneity likely limited the angular assignment accuracy and resolution of cryo-EM maps, particularly in the case of the HCoV-NL63 SL5. These shortcomings highlight the need for further methods to be developed and tested on RNA constructs, which may reveal additional, unobserved heterogeneity, especially for small, helical structures with continuous hinge motions. Additionally, while imaging RNA in vitreous ice is a step towards achieving more near-native conditions, the effects of excising these RNA elements from genomic and cellular contexts, as well as the effects of the grid environment and vitrification on RNA structure and heterogeneity are unknown. Complementary, lower resolution experimental techniques such as single molecule FRET or solution X-ray scattering paired with molecular dynamics, could be used to further understand biases of the different methods and more quantitatively assess the relative populations of RNA species in solution (51, 52).

Despite these caveats, the ability to solve the “cryo-ensemble” for such small RNA molecules, although likely not representing the full, biologically relevant ensemble, was important for understanding the conservation of the 3D fold in the coronaviruses’ SL5 domain. It is possible that the crystallized structures of these RNAs would not have revealed the same conservation as evident when we analyze the “cryo-ensemble” – for example, the SL5 domain from HCoV-NL63 may crystallize as conformation 2, the conformation with alternative base-stacking. An “ensemble-view” of RNA molecules enhances structure-function interpretations, and may be more readily brought to bear in RNA systems in the future through cryo-EM (53).

Finally, the analysis of conserved structural features of the SL5 domain suggests strategies for the structure-guided design of pan-coronavirus therapeutics. In particular, there may be druggable pockets at the four-way junction, conserved among the betacoronaviruses studied here, or at the SL5a-SL5c tertiary interaction in MERS, the human-infecting coronavirus with the highest fatality rate. These pockets could be the targets for small molecules such as ribonuclease-targeting chimeras (RIBOTACs) (20). Alternatively, targeting two regions could improve the specificity of a therapeutic. For example, antisense oligonucleotides that target both the start codon and the four-way junction region of SL5 would serve the dual purpose of slowing viral translation while also preventing formation of the SL5 tertiary structure, which appears important for a distinct viral function. Different classes of therapeutics could also take advantage of the stereotyped positioning of the conserved UUYYGU hexaloops. For example, circularized or chemically modified RNAs could present UUYYGU hexaloops positioned 82-92 Å apart and thereby compete for the binding of proteins with the viral genomic RNA. Additionally, the catalog of SL5 structures may enable faster response for emerging threats by enabling the design of therapeutics against orthologous structures from any of the viruses resolved herein, which represent most known human-infecting coronaviruses. Finally, the models of the “cryo-ensemble” could present an opportunity for structure-guided drug design, enabling the targeting of one conformation over the others with the aim of trapping the RNA in a non-functional conformation. These efforts will likely require the complementary contributions from X-ray crystallography and NMR to resolve higher resolution details important for designing and refining structure-guided small molecule therapeutics.

## Data Availability

The data supporting the findings of this manuscript are available from the corresponding authors upon reasonable request. The reactivity traces can be found on the RNA Mapping DataBase (RMDB). The “scarless” M2-seq SL5 library reactivity files have the following RMDB IDs: COVSL5_DMS_0001, COVSL5_DMS_0002, COVSL5_NOM_0001, and COVSL5_NOM_0002. The “large-library” reactivity RDAT files have the following RMDB IDs: SL5HKU_DMS_0001, SL5HKU_2A3_0001, SL5HKU_NOM_0001, SL5HKU_NOM_0002, SL5MER_DMS_0001, SL5MER_2A3_0001, SL5MER_NOM_0001, SL5MER_NOM_0002, SL5CV2_DMS_0001, SL5CV2_2A3_0001, SL5CV2_NOM_0001, and SL5CV2_NOM_0002. The sequencing data can be found on the NIH Sequence Read Archive with the BioProject accession number: PRJNA1039878. The “scarless” M2-seq SL5 library FASTQ files have the following SRA accession numbers: SRR26810683, SRR26810682, SRR26810681, SRR26810680; the “large-library” combined FASTQ files have the SRA accession number: SRR26827601. The cryo-EM maps are deposited in the Electron Microscopy Data Bank (EMDB) under the following accession codes: SARS-CoV-2 SL5: EMD-42818, SARS-CoV-2 SL5-6: EMD-42821, SARS-CoV-2 SL5-6 with SL5b extended: EMD-42820, SARS-CoV-2 SL5-6 with SL5c extended: EMD-42819, SARS-CoV-1 SL5: EMD-42816, MERS SL5 conformation 1: EMD-42809, MERS SL5 conformation 2: EMD-42810, MERS SL5 conformation 3: EMD-42811, BtCoV-HKU5 SL5 conformation 1: EMD-42801, BtCoV-HKU5 SL5 conformation 2: EMD-42805, BtCoV-HKU5 SL5 conformation 3: EMD-42802, BtCoV-HKU5 SL5 conformation 4: EMD-42808, HCoV-229E SL5: EMD-42803, HCoV-NL63 SL5 conformation 1: EMD-42813, and HCoV-NL63 SL5 conformation 2: EMD-42814. The atomic models are deposited in the Protein Data Bank (PDB) under the following accession codes: SARS-CoV-2 SL5: 8UYS, SARS-CoV-1 SL5: 8UYP, MERS SL5 conformation 1: 8UYK, MERS SL5 conformation 2: 8UYL, MERS SL5 conformation 3: 8UYM, BtCoV-HKU5 SL5 conformation 1: 8UYE, BtCoV-HKU5 SL5 conformation 2: 8UYG, BtCoV-HKU5 SL5 conformation 4: 8UYJ. Models were not deposited in the PDB for SARS-CoV-2 SL5-6, SARS-CoV-2 SL5-6 with SL5b extended, SARS-CoV-2 SL5-6 with SL5c extended, BtCoV-HKU5 SL5 conformation 3, and HCoV-229E SL5 and can instead be found in the accompanying GitHub repository (https://github.com/DasLab/Coronavirus_SL5_3D). All raw movies and particle stacks are being uploaded to the Electron Microscopy Public Image Archive (EMPIAR).

## Author contributions

R.C.K. and I.N.Z. performed and analyzed scarless M2-seq experiments. R.C.K. designed the library for library-based chemical mapping and R.H. performed the chemical mapping. I.N.Z., K.Z., and S.L. conducted preliminary work optimizing the SARS-CoV-2 SL5 RNA for cryo-EM with I.N.Z preparing the SL5-6 and first SL5 RNA and K.Z. and S.L. preparing cryo-EM grids and performing imaging. R.C.K. and L.X. chose SL5 domains and designed extension constructs. R.C.K. and L.X. prepared the remainder of RNA samples and R.C.K, L.X., and X.Z. prepared cryo-EM grids and performed imaging. R.C.K. processed the cryo-EM data and built and validated the models. G.P.N. compared the validation metrics before and after refinement. R.C.K., L.X., K.Z., I.N.Z., R.D., and W.C., conceived of the study, and wrote and edited the manuscript.

## Acknowledgements

We thank Paul Berg, Jeffrey S. Glenn, and members of the Glenn, Barna, and Puglisi labs (Stanford) for helpful discussions on the nature and structure of SL5. We thank Stanford University, Stanford Research Computing Center, SLAC Shared Scientific Data Facility, the Stanford-SLAC Cryo-EM Center, and the Stanford Biochemistry administrative staff for their computational resources and support that contributed to this research. This work was supported by: Stanford BioX (Bowes Graduate Student Fellowship to R.C.K., Interdisciplinary Initiative Program to R.D.), the National Institutes of Health (R35 GM122579 to R.D.), Howard Hughes Medical Institute (to R.D.), the National Science Foundation (GRFP DGE-1656518 to L.X.), the National Institutes of Health Common Fund Transformative High-Resolution Cryo-Electron Microscopy program (U24 GM129541 to W.C.) for cryo-EM data collection, National Key R&D Program of China (2022YFC2303700 to K.Z. and S.L., and 2022YFA1302700 to K.Z.). This project was initiated during the COVID-19 pandemic with support from a Stanford University ChEM-H COVID-19 Drug and Vaccine Prototyping seed grant (P. Berg, J. S. Glenn, R.D., W.C.). This article is subject to HHMI’s Open Access to Publications policy. HHMI lab heads have previously granted a nonexclusive CC BY 4.0 license to the public and a sublicensable license to HHMI in their research articles. Pursuant to those licenses, the author-accepted manuscript of this article can be made freely available under a CC BY 4.0 license immediately upon publication.

## Materials and Methods

“Scarless” two-dimensional chemical mapping

2D chemical mapping was performed using an optimized M2-seq pipeline (39), which uses mutational sequencing-based inference of dimethyl sulfide (DMS) modifications on a library of folded RNA that contains purposeful random sequence variations to better infer stems. The scarless protocol was modified to remove primer binding sequences from the RNA to prevent unwanted secondary structure interference during DMS modification. This modification was achieved by appending removable primer sequences to the 3′ end of the DNA encoding the region of interest, which is used for error-prone PCR (epPCR) and cleaved off prior to *in vitro* transcription. After DMS modification, sequencing libraries were made using two ligation steps on ssRNA and ssDNA followed by a primer-biased PCR.

The same segments of the NC.045512.2 reference genome used for cryo-EM of the SARS-CoV-2 SL5 domain (residues 159-282) and SL5-6 domains (residues 148-343) were used for M2-seq. The SL5 domain was prepended with a single-point mutated Φ6.5 T7 RNA polymerase promoter in order to remove the MlyI recognition site and the SL5-6 domains were prepended with a Φ2.5 T7 RNA polymerase promoter. The SL5 domain was appended with a 20 bp region that had a MlyI recognition site such that this appended region could be cut-off with a blunt edge by MlyI. The SL5-6 domains were appended with a 20 bp region that had a BsaI recognition site such that a five nucleotide overhang was left that was later digested. Primers to assemble the sequences were designed for PCR assembly using Primerize (listed in **Supplemental Table S6**) (54), ordered from Integrated DNA Technologies, assembled into full-length double-stranded DNA by PCR assembly following the Primerize protocol using Phusion polymerase (in-house), ′High-Fidelity′ buffer (Thermo Scientific #F-530), and an annealing temperature of 64°C, and purified QIAquick PCR Purification Kit (QIAGEN, #28104). epPCR was conducted with 10 mM Tris (pH 8.3), 50 mM KCl, 0.5 mM MnCl_2_, 5 mM MgCl_2_, 1 mM dTTP, 1 mM dCTP, 0.2 mM dATP, 0.2 mM dGTP, 2 µM forward primer, and 2 µM reverse primer (listed in **Supplemental Table S6**), 160 ng dsDNA template, and Taq polymerase (New England Biolabs #M0495), using an annealing temperature of 49°C. The DNA of the SL5 domain was then digested using MlyI (New England Biolabs #R0610) and the DNA of the SL5-6 domains was then digested using BsaI (New England Biolabs #R0535) followed by Mung Bean Nuclease (New England Biolabs #M0250), all in CutSmart Buffer (New England Biolabs #B6004). Product homogeneity was assessed by 1x TBE - 4% agarose gel electrophoresis and visualized with SYBR Safe after PCR assembly, epPCR, and restriction enzyme digest.

RNA was synthesized by *in vitro* transcription (TranscriptAid T7 High Yield Transcription Kit, Thermo Scientific #K0441), then purified by column purification (RNA Clean & Concentrator, Zymo Research #R1017). After denaturing the RNA for 3 minutes at 90°C followed by 10 minutes at room temperature, RNA was refolded at 50°C for 20 minutes, followed by 3 minutes at room temperature in a buffer containing 300 mM sodium cacodylate (pH 7.0) and 10 mM MgCl_2_.

Samples were incubated at 37°C for 6 minutes with 1.5% DMS, unmodified samples were also incubated at this temperature without DMS. The reactions were quenched with β-mercaptoethanol (50% by volume) followed by column purification (Oligo Clean & Concentrator, Zymo Research #D4060, here and below). The adenylated 18 nt unique molecular indicator (UMI) linker (listed in **Supplemental Table S6**) was ligated to the 3′ end using T4 RNA Ligase 2 truncated KQ (New England Biolabs #M0373, 400 units per sample) in T4 Ligase reaction buffer (New England Biolabs #B0216) and PEG 8000 (33% w/v) at 25°C for 1.5 hours followed by a column purification. The residual linker was deadenylated using a 5ʹ deadenylase (New England Biolabs #M0331, 50 units per sample) in NEBuffer 1 (New England Biolabs #B7030) at 30°C for 1 hour followed by column purification. The residual DNA was then digested using RecJf (New England Biolabs #M0264, 30 units per sample) in NEBuffer 2 (New England Biolabs #B6002) at 37°C for 30 minutes followed by column purification. The RNA was then reverse transcribed using TGIRT-III (InGex, 100 units) in 50 mM Tris-HCL (pH 8.0), 3 mM Mg Cl_2_, 75 mM KCl, 3 mM DDT (Invitrogen), 1.5 µM of primers listed in **Supplemental Table S6**, ramping the temperature from 60°C to 75°C at a rate of 0.1°C/s and holding for 15 minutes prior to adding the dNTPs (1.25 mM), after which the solution was held at 64°C for 3 hours. The reaction was quenched (122 mM NaCl, 49 mM HCl, 110 mM Na-acetate) neutralizing with 78 mM NaOH at 90°C for 3 minutes followed by column purification. The 3′ ligation oligo (listed in **Supplemental Table S6**) was ligated onto the cDNA using CircLigase I (Lucigen #CL4111K, 100 units) with CircLigase Reaction Buffer, PEG 8000 (16.25% w/v), 2.5 mM MnCl_2_, 50 µM rATP, and 2.5 µM 60°C overnight, then for 10 minutes at 80°C followed by column purification. The cDNA was size selected using denatured polyacrylamide gel electrophoresis followed by dPAGE purification (small-RNA PAGE Recovery Kit, Zymo Research #R1070). The cDNA was then amplified using Q5-Ultra II DNA polymerase (New England Biolabs #M0544) using the primers listed in **Supplemental Table S6**. Product size, purity, and concentration were confirmed on a 1% agarose gel, Agilent Bioanalyzer 2100 Small RNA, and Qubit (Thermo Fisher). The DNA was pooled for sequencing on an Illumina MiSeq 2×300. Demultiplexed sequencing results were analyzed using the M2-seq pipeline (https://github.com/ribokit/M2seq) (39) and ShapeMapper (55) to create mutation strings for each read and then 2D mutational profiles through the script simple_to_rdat.py.

### “Large-library” two-dimensional chemical mapping

To accelerate mutate-and-map characterization of secondary structures of multiple orthologs, we explored the use of oligonucleotide libraries that encoded SL5 domains as well as all of their single mutants, with 3′ barcode hairpins to allow unambiguous deconvolution of the mutant profiles. The library sequences were prepared using custom scripts (https://github.com/DasLab/big_library_design), with 3′ barcode hairpins screened computationally. EternaFold was used to predict the secondary structures and to ensure that the barcode is predicted to fold into a hairpin and that the SL5 domain was not predicted to interact significantly with the barcode or flanking sequences. The library sequences are listed in **Supplemental Table S6**.

The oligonucleotide library (synthesized by Agilent) was amplified using emulsion PCR (per reaction oil phase: 12 μL of ABIL EM90 (Evonik), 0.15 μL of Triton X-100 (Sigma Aldrich #T8787), 287.85 μL of mineral oil (Sigma Aldrich #M5904); per reaction aqueous phase: 26.625 μL of DNase/RNase-free water, 3 μL of 100 μM “Eterna” forward primer, 3 μL of 100 μM “Tail2” reverse primer (listed in **Supplemental Table S6**), 3 μL of oligo pool template with concentration of 1 ng/μL, 1.875 μL of Bovine Serum Albumin (20 mg/mL, Thermo Fisher #B14), and 37.5 μL of 2X Phire Hot Start II PCR Master Mix (Thermo Fisher #F125L); with an annealing temperature of 55°C. The emulsion PCR product was purified using QIAquick PCR Purification Kit (QIAGEN, #28104), and the RNA library was synthesized by *in vitro* transcription (TranscriptAid T7 High Yield Transcription Kit, Thermo Scientific #K0441). RNA was refolded at 50°C for 30 minutes in a buffer containing 200 mM Bicine, 200 mM KOAc, and 13.3 mM MgCl_2_. One copy of the RNA library was modified by 3% DMS, and the other copy was modified by 100 mM 2A3 ((2-Aminopyridin-3-yl)(1*H*-imidazol-1-yl)methanone, TOCRIS #7376). The DMS modified RNA was reverse transcribed by MarathonRT Reverse Transcriptase (Kerafast #EYU007), and the 2A3 modified RNA was reverse transcribed by SuperScript II Reverse Transcriptase (Thermo Fisher #18064022) and primers listed in **Supplemental Table S6**. We used denatured polyacrylamide gel electrophoresis to size-select the cDNA, and the cDNA was amplified (PCR mixture: 1 μL of 100 μM “cDNAamp” forward primer, 1 μL of 100 μM “cDNAamp” reverse primer, 8 μL of water, 3 μL of cDNA template, and 12.5 μL of 2X Phire Hot Start II PCR Master Mix (Thermo Fisher #F125L); with an annealing temperature of 65°C) and pooled for Illumina NovaSeq X Plus next generation sequencing. We analyzed the sequencing data FASTQ files using the Ultraplex-Bowtie2-RNAFramework pipeline (https://github.com/DasLab/ubr), and generated RDAT files, which contain the RNA sequences and their respective reactivity profiles.

### Secondary structure modeling

Chemical mapping profiles acquired in the mutate-and-map experiments above were analyzed with Biers (https://ribokit.github.io/Biers/) to generate normalized 1D DMS profiles and 2D Z-scores. Biers was then used to create secondary structure predictions guided by the 1D DMS profiles and 2D Z-scores using ShapeKnots, with 100 bootstrapping iterations to estimate stem confidence values. The secondary structure with 1D DMS profile was depicted using RiboDraw (https://github.com/ribokit/RiboDraw/) (56). The raw data and Z-score plots were visualized using custom scripts. All scripts can be found in the accompanying GitHub repository (https://github.com/DasLab/Coronavirus_SL5_3D).

### Sample preparation for cryo-EM

Primers to assemble the sequences (sequence of interest with a T7 promoter) were designed for PCR assembly using Primerize (listed in **Supplemental Table S6**) (54), ordered from Integrated DNA Technologies, assembled into full-length double-stranded DNA by PCR assembly following the Primerize protocol using Phusion polymerase (in-house), and purified QIAquick PCR Purification Kit (QIAGEN, #28104). RNA was synthesized by *in vitro* transcription (TranscriptAid T7 High Yield Transcription Kit, Thermo Scientific #K0441), then purified by column purification (RNA Clean & Concentrator Kits, Zymo Research #R1017) and by denaturing PAGE gel extraction (ZR small-RNA PAGE Recovery Kit, Zymo Research #R1070). RNA concentration was measured using a NanoDrop. Purified RNA was refolded prior to sample vitrification as follows: Purified RNA was diluted to a target concentration of 20-30 µM in 50 mM Na-HEPES, pH 8.0, denatured at 90°C for 3 minutes, then cooled at room temperature for 10 minutes. RNA was incubated with 10 mM MgCl_2_ at 50°C for 20 minutes, then cooled at room temperature for 10 minutes. 3 μL of refolded RNA was frozen by Vitrobot Mark IV (2.5-4 seconds blot time, 1-5 seconds wait time) onto Quantifoil R 2/1 grids (Cu, 200 mesh) or Quantifoil R 1.2/1.3 grids (Cu, 300 mesh) following glow discharge (30 seconds glow, 15 seconds hold). Refer to **Supplemental Table S7** for details on target RNA concentrations, grid type, and sample freezing conditions for each sample.

### Cryo-EM data acquisition

Cryo-EM data was collected on a 300 kV Titan Krios G3i with a FEG electron source and autoloader cryo-specimen holder. Data for SARS-CoV-2 SL5-6 domains with SL5c extended and SARS-CoV-1 SL5 domain was collected on a Titan Krios with a Gatan K3 detector with a BioQuantum energy filter (20 eV slit width). Data for SARS-CoV-2 SL5 domain, SARS-CoV-2 SL5-6 domains, MERS SL5 domain, and BtCoV-HKU5 SL5 domain was collected on a Titan Krios with a Falcon 4 detector with no energy filter. Data for SARS-CoV-2 SL5-6 domains with SL5b extended; SARS-CoV-2 SL5-6 domains with SL6 extended and SL5a, SL5b, and SL5c removed; HCoV-229E SL5 domain; and HCoV-NL63 SL5 domain were collected on the same Titan Krios with a Falcon 4 detector with a Selectris energy filter (10 eV slit width). EPU was used for grid screening, data collection, and direct beam alignments, which included AutoFocus, AutoStigmatism, AutoComa, and objective aperture (100 µm) centering. Microscope alignments were performed using Digital Micrograph and included GIF tuning and ZLP centering with the K3 camera. For data collected with the Falcon 4 detector equipped with a Selectris energy filter, microscope alignments were performed using Sherpa. Micrographs were collected at nominal magnifications of 75kx to 165kx magnification at a total dose of 50-60 e^−^/Å² with a defocus range of -1.0 to -2.5 µm. Micrographs were gain-corrected during capture for the Falcon 4 detector or after capture for the K3 detector. Refer to **Supplemental Table S8** for exact values for nominal magnification, pixel size, total dose, dose per frame, frame duration, exposure time, and number of acquired micrographs.

### Cryo-EM data processing

Single-particle image processing and 3D reconstruction was performed using CryoSPARC 3.2.0 (57). Patch motion-correction and patch CTF-estimation were used in pre-processing. Information regarding the data processing for each dataset can be found in **Supplemental Table S9** and the pipelines to process each dataset can be seen in **Supplemental Figures S3, S7, S9, S11, S14,** and **Supplemental Note S2**. General strategies included one to three iterations of 2D classification with a larger box size to remove unfolded and aggregated RNA particles, which appear as elongated classes only when a larger box size is used; and 3D heterogeneous refinement using a spherical class to remove noise particles common when analyzing small particles which have a low signal-to-noise ratio.

### Modeling cryo-EM maps

All cryo-EM maps were modeled using auto-DRRAFTER, except for maps where major grooves were not resolved: the HCoV-NL63 SL5 and SARS-CoV-2 SL5-6 with SL6 extended and SL5a, SL5b, and SL5c removed. All secondary structures identified were used in separate auto-DRRAFTER runs, resulting in multiple ensembles of models for each map. Notably, for the SARS-CoV-2 SL5-6 domains, SARS-CoV-2 SL5-6 domains with SL5b extended, and SARS-CoV-2 SL5-6 domains with SL5c extended, the sequences and secondary structures were cut off at the bottom of the SL5-stem. For all other constructs, the full sequences were used for modeling. For all modeling, the sharpened map was used, except for the SARS-CoV-2 SL5-6 domains; however, given the low resolution of each map, the sharpened and unsharpened maps are not expected to result in different modeling results. Using autoDRRAFTER, nodes were fitted into the map after it was low pass filtered at 20 Å using a map threshold specified in the legend of **Supplemental Figures S4, S10,** and **S13**. The helical placements were exhaustively searched by initially placing the ends of the four stems in an “end-node,” as displayed in **Supplemental Figures S4, S10,** and **S13**. This node was consistent for each secondary structure modeled. Rounds of 5,000 decoys for each initial stem placement were run until convergence, defined as less than 10 Å mean pairwise r.m.s.d. between the top ten models. After these initial runs, two final rounds were run, also creating 5,000 decoys, to obtain a final top ten models. The exact commands can be found in the accompanying GitHub repository (https://github.com/DasLab/Coronavirus_SL5_3D). The ten final auto-DRRAFTER models were refined using ERRASER2 (https://new.rosettacommons.org/docs/latest/ERRASER2) with the following command in Rosetta:

erraser2

-s $PDB

-edensity:mapfile $MAP

-edensity::mapreso $RESOLUTION

-score:weights stepwise/rna/rna_res_level_energy7beta.wts

-set_weights elec_dens_fast 10.0 cart_bonded 5.0 linear_chainbreak 10.0

chainbreak 10.0 fa_rep 1.5 fa_intra_rep 0.5 rna_torsion 10 suiteness_bonus 5

rna_sugar_close 10

-rmsd_screen 3.0

-mute core.scoring.CartesianBondedEnergy core.scoring.electron_density.xray_scattering

-rounds 3

-fasta $FASTA

-cryoem_scatterers

The ten models were then combined into a single PDB file and pdb_extract (https://pdb-extract.wwpdb.org/) was used to convert them to mmCIF format. All jobs were run on the Stanford high performance computing cluster, Sherlock 2.0, using Rosetta 3.10 (2020.42).

### Model validation

The modeling convergence, mean pairwise heavy-atom r.m.s.d., was calculated using the Rosetta command:

drrafter_error_estimation -s $PDBs -mute core -rmsd_nosuper true -- per_residue_convergence true

The results can be found in **Supplemental Table S4**. The stereochemical and map-to-model scores were calculated using the pipeline (https://github.com/DasLab/CASP15_RNA_EM), which includes using MolProbity (58), Phenix cross-correlation scores, CC_volume_, CC_mask_, and CC_peaks_ (59), Q-score (42), and TEMPy for Mutual Information (MI) and segment-based Manders’ overlap coefficient (SMOC) scores (60). All calculations were carried out using the sharpened map and default parameters for each program. The mean per-residue convergence and Q-score of each ten model ensemble were then calculated, saved as B-factors on a representative structure, and visualized using ChimeraX (61) using in-house scripts. The average scores can be found in **Supplemental Figures S4, S10,** and **S13** and all scores can be found in **Supplemental Table S5**. Models from the EternaFold secondary structure for all BtCoV-HKU5 SL5 conformations and from library-based DMS M2-seq from BtCoV-HKU5 conformation 4 were found by these metrics to not fit in the map sufficiently well and hence were not considered further (**Supplemental Figure S10**). Finally, the effect of refinement using ERRASER2 on these validation metrics was plotted using in-house scripts. All scripts can be found in the accompanying GitHub repository (https://github.com/DasLab/Coronavirus_SL5_3D).

### Model analysis

The distance between UUYYGU hexaloops was defined as the distance between centroids of the C1’ atoms of the hexaloop. The inter-helical angle was defined as follows. A vector representing the SL5:SL5-stem stack was defined by minimizing the distance between this vector and all heavy atoms pointing away from SL5c towards SL5-stem. Likewise, a vector was defined for the SL5a:SL5b stack pointing towards SL5b. The angle of the SL5c-to-SL5-stem vector relative to the SL5a-to-SL5b vector was defined as the inter-helical angle, with clockwise defined as positive. The direction of view was defined with the SL5c-to-SL5-stem vector on top of the SL5a-to-SL5b vector. Hence a parallel configuration would result in 0° inter-helical angle and antiparallel a 180° inter-helical angle. The hinge angle was similarly defined but the vector was defined by the atoms in the apical residues SL5a stem-loop after the hinge, and a second vector as the remaining residues in the SL5a:SL5b stack. An angle of 0° would be a perfect coaxial stack, positive angle indicates bends towards SL5-stem, negative away from SL5-stem. See **Supplemental Figure S5** for a pictorial representation of these angles. The exact residues used to define the vectors can be found in the accompanying GitHub repository (https://github.com/DasLab/Coronavirus_SL5_3D). For figures, the pixel size of the SARS-CoV-2 SL5-6 domains map was increased from 1 Å/pixel to 1.1 Å/pixel to match other maps and the geometry of RNA A-form helices. Figures were prepared using ChimeraX (61) and scripts can be found in the accompanying GitHub repository (https://github.com/DasLab/Coronavirus_SL5_3D).

