## Supplemental Figures S1-14, Notes S1-2, Tables S1-3,7-9 for "Tertiary folds of the SL5 RNA from the 5′ proximal region of SARS-CoV-2 and related coronaviruses"

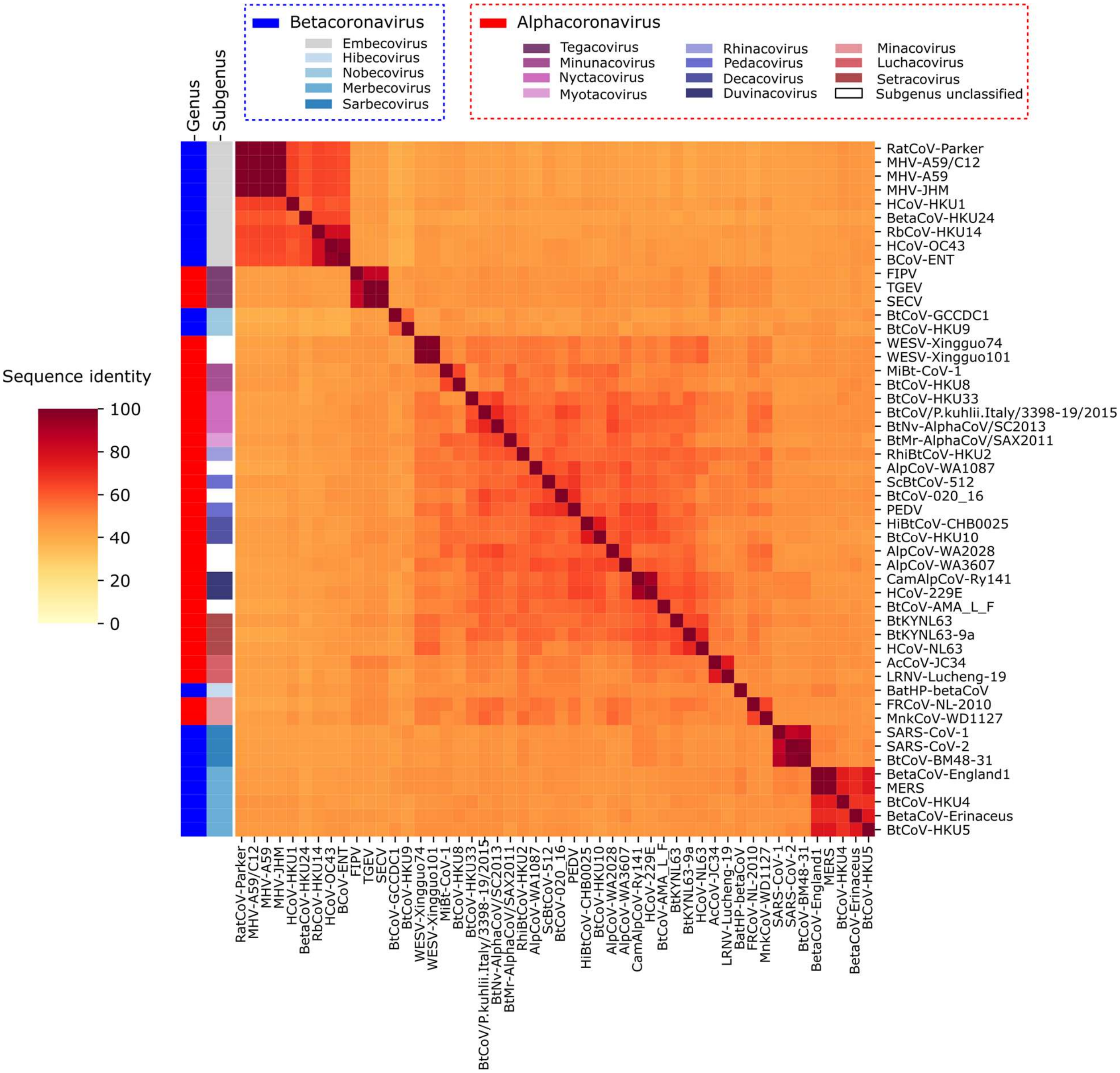

**Supplemental Figure S1: Sequence identity between reference alpha- and betacoronavirus SL5s.** All 54 reference genomes for alpha- and betacoronaviruses were retrieved from NCBI and the genomic location of SL5 was annotated for 51 of the sequences (see Supplemental Table S1). Each pair of SL5 sequences was then aligned and sequence identity was calculated; see Supplemental Note S1 for details. The sequence identity is plotted as a heat map from highest sequence identity in red to lowest sequence identity in yellow. On the left, the genus and subgenus of each virus is noted according to the color legend on top, with white representing a species that has not been classified into a subgenus. The average sequence identity over all pairwise comparisons depicted is 51%.

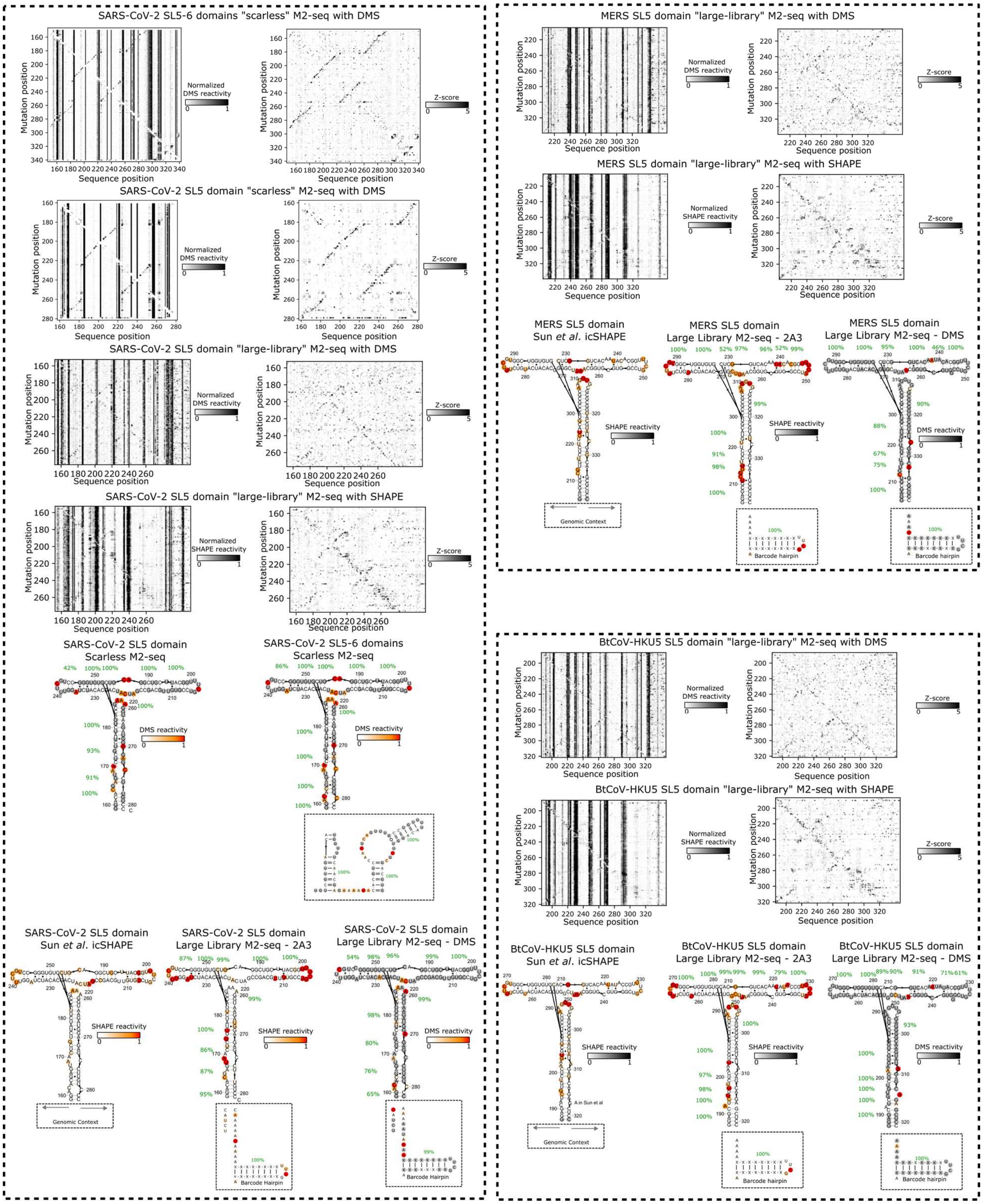

**Supplemental Figure S2: Map-and-mutate chemical mapping of SARS-CoV-2 SL5 and SL5-6, MERS SL5, and BtCoV-HKU5 SL5.** The black-and-white graphs are the results of mutate-and-map experiments. The left column contains the raw reactivity values across the sequence along the x-axis, with each row representing a mutated sequence. Loops can be identified as a dark bar representing consistently reactive nucleotides. For each sequence position, the Z-values (number of standard deviations above the mean) are taken and plotted on the right. A diagonal dark line is a signal for a stem. Note that the "large-library" samples had a 3' barcode stem-loop and the SARS-CoV-2 SL5 additionally had a 3' and 5' padding. Additionally, icSHAPE (1) and the SL5-6 domain were in a larger genomic context. The secondary structure diagrams are displayed and colored by reactivity. The green numbers represent the bootstrapped probability of the stem forming (100 bootstraps). The boxed regions below the secondary structure diagrams represent additional sequences that were present for the chemical mapping.

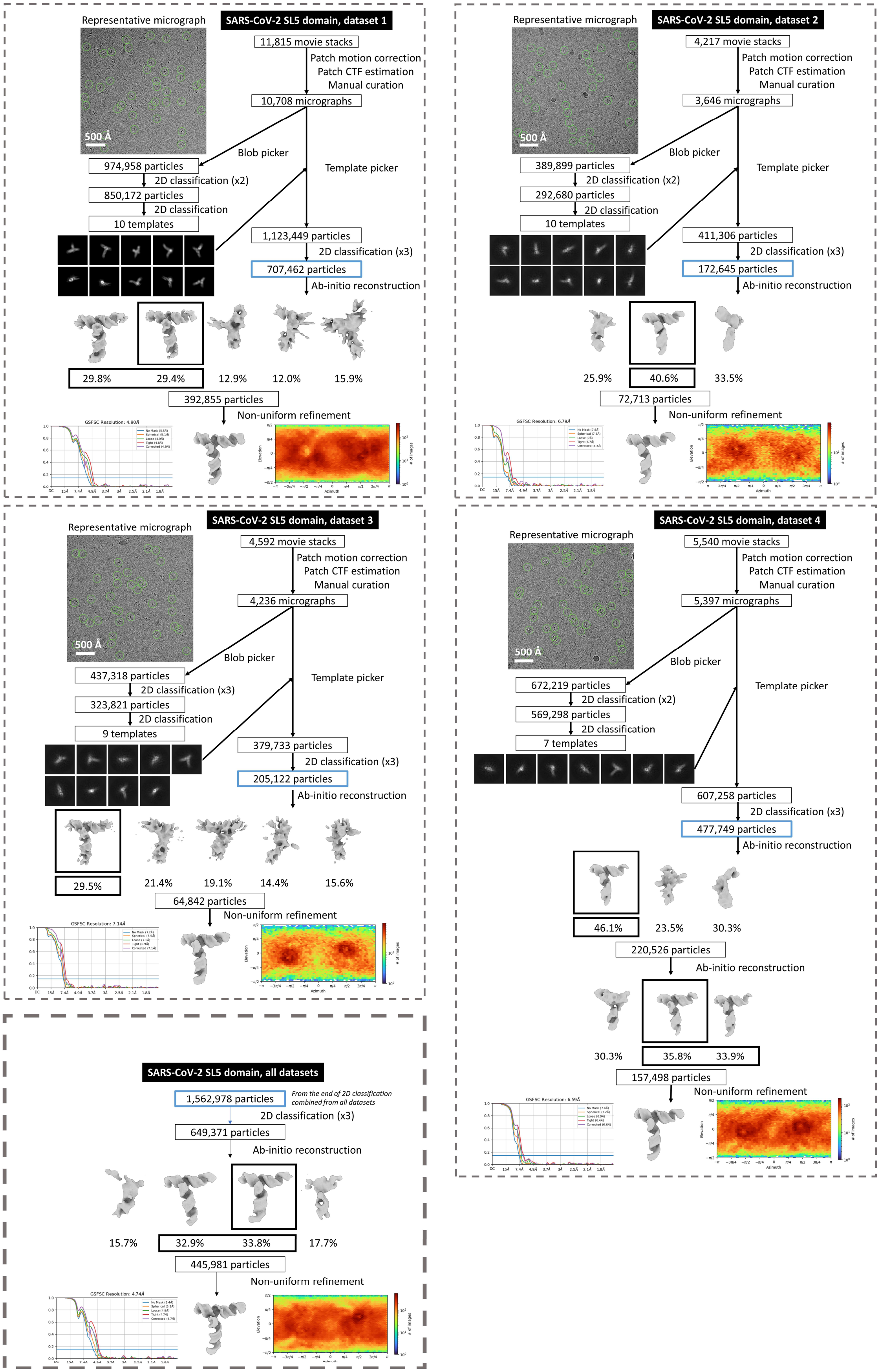

**Supplemental Figure S3: SARS-CoV-2 SL5 domain data processing flowchart.** See Supplemental Note S2 for details on interpreting this flowchart. The blue boxed particles for the four individual datasets are combined for the final analysis displayed on the bottom left.

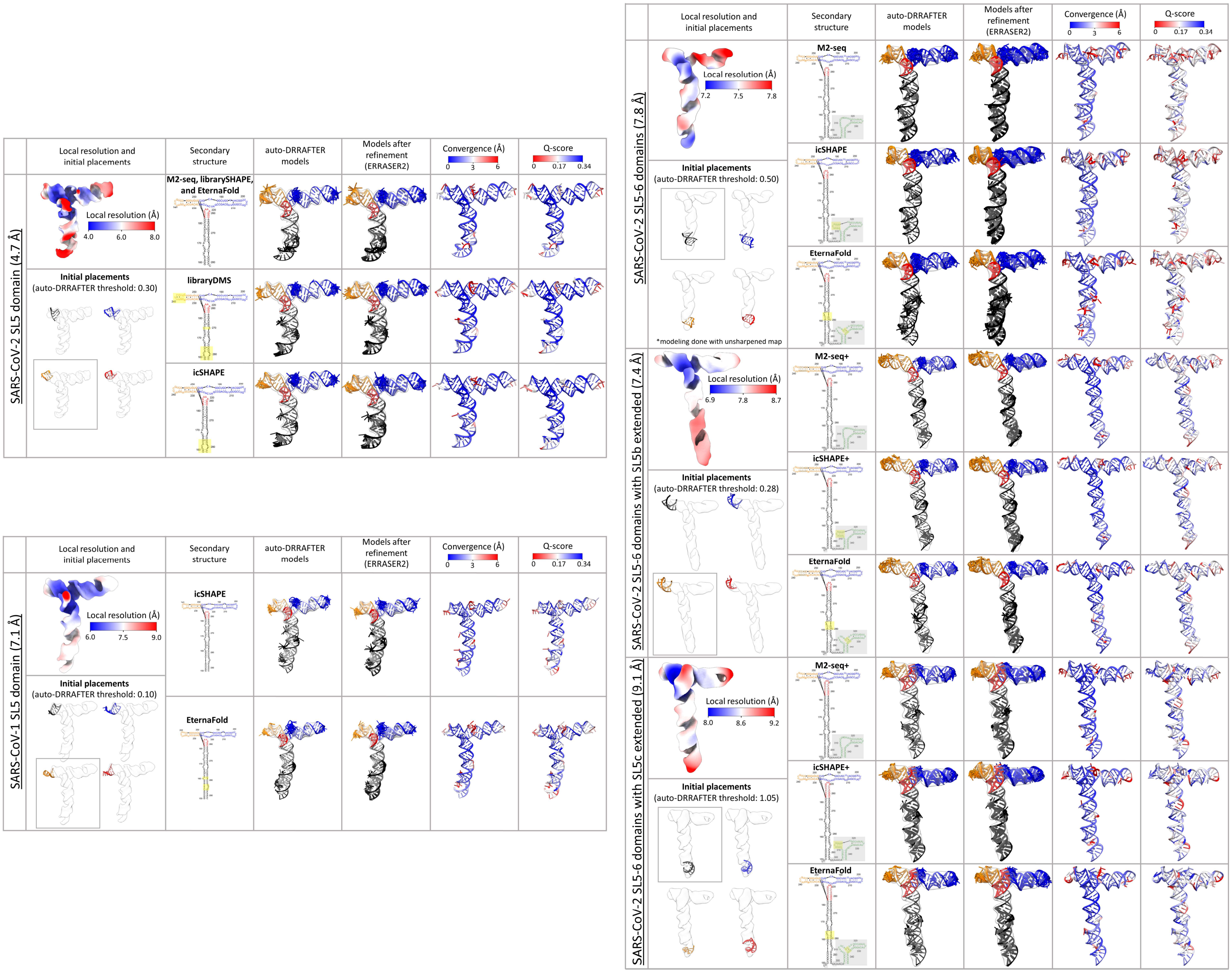

**Supplemental Figure S4: SARS-CoV-2 and SARS-CoV-1 SL5 domain modeling**. For each map, the local resolution, as estimated by CryoSPARC, is displayed in the left most column, from blue (higher resolution) to red (lower resolution). The secondary structure, depicting the base-pairs used as constraints during modeling, and its source are listed in the second column: M2-seq: derived from the “scarless” two-dimensional chemical mapping, librarySHAPE: derived from the “large-library” two-dimensional chemical mapping using 2A3, libraryDMS: derived from the “large-library” two-dimensional chemical mapping using DMS (Supplemental Figure S2), icSHAPE: as identified in Sun *et al.* (1), EternaFold: predicted by EternaFold with default settings (2), M2-seq+, icSHAPE+: derived the named secondary structure but assumes the stem extension forms as designed and does not change the secondary structure elsewhere. For each map, the differences in secondary structure, compared to the first row, are highlighted in yellow. Nucleotides highlighted with grey boxes were not included during modeling. They are colored by stem: SL5-stem in black, SL5a in blue, SL5b in orange, and SL5c in red. For each of the secondary structures listed, the map was modeled using auto-DRRAFTER (3) using a threshold and the initial helical placement depicted at the left of the tables. The top ten auto-DRRAFTER models are depicted, colored, and oriented to match the secondary structure diagram, before (column 3) and after refinement with ERRASER2 (4) (column 4). A representative model is colored by the per-nucleotide convergence, mean pairwise r.m.s.d., (column 5) and mean Q-score (column 6) of the ten models after refinement.

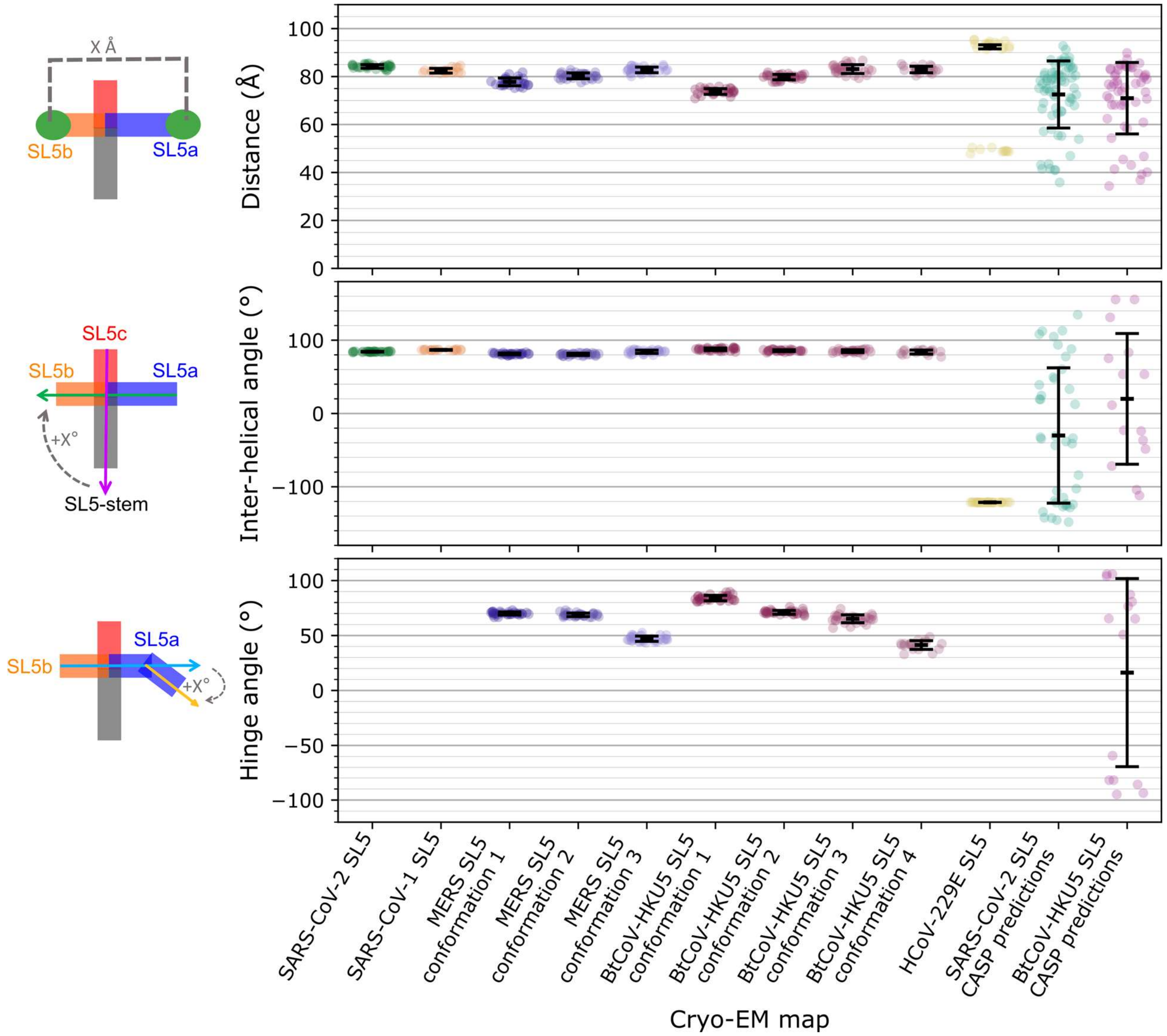

**Supplemental Figure S5: 3D features of the SL5 domains of coronaviruses**. The top plot shows the distance between the apical loops of SL5a and SL5b for each refined auto-DRRAFTER model in color. Distance is defined as the distance between the centroid of the C1' atoms of each loop. The middle plot shows the inter-helical angle between the SL5-stem:SL5c coaxial stack and SL5a:SL5b coaxial stack. The parallel configuration is defined as 0° and the anti-parallel configuration as 180°. The positive rotation is defined as rotating the SL5a:SL5b coaxial stack clockwise in the orientations in Figure 3. The bottom plot displays the hinge angle between the SL5a:SL5b coaxial stack and SL5a after the hinge at the asymmetric internal loop, with a positive value indicating rotation towards the SL5-stem and a negative value indicating rotation away from the SL5-stem. The measurements are described pictorially on the left. In all plots, the black bars represent the mean and standard deviation of the distances or angles across the ERRASER2 refined auto-DRRAFTER models. For the HCoV-229E SL5 distance measurement, only the distance between SL5a and SL5b hexaloops was considered for the error bar, but distances between the other pairs of hexaloops were plotted in the scatter plot. For the CASP models, only models that had the correct base-pairing at the junction (81/148 for SARS-CoV-2 SL5 and 92/185 for BtCoV-HKU5 SL5) were included in the top figure, and only models that had the correct base pairing and base stacking at the junction (50/148 for SARS-CoV-2 SL5 and 44/185 for BtCoV-HKU5 SL5) were included in the middle and bottom plots.

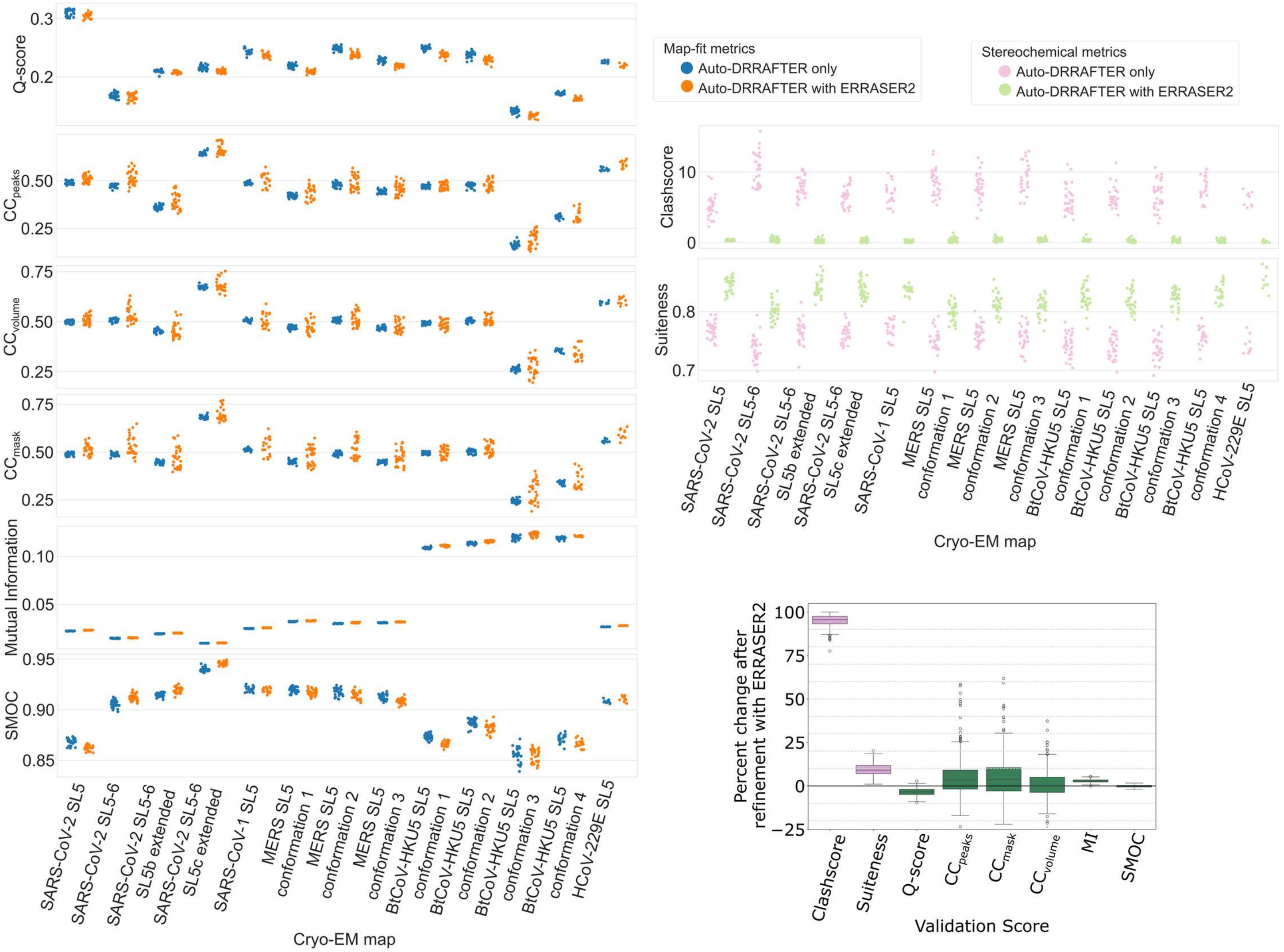

**Supplemental Figure S6: Stereochemical and map-fit metrics before and after refinement with ERRASER2.** The stereochemical metrics (Clashscore (5) and Suiteness (6)) are calculated for each of the models fit into each map. The left plots show scores before refinement in pink and scores after refinement in green. Note that for Clashscore, a lower score shows improvement, whereas for Suiteness, a higher score shows improvement. The map-fit metrics (mutual information (MI), segment-based Manders' overlap coefficient (SMOC), (7) $CC_{peaks}$, $CC_{volume}$, $CC_{mask}$, (8) and Q-score (9)), are also calculated for the models for each map, with scores before refinement in blue and scores after refinement in orange. For each of these scores, a higher score shows improvement in the map-model fit. The insert on the right shows the percent change in the validation scores for the top ten models fit into a single map. The box shows the quartiles of the dataset, the whiskers show the rest of the distribution, and outliers (more than 1.5 times the inter-quartile range below the first quartile or above the third quartile) are shown as points past the limits of the whiskers. For Clashscore, the percent change was multiplied by -1 so that a positive percentage change represents improvement across all scores.

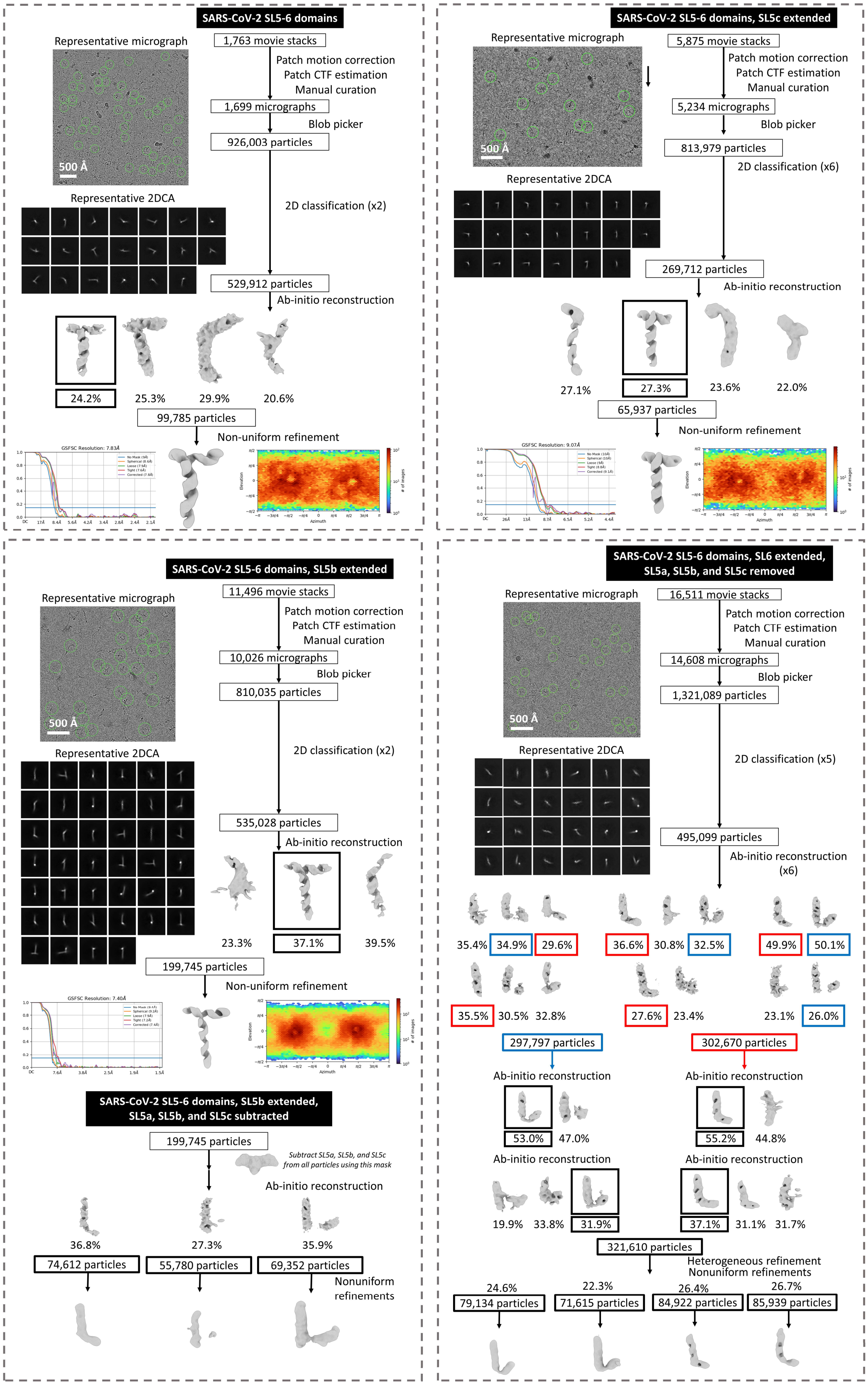

**Supplemental Figure S7: SARS-CoV-2 SL5-6 domains and extension constructs data processing flowcharts.** See Supplemental Note S2 for details on interpreting these flowcharts. Note that on the bottom left, SL5a, SL5b, and SL5c are digitally removed but in the bottom right, SL5a, SL5b, and SL5c are biochemically removed (the construct imaged does not contain them).

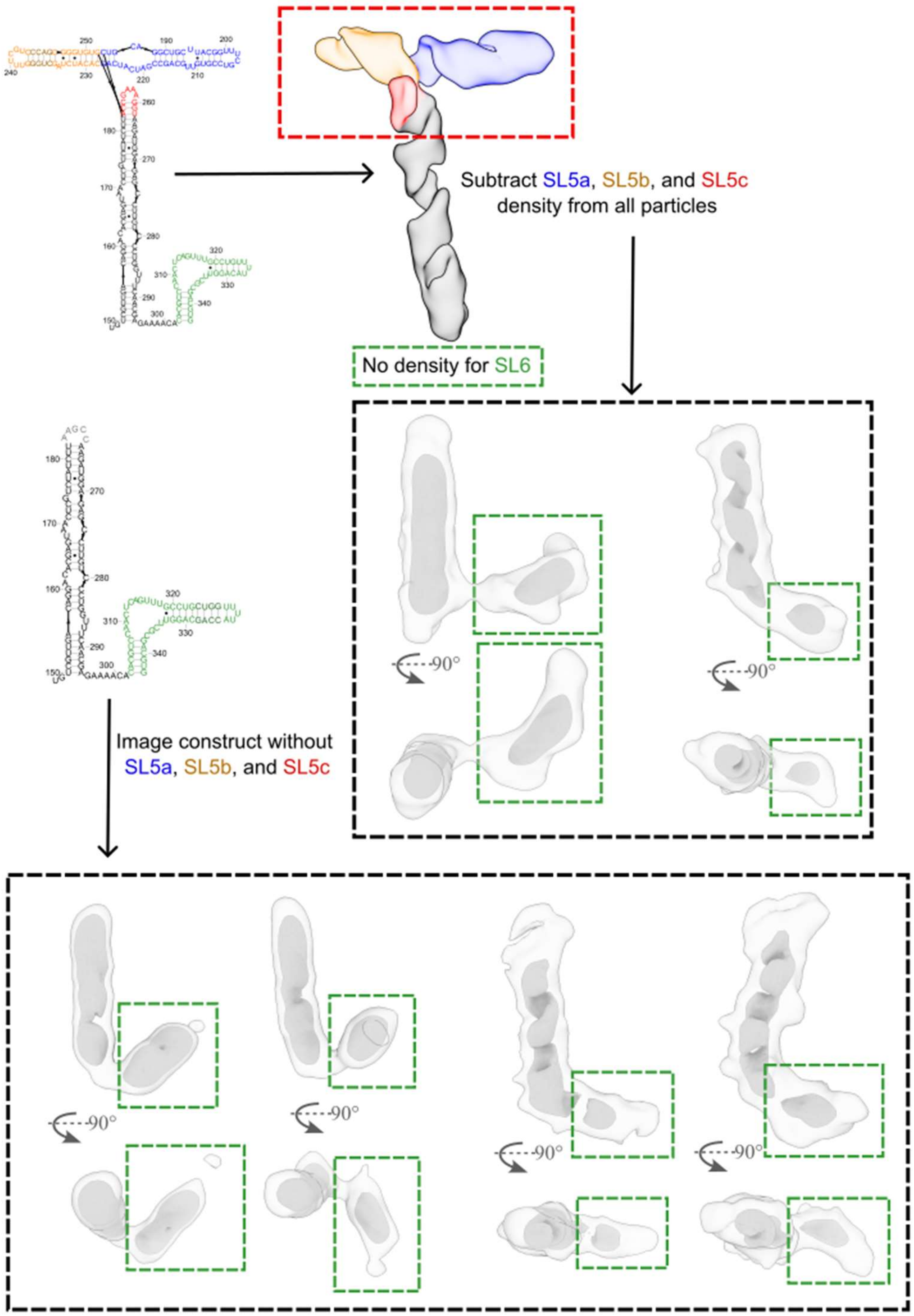

**Supplemental Figure S8: Flexible linker between the SL5 domain and SL6 domain in SARS-CoV-2.** The secondary structure of the SARS-CoV-2 SL5-6 construct with SL5b extended is displayed in the top left corner. The cryo-EM map is colored by domain, showing that it lacks density for the SL6 domain, but the other stems all have well resolved helical grooves. The density for SL5a, SL5b, and SL5c is subtracted for particle images (Supplemental Figure S7), resulting in the cryo-EM maps boxed in black. The putative density for the SL6 domain is boxed in green. A construct that lacks SL5a, SL5b, and SL5c, was also imaged (Supplemental Figure S7) and the resulting cryo-EM maps are displayed at the bottom of the figure, with the putative SL6 density boxed in green. The cryo-EM maps resulting from experimentally removing the SL5a, SL5b, and SL5c stems and digitally removing their densities appear similar.

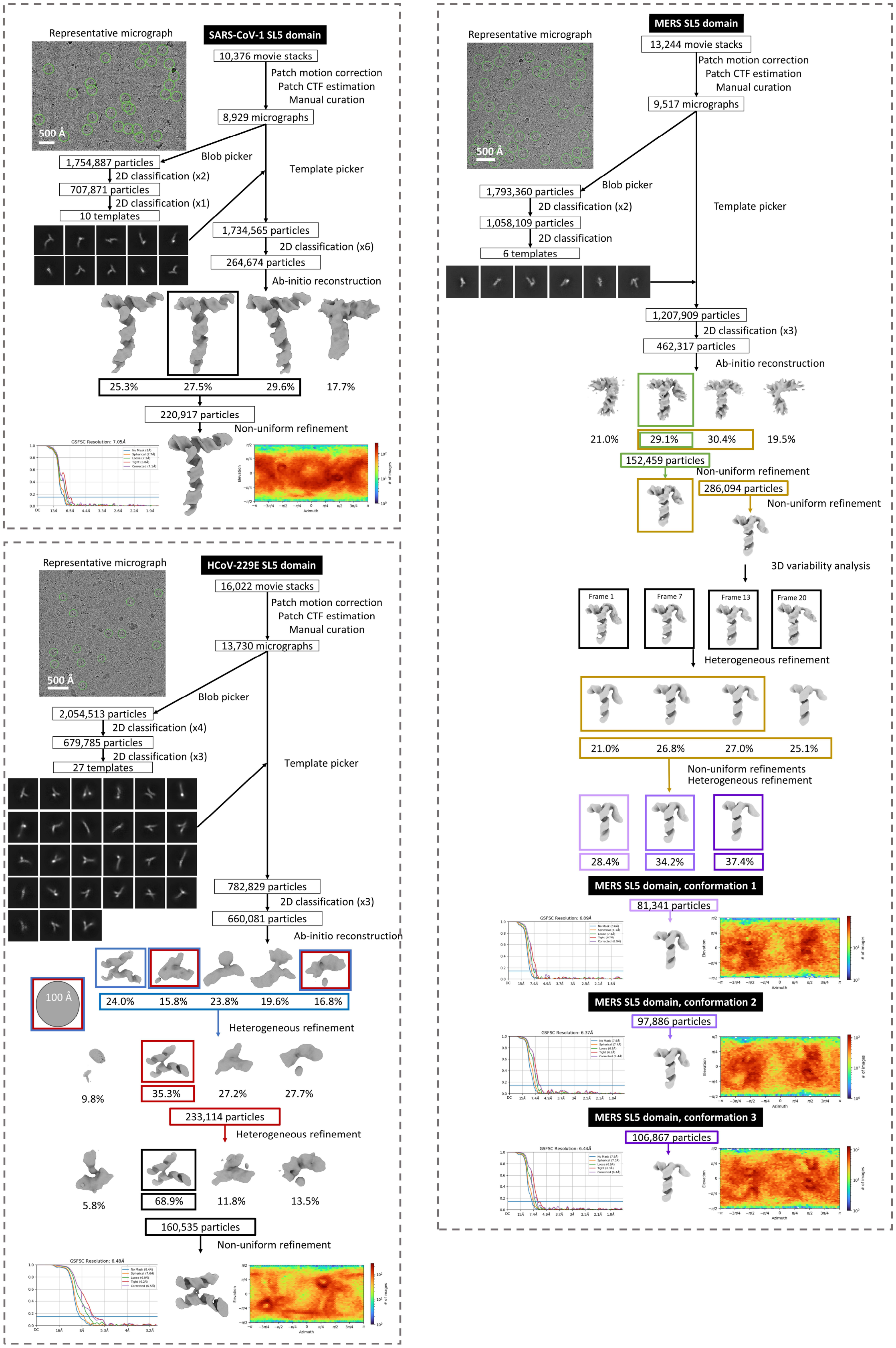

**Supplemental Figure S9: SARS-CoV-1, MERS, and HCoV-229E SL5 domain data processing flowcharts.** See Supplemental Note S2 for details on interpreting these flowcharts.

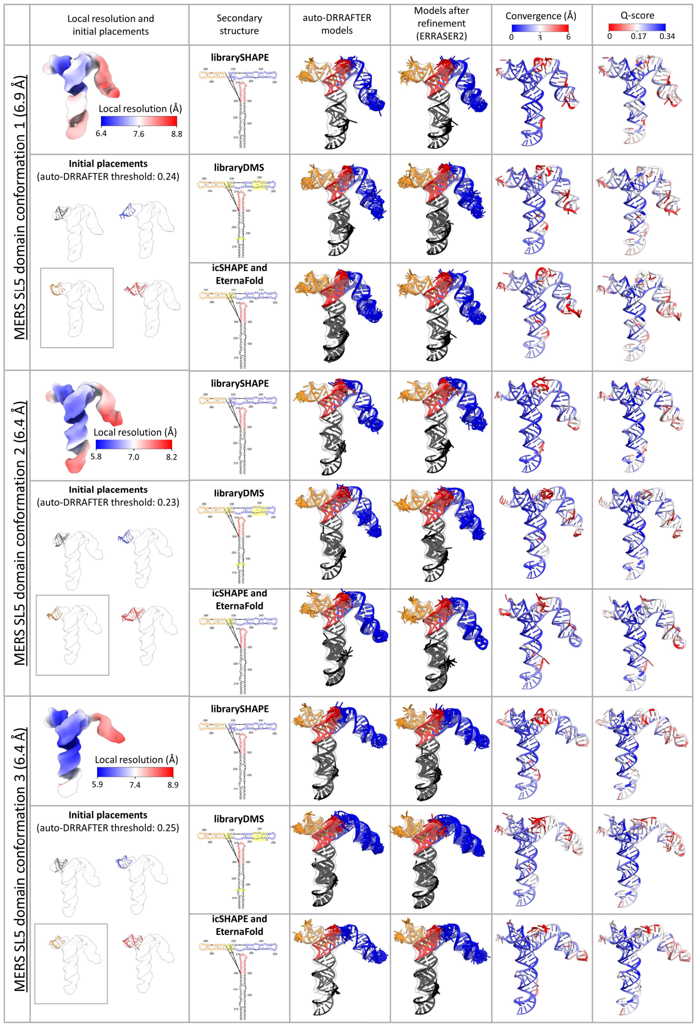

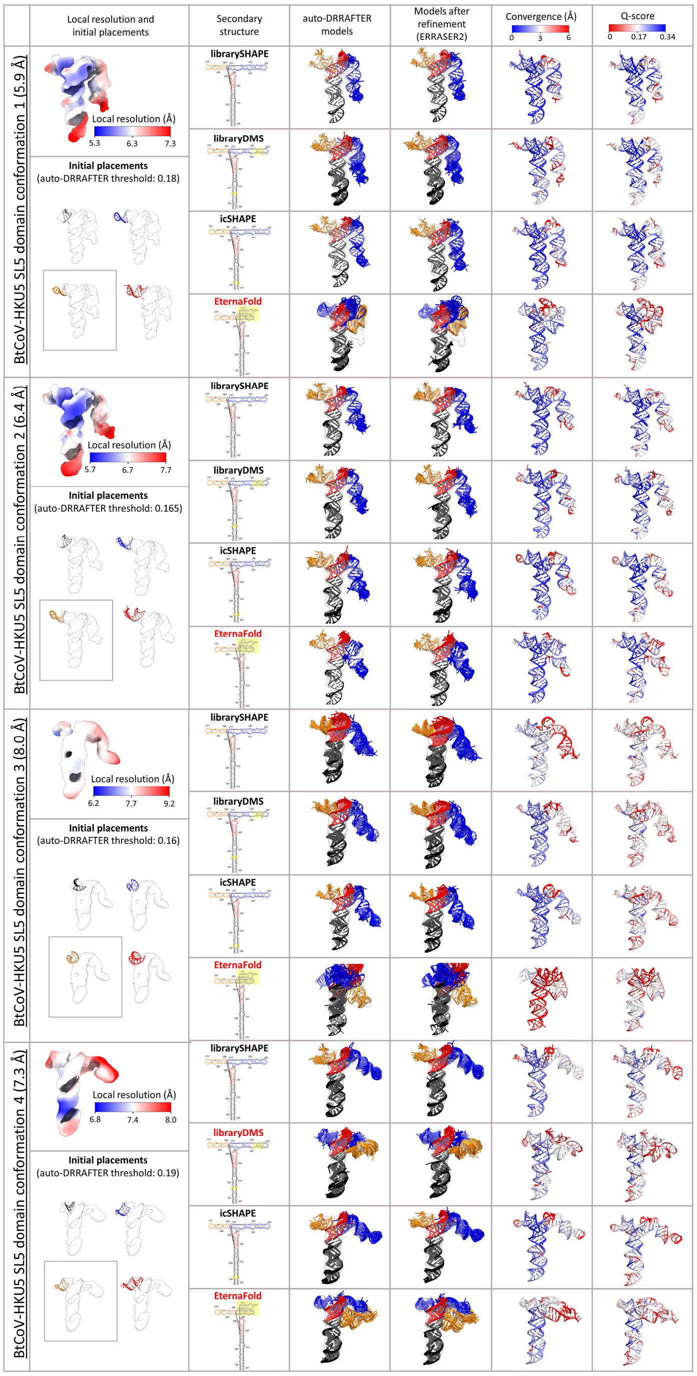

**Supplemental Figure S10: MERS and BtCoV-HKU5 SL5 domain modeling.** For each map, the local resolution, as estimated by CryoSPARC, is displayed in the left most column, from blue (higher resolution) to red (lower resolution). The secondary structure, depicting the base-pairs used as constraints during modeling, and its source are listed in the second column: librarySHAPE: derived from the “large-library” two-dimensional chemical mapping using 2A3, libraryDMS: derived from the “large-library” two-dimensional chemical mapping using DMS (Supplemental Figure S2), icSHAPE: as identified in Sun *et al.* (1), EternaFold: predicted by EternaFold with default settings (2). For each map, the differences in secondary structure, compared to the first row, are highlighted in yellow. They are colored by stem: SL5-stem in black, SL5a in blue, SL5b in orange, and SL5c in red. For each of the secondary structures listed, the map was modeled using auto-DRRAFTER (3) using a threshold and the initial helical placement depicted at the left of the tables. The top ten auto-DRRAFTER models are depicted, colored, and oriented to match the secondary structure diagram, before (column 3) and after refinement with ERRASER2 (4) (column 4). A representative model is colored by the per-nucleotide mean pairwise r.m.s.d., convergence, (column 5) and mean Q-score (column 6) of the ten models after refinement. Models with red titled secondary structure were not used because of poor fit in the map.

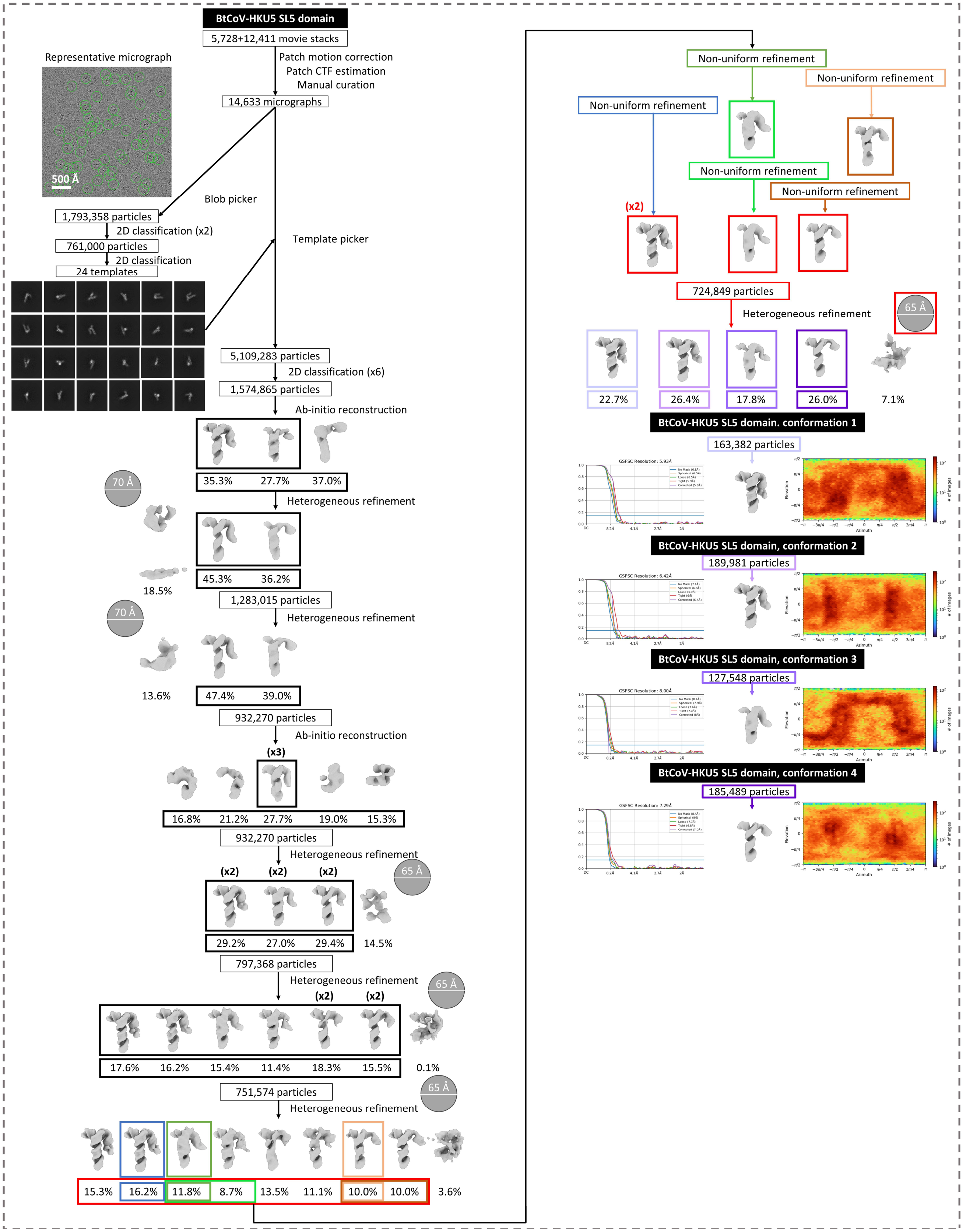

**Supplemental Figure S11: BtCoV-HKU5 SL5 domain data processing flowchart.** See Supplemental Note S2 for details on interpreting this flowchart.

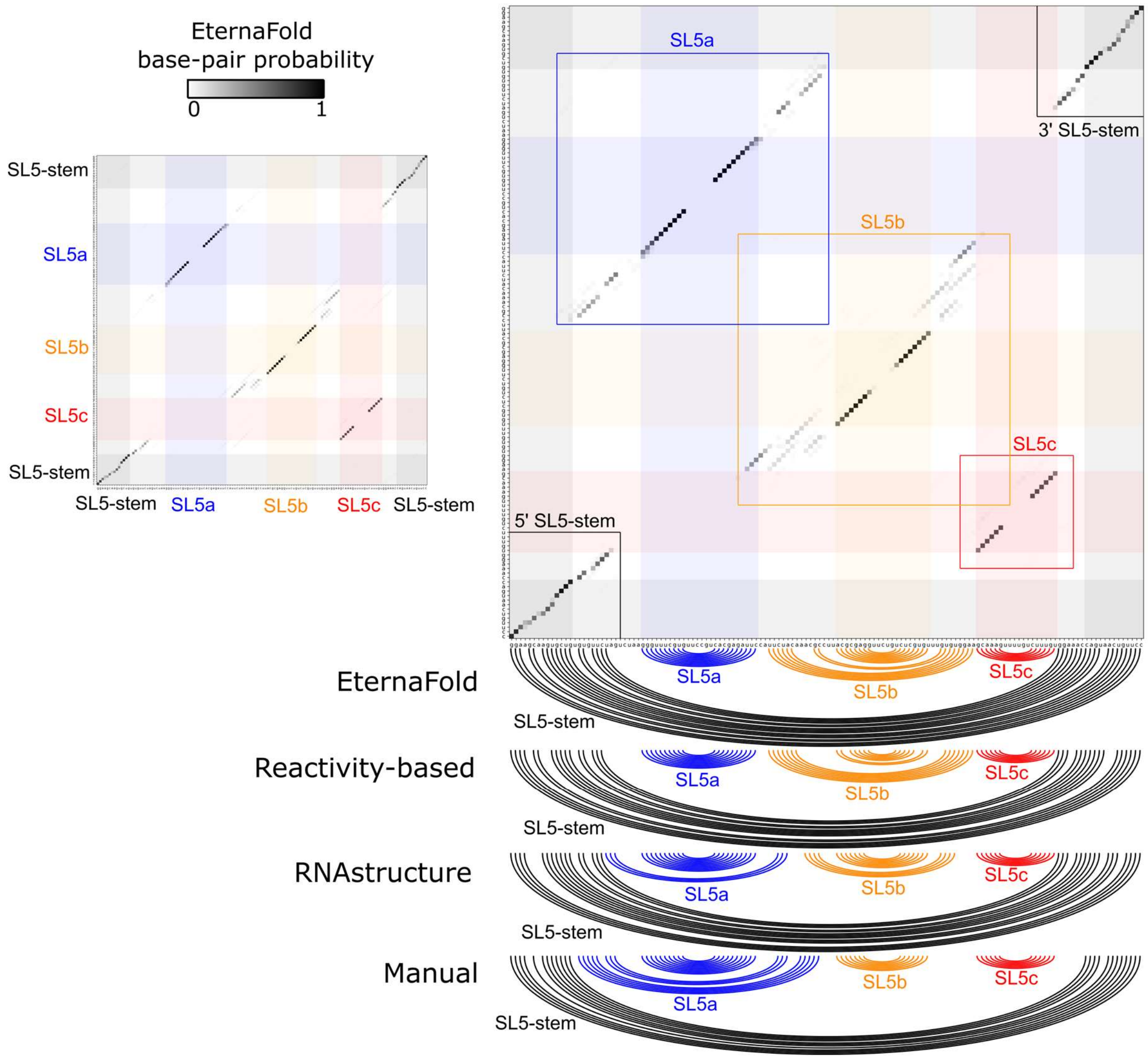

**Supplemental Figure S12: HCoV-229E base pair probability matrices and secondary structure.** The EternaFold (2) predicted base pair-probability matrix for the HCoV-229E SL5 sequence is displayed on the left, with high probability of base-pairing in black. The regions of high probability were identified and shaded. Sub-sequences were predicted for each stem, encompassing the high confidence region as well as the more uncertain in-between regions. The base-pair probability predictions from EternaFold for the sub-sequences are displayed in the colored, boxes regions on the right. Note the black boxes were predicted together; the 3' end of the left box was appended to the 5' end of the right box. The base-pairing derived from EternaFold with default settings (2), the structure predicted using unpublished SHAPE reactivity (10), minimum free energy structure using RNAfold with default parameters (11), and a manually created alternative based on this base-pair probability matrix and sequence alignments are displayed as arc diagrams below the base-pair probability matrix.

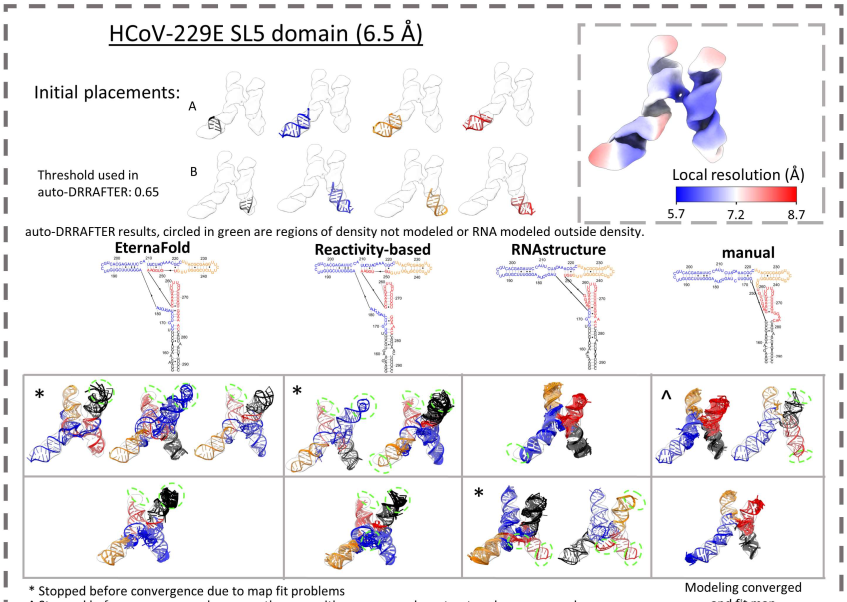

* Stopped before convergence due to map fit problems
^ Stopped before convergence because other run with same secondary structure has converged

Modeling converged and fit map

| Secondary structure | Conv. (Å) | Q-score | Clashscore | Average suiteness | $CC_{volume}$ | $CC_{mask}$ | $CC_{peaks}$ | MI | SMOC |
|---|---|---|---|---|---|---|---|---|---|
| EternaFold A | 33.82* | 0.161 | 8.418 | 0.729 | 0.436 | 0.386 | 0.349 | 0.023 | 0.844 |
| EternaFold B | 3.20 | 0.182 | 10.596 | 0.729 | 0.501 | 0.459 | 0.439 | 0.024 | 0.874 |
| Reactivity A | 16.34* | 0.175 | 8.531 | 0.736 | 0.474 | 0.427 | 0.402 | 0.023 | 0.861 |
| Reactivity B | 3.74 | 0.193 | 8.305 | 0.732 | 0.519 | 0.474 | 0.455 | 0.024 | 0.878 |
| RNAstructure A | 3.67 | 0.184 | 9.497 | 0.719 | 0.504 | 0.456 | 0.435 | 0.024 | 0.875 |
| RNAstructure B | 18.57* | 0.161 | 10.306 | 0.735 | 0.446 | 0.392 | 0.353 | 0.024 | 0.839 |
| Manual A | 14.57^ | 0.204 | 7.522 | 0.726 | 0.551 | 0.504 | 0.503 | 0.025 | 0.889 |
| **Manual B** | 2.19 | **0.226** | 6.264 | 0.742 | **0.594** | **0.558** | **0.560** | **0.027** | **0.908** |

Single secondary structure converged and was refined:

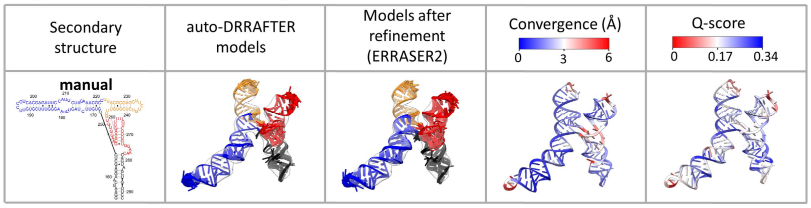

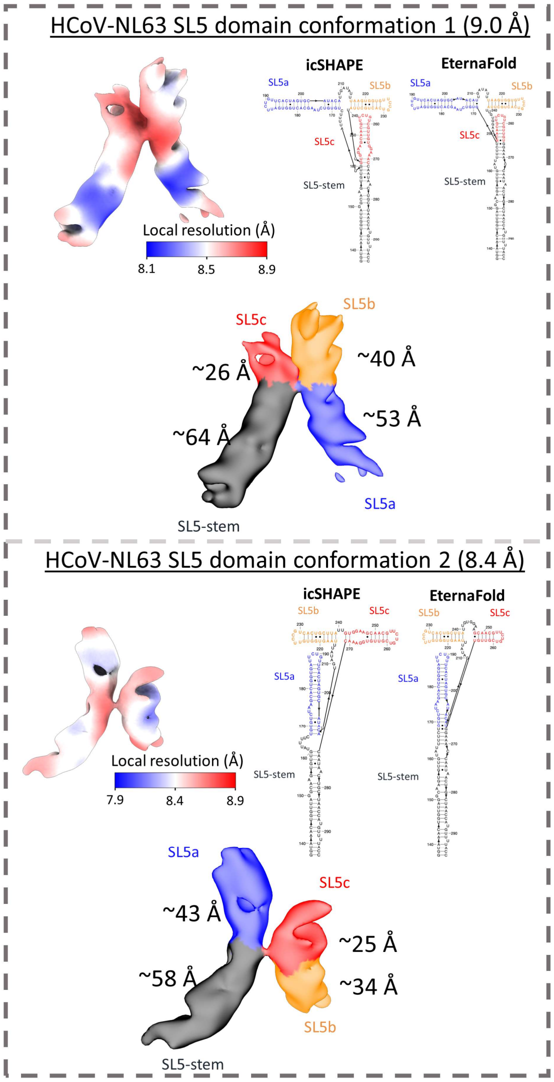

**Supplemental Figure S13: HCoV-229E and HCoV-NL63 SL5 domain modeling.** For each map, the local resolution, as estimated by CryoSPARC, is displayed by the title, from blue (higher resolution) to red (lower resolution). For HCoV-NL63, the map was modeled using auto-DRRAFTER (3) using a threshold and the initial helical placement depicted at the top of the figure. In the table below, the top ten auto-DRRAFTER models are depicted and colored matching the secondary structure which are depicted above the column (Supplemental Figure S12). Underneath, the validation scores for these models are displayed showing only one secondary structure; only models corresponding to manual had reasonable scores. In the bottom table the top ten auto-DRRAFTER models are depicted before (column 3) and after refinement with ERRASER2 (4) (column 4). A representative model is colored by the per-nucleotide mean pairwise r.m.s.d., convergence, (column 5) and mean Q-score (column 6) of the ten models after refinement. For both HCoV-NL63 conformations, the two secondary structures, are shown on the right, arranged according to the 3D arrangement of stems: icSHAPE: as identified in Sun *et al.* (1), EternaFold: predicted by EternaFold with default settings (2). In the center, the cryo-EM density is displayed with each stem annotated with its approximate length. The relative lengths of the cryo-EM densities enable the assignment of the stems based on the secondary structure. The cryo-EM map is colored by these assignments.

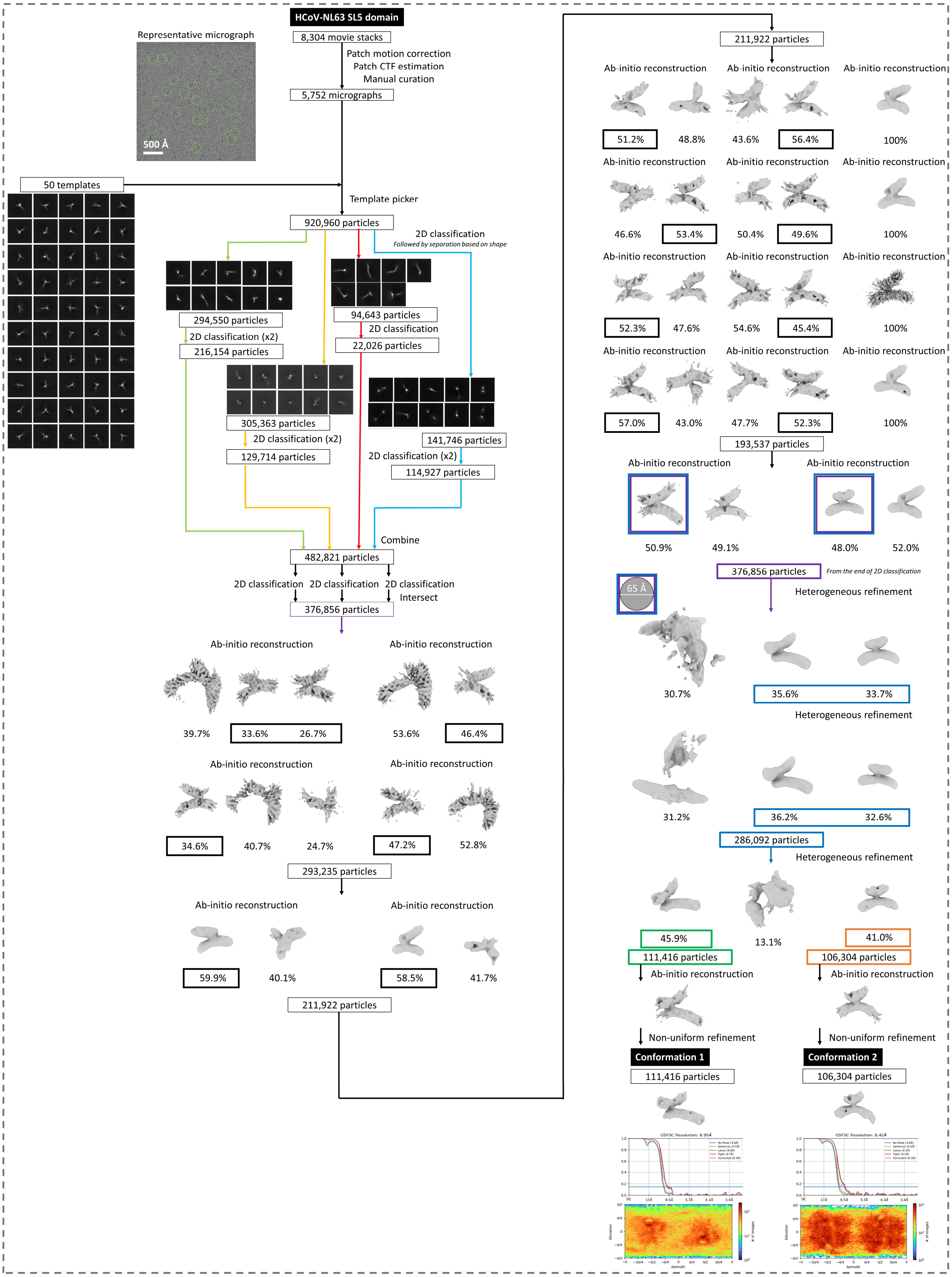

**Supplemental Figure S14: HCoV-NL63 SL5 domain data processing flowchart.** See Supplemental Note S2 for details on interpreting this flowchart.

**Supplemental Table S1: Alpha- and betacoronavirus reference genomes' SL5 properties.**

| NCBI reference genome | Genus | Subgenus | Species | Start codon location | SL5 location^ | Predicted apical loop sequence | | | Predicted number of nucleotides in stem | | | |
|---|---|---|---|---|---|---|---|---|---|---|---|---|
| | | | | | | SL5a | SL5b | SL5c | SL5-stem | SL5a | SL5b | SL5c |
| **NC_012936.1** | Betacoronavirus | Embecovirus | RatCoV-Parker | 183 | 114-336 | cacccaua | uuccgaac | uuuug | 56 | 66 | 81 | 20 |
| **NC_001846.1** | Betacoronavirus | Embecovirus | MHV-A59/C12 | 210 | 141-363 | cacccaua | uuccgaac | uuuug | 56 | 66 | 81 | 20 |
| **NC_048217.1** | Betacoronavirus | Embecovirus | MHV-A59 | 211 | 142-364 | cacccaua | uuccgaac | uuuug | 56 | 66 | 81 | 20 |
| **AC_000192.1** | Betacoronavirus | Embecovirus | MHV-JHM | 215 | 146-368 | cacccaua | uuccgaac | uuuug | 56 | 66 | 81 | 20 |
| **NC_006577.2** | Betacoronavirus | Embecovirus | HCoV-HKU1* | 206 | 139-359 | cccau | uuccggau | uuaug | 55 | 60 | 83 | 20 |
| **NC_026011.1** | Betacoronavirus | Embecovirus | BetaCoV-HKU24 | 213 | 140-369 | cacccaua | ugag | gguuuug | 54 | 74 | 77 | 22 |
| **NC_017083.1** | Betacoronavirus | Embecovirus | RbCoV-HKU14 | 209 | 140-362 | cacccaua | ugag | ugguu | 56 | 66 | 79 | 22 |
| **NC_006213.1** | Betacoronavirus | Embecovirus | HCoV-OC43* | 210 | 142-363 | cgcccaua | ugag | uaugauuugc | 51 | 70 | 74 | 26 |
| **NC_003045.1** | Betacoronavirus | Embecovirus | BCoV-ENT | 211 | 143-364 | cacccaua | ugag | uauaguaugc | 51 | 70 | 74 | 26 |
| **NC_002306.3** | Alphacoronavirus | Tegacovirus | FIPV | 311 | 153-323 | uuccgu | uuccgc | uuccgu | 76 | 36 | 44 | 14 |
| **NC_038861.1** | Alphacoronavirus | Tegacovirus | TGEV | 315 | 156-327 | uuccgu | uuccgc | uuccgu | - | - | - | - |
| **NC_028806.1** | Alphacoronavirus | Tegacovirus | SECV | 307 | 148-319 | uuccgc | uuccgc | uuccgu | - | - | - | - |
| **NC_030886.1** | Betacoronavirus | Nobecovirus | BtCoV-GCCDC1 | 235 | 147-264 | uuccgu | uuccgc | | 58 | 33 | 27 | 0 |
| **NC_009021.1** | Betacoronavirus | Nobecovirus | BtCoV-HKU9 | 229 | 145-258 | ugucgu | uuucgu | | 54 | 31 | 25 | 0 |
| **NC_048211.1** | Alphacoronavirus | | WESV-Xingguo74 | 212 | 78-225 | uucug | uuccgu | uuccgu | 56 | 33 | 27 | 28 |
| **NC_035191.1** | Alphacoronavirus | | WESV-Xingguo101 | 228 | 94-241 | uucug | uuccgu | uuccgu | 56 | 33 | 27 | 28 |
| **NC_010437.1** | Alphacoronavirus | Minunacovirus | MiBt-CoV-1 | 272 | 123-275 | uuccgu | uuccgu | uuccgu | 39 | 55 | 27 | 30 |
| **NC_010438.1** | Alphacoronavirus | Minunacovirus | BtCoV-HKU8 | 270 | 124-273 | uuccgu | uucugu | uuccgu | 39 | 53 | 26 | 30 |
| **NC_054015.1** | Alphacoronavirus | Nyctacovirus | BtCoV-HKU33 | 278 | 124-290 | uuccgu | uuccau | uucugu | 59 | 53 | 30 | 23 |
| **NC_046964.1** | Alphacoronavirus | Nyctacovirus | BtCoV/P.kuhlii.Italy/3398 | 211 | 125-293 | uuccgu | uuccgu | uuccgu | 53 | 57 | 22 | 29 |
| **NC_028833.1** | Alphacoronavirus | Nyctacovirus | BtNv-AlphaCoV/SC2013 | 214 | 59-226 | uuccgu | uucugu | uuccgu | 62 | 45 | 34 | 18 |
| **NC_028811.1** | Alphacoronavirus | Myotacovirus | BtMr-AlphaCoV/SAX2011 | 299 | 142-311 | uuccgu | uuccgu | uuccgu | 51 | 54 | 29 | 31 |
| **NC_009988.1** | Alphacoronavirus | Rhinacovirus | RhiBtCoV-HKU2 | 297 | 143-309 | uuccgu | uucugu | uuccgu | 56 | 54 | 25 | 30 |
| **NC_054003.1** | Alphacoronavirus | | AlpCoV-WA1087 | 295 | 149-312 | uuccgu | uuccgu | uucugu | 49 | 57 | 26 | 30 |
| **NC_009657.1** | Alphacoronavirus | Pedacovirus | ScBtCoV-512 | 294 | 135-306 | uuccgu | uuuugu | uuccgu | 57 | 57 | 26 | 30 |
| **NC_076629.1** | Alphacoronavirus | | BtCoV-020_16 | 270 | 113-282 | uuccgu | uuccgu | uuccgu | 45 | 57 | 22 | 40 |
| **NC_003436.1** | Alphacoronavirus | Pedacovirus | PEDV | 297 | 137-309 | uuccgu | uuccgu | uuccgu | 58 | 57 | 26 | 27 |
| **NC_054004.1** | Alphacoronavirus | Decacovirus | HiBtCoV-CHB0025 | 284 | 132-288 | uuccgu | uuccgu | ucua | 44 | 54 | 27 | 31 |
| **NC_018871.1** | Alphacoronavirus | Decacovirus | BtCoV-HKU10 | 303 | 151-307 | uuccguc | uuccgu | uuccgu | 44 | 54 | 27 | 30 |
| **NC_076684.1** | Alphacoronavirus | | AlpCoV-WA2028 | 435 | 279-447 | uuccgu | uuccgu | uuccgu | 62 | 46 | 36 | 18 |
| **NC_076685.1** | Alphacoronavirus | | AlpCoV-WA3607 | 238 | 90-259 | uuccgu | uuccgu | uuccgu | 56 | 55 | 27 | 30 |

| NCBI reference genome | Genus | Subgenus | Species | Start codon location | SL5 location^ | Predicted apical loop sequence | | | Predicted number of nucleotides in stem | | | |
|---|---|---|---|---|---|---|---|---|---|---|---|---|
| | | | | | | SL5a | SL5b | SL5c | SL5-stem | SL5a | SL5b | SL5c |
| **NC_028752.1** | Alphacoronavirus | Duvinacovirus | CamAlpCoV-Ry141 | 293 | 137-305 | uucugu | uucug | uucugu | - | - | - | - |
| **NC_002645.1** | Alphacoronavirus | Duvinacovirus | HCoV-229E* | 293 | 137-305 | uuccgu | uuccg | uucugu | 69 | 26 | 46 | 18 |
| **NC_055953.1** | Alphacoronavirus | | BtCoV-AMA_L_F | 242 | 87-254 | uuccgu | uucugu | uucugu | 57 | 52 | 27 | 30 |
| **NC_048216.1** | Alphacoronavirus | Setracovirus | BtKYNL63 | 246 | 93-255 | uuccgu | uuccgu | uuccgu | 62 | 47 | 31 | 16 |
| **NC_032107.1** | Alphacoronavirus | Setracovirus | BtKYNL63-9a | 242 | 87-254 | uuccgu | uuccgu | uucugu | 56 | 54 | 26 | 30 |
| **NC_005831.2** | Alphacoronavirus | Setracovirus | HCoV-NL63* | 287 | 136-297 | uucugu | uuccgu | uucugu | 65 | 40 | 24 | 18 |
| **NC_034972.1** | Alphacoronavirus | Luchacovirus | AcCoV-JC34 | 437 | 277-446 | uuccggc | uucugc | uuccgu | 55 | 57 | 18 | 33 |
| **NC_032730.1** | Alphacoronavirus | Luchacovirus | LRNV-Lucheng-19 | 332 | 169-341 | uuccgc | guucc | uuccgac | 57 | 55 | 22 | 31 |
| **NC_025217.1** | Betacoronavirus | Hibecovirus | BatHP-betaCoV | 303 | 152-330 | uuucgu | uuucgu | uuucgu | 63 | 45 | 23 | 48 |
| **NC_030292.1** | Alphacoronavirus | Minacovirus | FRCoV-NL-2010 | 276 | 140-288 | uuccgu | uuccgu | uuccgu | 61 | 31 | 32 | 16 |
| **NC_023760.1** | Alphacoronavirus | Minacovirus | MnkCoV-WD1127 | 273 | 131-292 | uuccgu | uuccgu | uuccgu | 60 | 32 | 30 | 26 |
| **NC_004718.3** | Betacoronavirus | Sarbecovirus | SARS-CoV-1* | 265 | 149-293 | uuucgu | uuucgu | gaaa | 65 | 45 | 25 | 10 |
| **NC_045512.2** | Betacoronavirus | Sarbecovirus | SARS-CoV-2* | 266 | 150-294 | uuucgu | uuucgu | gaaa | 65 | 45 | 25 | 10 |
| **NC_014470.1** | Betacoronavirus | Sarbecovirus | BtCoV-BM48-31 | 191 | 75-219 | uuucgu | uuucgu | gaaa | 65 | 45 | 25 | 10 |
| **NC_038294.1** | Betacoronavirus | Merbecovirus | BetaCoV-England1 | 278 | 195-348 | uuucgu | uuucgu | aaggugc | 61 | 44 | 27 | 21 |
| **NC_019843.3** | Betacoronavirus | Merbecovirus | MERS* | 279 | 196-349 | uuucgu | uuucgu | aaggugc | 61 | 44 | 27 | 21 |
| **NC_009019.1** | Betacoronavirus | Merbecovirus | BtCoV-HKU4 | 267 | 183-337 | uuccgu | ucca | aaggugc | 62 | 44 | 27 | 21 |
| **NC_039207.1** | Betacoronavirus | Merbecovirus | BetaCoV-Erinaceus | 241 | 158-311 | uuccgu | uuccgc | aaggugc | 61 | 44 | 27 | 21 |
| **NC_009020.1** | Betacoronavirus | Merbecovirus | BtCoV-HKU5 | 261 | 178-331 | uuucgu | uuucgu | aaggugc | 61 | 45 | 27 | 21 |
| **NC_028824.1** | Alphacoronavirus | Rhinacovirus | BtRf-AlphaCoV/YN2012 | 135 | † | | | | | | | |
| **NC_076697.1** | Alphapironavirus | Samovirus | PCNV-H14 | 111 | † | | | | | | | |
| **NC_028814.1** | Alphacoronavirus | Decacovirus | BtRf-AlphaCoV/HuB2013 | 42 | † | | | | | | | |
| **NC_022103.1** | Alphacoronavirus | Colacovirus | BtCoV-CDPHE15 | 80 | † | | | | | | | |

^ Locations were determined according to secondary structures proposed in the literature (1, 12). When a species was not explicitly modeled, sequence alignment was used to identify a proposed SL5 region.

† After aligning sequences, the SL5 region was uncertain. Highest sequence similarity was found in the 5' proximal region, such that SL5 should extend beyond the first nucleotide.

* Human-infecting coronavirus.

**Supplemental Table S2: Alpha- and betacoronavirus reference genomes' SL5 sequence and predicted secondary structure.**

| NCBI reference genome | SL5 sequence and EternaFold predicted secondary structure |
|---|---|
| NC_012936.1 | UUCCUUGACUUUCGUUCUCUGCCAGUGACGUGUCCAUUCGGCGCCAGCAACCCACCCAUAGGUUGCAUAAUGGCAAAGAUGGGCAAAUACGGUCUCGGCUUCAAAUGGGCCCCAGAAUUUCCAUGGAUGCUUCCGAACGCAUCGGAGAAGUUGGGUAACCCUGAGAGGUCAGAGGAGGAUGGGUUUUGCCCCUCUGCUGCGCAAGAACCGAAAGUUAAAGGAA<br>(((((((((((((((.((((((((((((((((.((...(((((((((........))))))....))))...))))))))....)).)(((((((((....(((.(((((..(((((...(((((........))))))))))..)))))...)))...)))))).)))(((...(((.....)))))))))))).))))))).))))))).)))))) |
| NC_001846.1 | UUCCUUGACUUUCGUUCUCUGCCAGUGACGUGUCCAUUCGGCGCCAGCAGCCCACCCAUAGGUUGCAUAAUGGCAAAGAUGGGCAAAUACGGUCUCGGCUUCAAAUGGGCCCCAGAAUUUCCAUGGAUGCUUCCGAACGCAUCGGAGAAGUUGGGUAACCCUGAGAGGUCAGAGGAGGAUGGGUUUUGCCCCUCUGCUGCGCAAGAACCGAAAGUUAAAGGAA<br>(((((((((((((((.((((((((((((((((.((...(((((((((........))))))....))))...))))))))....)).)(((((((((....(((.(((((..(((((...(((((........))))))))))..)))))...)))...)))))).)))(((...(((.....)))))))))))).))))))).))))))).)))))) |
| NC_048217.1 | UUCCUUGACUUUCGUUCUCUGCCAGUGACGUGUCCAUUCGGCGCCAGCAGCCCACCCAUAGGUUGCAUAAUGGCAAAGAUGGGCAAAUACGGUCUCGGCUUCAAAUGGGCCCCAGAAUUUCCAUGGAUGCUUCCGAACGCAUCGGAGAAGUUGGGUAACCCUGAGAGGUCAGAGGAGGAUGGGUUUUGCCCCUCUGCUGCGCAAGAACCGAAAGUUAAAGGAA<br>(((((((((((((((.((((((((((((((((.((...(((((((((........))))))....))))...))))))))....)).)(((((((((....(((.(((((..(((((...(((((........))))))))))..)))))...)))...)))))).)))(((...(((.....)))))))))))).))))))).))))))).)))))) |
| AC_000192.1 | UUCCUUGACUUUCGUUCUCUGCCAGUGACGUGUCCAUUCGGCGCCAGCAGCCCACCCAUAGGUUGCAUAAUGGCAAAGAUGGGCAAAUACGGUCUCGGCUUCAAAUGGGCCCCAGAAUUUCCAUGGAUGCUUCCGAACGCAUCGGAGAAGUUGGGUAACCCUGAGAGGUCAGAGGAGGAUGGGUUUUGCCCCUCUGCUGCGCAAGAACCGAAAGUUAAAGGAA<br>(((((((((((((((.((((((((((((((((.((...(((((((((........))))))....))))...))))))))....)).)(((((((((....(((.(((((..(((((...(((((........))))))))))..)))))...)))...)))))).)))(((...(((.....)))))))))))).))))))).))))))).)))))) |
| NC_006577.2 | UUCCACACUUUUCAUCUCUCUGCCAGUGACGUGUUGGUUGUCCUCAGCGUCCCUCCCAUAGGUCGCAAUGAUUAAAACCAGCAAAUACGGUCUCGGCUUCAAGUGGGCGCCAGAAUUUCGUUGGCUGCUUCCGGAUGCAGCGGAGGAGUUGGCUAGUCCUAUGAAGUCAGAUGAGGGUGGGUUAUGCCCCUCUACUGGUCAAGCGAUGGAAAGUGUUGGAU<br>.((((((((((((((.((.(((((((.((((((((((...((((((.((.(.....))).))).))..))))...)))))))....)))((((.((((((...(((((((..(((((....((((........))))...)))))))))..))))..)))))))))))(((((...(.....))))))))))))).))).)))))))))))).))). |
| NC_026011.1 | UUUGACUGAUAACACUUCCCUGCUAACGGCGUGUUGCACUCUCAGUCUAAGCCUCCCACCCAUAGGAGGUAUCAUGGCUAAGCGAGUCAACAAAUACGGCCUCACGCUAAAAUGGGCCCCAGAAUUUCCGUGGAUGUUUGAGGACACAGAGGAGAAGUUGGGUAACCCUGAUAGCGGAGAUGGGGAUGGUUUUGCUCCCCAGUUAAGCAGGAAGAGGGAUCGGUCAAAG<br>(((((((((..(.(((.((((((((((((((((((.((((.(((((...((((((........))))).....))))..).))))))))....)).)))((.(((((....(((.(((((..(((((.((..((((....)))))).)))))..)))))...)))...)))))))..((((((.(.......)))))))))).)))))))).)..))))))))). |
| NC_017083.1 | UUCCAUUUGUUAACAGCUUUCAGCCGGUGACGUGUUGUAUCCUAGGCAGUGGCCCACCCAUAGGUCAUAAUGCCGAAGAUCAACAAAUACGGUCUCGAGCUGCAAUGGGCUCCAGAAUUUCCGUGGAUGUUUGAGGACACAGAGGAGAAGUUGGACUACCCUAGCAGUUCAGAAGUGGGUAUGGUUUGCCCCACUACCGCGCAAAAGCUGGGAUCAAGUGGAA<br>((((((((.(..((((((..((((((((((((((.(((.(.((((((((((........))))))..)))).).))))))))....))).))(((((((((.(.(((.(((((..(((((.((..((((....)))))).)))))..)))))...)))).)))))))).))(((((((((.....)).)))))))))).)).)))))))..).))))))))) |
| NC_006213.1 | UUCUGCUGUUAACAGCUUUCAGCCAGGGACGUGUUGUAUCCUAGGCAGUGGCCCGCCCAUAGGUCACAAUGUCGAAGAUCAACAAAUACGGUCUCGAACUACACUGGGCUCCAGAAUUUCCAUGGAUGUUUGAGGACGCAGAGGAGAAGUUGGAUAACCCUAGUAGUUCAGAGGUGGAUAUGAUUUGCUCCACCACUGCGCAAAAGCUGGAAACAGACGGAA<br>(((((((((..((((((..(((((((((((((((.(((.(.((((((((((........))))))..)))).).))))))))....))).))))((((((...(((.(((((..(((((.((..((((....)))))).)))))..)))))...)))..))))))).(.((((((..........)))))).))).)).)))))))..))))).))))) |
| NC_003045.1 | UUCUGCUGUUAACAGCUUUCAGCCAGGGACGUGUUGUAUCCUAGGCAGUGGCCCACCCAUAGGUCACAAUGUCGAAGAUCAACAAAUACGGUCUCGAACUACACUGGGCUCCAGAAUUUCCAUGGAUGUUUGAGGACGCAGAGGAGAAGUUGGACAACCCUAGUAGUUCAGAGGUGGAUAUAGUAUGCUCCACCACUGCGCAAAAGCUGGAAACAGGCGGAA<br>(((((((((..((((((..(((((((((((((((.(((.(.((((((((((........))))))..)))).).))))))))....))).))))((((((...(((.(((((..(((((.((..((((....)))))).)))))..)))))...)))..))))))).(.((((((..........)))))).))).)).)))))))..))))).)))))) |
| NC_002306.3 | GUUUGGCAAUCACUCCUAGGAACGGGGUUGAGAGAACGGCGCACCAGGGUUCCGUCCCUGUUUGGUAAGUCGUCUAGUAUUAGCUGCGGCGGUUCCGCCCGUCGUAGUUGGGUAGACCGGGUUCCGUCCUGUGAUCUCCCUCGCCGGCCGCCAGGAGAAUGAGUUCCAAAC<br>(((((.((((((((((.((..(((((..((((..(((((..((((((......)))))....))..))))((((...((((((((((((((......)))))))))))))).)))).((((......)))))..))))))...)))....))))))..))).)))))))) |
| NC_038861.1 | GUUUGGCAAUCACUCCUUGGAACGGGGUUGAGCGAACGGUGCAGUAGGGUUCCGUCCCUAUUUCGUAAGUCGCCUAGUAGUAGCGAGUGCGGUUCCGCCCGUACAACGUUGGGUAGACCGGGUUCCGUCCUGUGAUCUCCCUCGCCGGCCGCCAGGAGAAUGAGUUCCAAAC<br>(((((.((((((((((.((..(((..(((.(((((((....(((((((......))))))).)))...((((.........))))((((((......))))))......(((.((((((((......)))).).))))))))))))))))))))))))..))).)))))))) |
| NC_028806.1 | GUUUGGCAAUCACUCCUUGGAACGGGGUUGAGCGAACGGUGCAGUAGGGUUCCGCCCCUAUUUCGUAAGUCGCCUAGUAGUAGCGAGUGCGGUUCCGCCCGUACAACGUUGGGUAGACCGGGUUCCGUCCUGUGAUCUCCCUCGCCGGCCGCCAGGAGAAUGAGUUCCAAAC<br>(((((.((((((((((.((..(((..(((.(((((((....(((((((......))))))).)))...((((.........))))((((((......))))))......(((.((((((((......)))).).))))))))))))))))))))))))..))).)))))))) |
| NC_030886.1 | AAGCUUUUGUGUCUUCAGCUUCCCUCUUCUCCUCGUGGGGUUCCGUUCCCGCGGUAGUGAGCCUGGUGUGGUUCCGCCCGCGCCAGUGAUGGAGGGUGCUUUAGGCACUAACAAGCUU<br>(((((((((((((..(((..(((((((((((((((((((((......))))))))..).)))((((((((((......))))))))).)).)))))).))).)))))).)).)))))) |
| NC_009021.1 | GAGCUUUUUGGUUAAGUACUUCCCCCUUGCGAGCAACAGGUGUCGUCCUGUUUGUUAGCCCUGCGCGGUUUCGUCCGUGUAGUGAUGGAGGGUGUGCCUGACCCACCUAAGCUU<br>((((((..((((((.((((..(((((..((.((((((((((......)))))).))).))((((((((((......)))))))).)..)).))).)))).))))).)...)))))) |
| NC_048211.1 | UUGCUCUUUGAGUGUUGCACAAGUGUGGUGUUUUUGUGCUAAGUUCUGUUUAGCACGAAAACCUUAAGUCUAGUUCCGUCUGGCUUAAGUGUGAAACUGUUGUUCCGUCUGCAGUGAAUCCCAUAACUGGCAACAUGUCUGUGAGCAA<br>((((((...(((((((((.(.(((((((.((((((((((((((.....))))))))))))))(((((((.((......)))))))))...(..(((((.(......).)))))....))))).))))))))))).))...)))))) |
| NC_035191.1 | UUGCUCUUUGAGUGUUGCACAAGUGUGGUGUUUUUGUGCUAAGUUCUGUUUGGCACGAAAACCUUAAGUCUAGUUCCGUCUGGCUUAAGUGUGAAACUGUUGUUCCGUCUGCAGUGAAUCCCAUAACUGGCAACAUGUCUGUGAGCAA<br>((((((...(((((((((.(.(((((((.((((((((((((((.....))))))))))))))(((((((.((......)))))))))...(..(((((.(......).)))))....))))).))))))))))).))...)))))) |
| NC_010437.1 | ACAUACUUGCACAAGUGUCGUGCAUGUCGCCAGUCCCUCCUUUCAGUUCCGUCUGUUAGGUAUACUAGGUGGCUGCCUUUGGUUCAGUUCCGUCUGGCCAUUGUGUGGAUAGUACGUUCCGUCGUGCUUGAAACCGAUAACUAGCAGGUAUGU<br>((((((((((...((((((.(((.((((((((....(((..(((......)))..)))...))).)))))))))(..((((.(((......)))))))..).(.....((((((......)))))).....))))).))).)))))))))) |
| NC_010438.1 | CCAUCCCGGCCCCAGUGUCGUGUCAUUCAGUGGUCCCUCGCACUGUUCCGUCAGUGCUUAAACCACUGUCUGACCUGCACGUAGUUCUGUCUGCGUGCAGUGUGGAUAGUACGUUCCGUCGUGCUUGAAACCGAUAACAGGUCGCUAUGG<br>((((..(((((...((((((.((((..(((((((.....((((((......))))))....)))))))..))))(((((((((((......)))))))))).((....((((((......))))))....)))))).)).))))).)))) |
| NC_054015.1 | GGUUGUCUAUUGGUUGGUUAAGUGCUGAGUGGCCUUUGACCUCUUUGUUGGUGUUCCGUCACCUUCAUAUGUCAUAGUGACACCCUCUUUCUGUUCCAUCAGUUAGAGUGAGGAUUCCAGGUUCUGUCUUGGUUGAAUUCAGCAACUGCUAAUCAUGUCUAGCAACC<br>(((((.(((.((((((((..((((((((((.(((.((((.....((..(((((......)))..))...)))).)).).)).(((((..(((......)))..))))....).(.(((((......)))))....))))))).)))))))))))....)))))))) |
| NC_046964.1 | GGUUGCUGAUUAAGUUGUUAAGUGUAGUGCGUCCUUGGACCUCUGCAGGGUGUUCCGUCACUCUGUACGUUUGGGGUAACGCCUUACUCACUGUUCCGUCAGUGGGUUUGUGUGGAUACUGGGUUCCGUCUCAGUUGAAACCUACAACUGCAACUCAUGGCUCGCAACC<br>(((((.((.(.((((((..(((((((.(((((((((((...(((((((((......)))))))))).))))))))..)))))..((((((((......))))))))......((..((((((......))))))....))))).)))))))))).)....)))))))) |
| NC_028833.1 | GGUUGCUGAUUAAGUUGUUAAGUGUAGUGGGUUCAUACACCUCUUCUGGUUAGUUCCGUCUACCAGUAAGUGUGUGAUACUCCCCUCUUCUGUUCUGUCAGUCGAGGUGUGUGGAUACCAGGUUCCGUCUUGGUUGAAACCUAUAACUGCAACUCAUGGCUCGCAACC<br>((((((.((...((((((..(((...(((((((((((((...(((((((.((......))))))).))))))))))).((.(.((((..(((......)))..)))).).))....((((((......))))))....)))))).)))))))))......)))))))) |
| NC_028811.1 | GGCUGCCCCAUUUUGCUUAAAAGUAGUGUGCCUCGUACACUCCCUGCCGUAGUGUUCCGUCACUACAGAUCACAGUGUGCCUGGACCCUCUACUGUUCCGUCAGUGGAGGUGUGUGGAUACAGUGUUCCGUCGCUGUUGAAACCACUAACAUAAGCAAUGGCUUGCAGUC<br>((((((.(((..((((((((...((((....((..(((((((....((.((((((......)))))).)..)..)))))))..))((((((((((......)))))))))).)).(((..((((((......))))))).....)))))).).))))))))))...)))))) |
| NC_009988.1 | GGUUGCCAGCCUUGUGUAACAAGUGUCGUGCUUCUAGGGGUUGCACUUGACGUGUUCCGUCACGUACUUGUGUCCUUAGUUGCCCCACGCUGUUCUGUCAGUGUGGUGUGUGGAUAUAGUGUUCCGUCACUAUUGAAACCGAUAACUGUGCACAAUGUCUAUCAACC<br>(((((..(((.((((((.((.(((((((.((..((((((...(((..((((((......))))).)..))))))))))..))(((((((((......))))))))).).((....((((((......))))))....))))))).)))))))))).).)).)..))))) |
| NC_054003.1 | GCUUAAAUAGUCUAAGCAAGUGCUGUGCUGUUAGUUUGUUCUUGGUGGAGGUUCCGUCCUCCAUUACAUACUUACUACUCAGCCUCCUCACGGUUCCGUCCUUGAGGUGUGUGGAUACUCUGUUCUGUCAGGGUUGAAACCAGUAACUGCCGACUAUGUCUAGC<br>(((...((((((...((.(((((((.((((.((((..((...(((((((((......)))))))))...)).)))))..))))(.(((((.((......)).))))).).((....((((((......))))))....)))))).)))))).))))))....))) |
| NC_009657.1 | GGUUUGCUUAGUAGCUAGUACAAGUGUCGUGCUGCUCCAGGGUUCUUGGUGUGGGUUCCGUCUCACACUUCACACCCUCAAAGUAGCCUUCCUGCAGUUUUGUCUGCAGGUUGUGUGGAAAGUCUGUCCGUCAGACUUGAAACCGAUAACUGUUAGCUAUGGCAAGCAACC<br>(((.(((((.((((((((..(.(((((((.((((((..(((((....((((((((......))))))))....)))))...))))))(..(((((((......)))))))..).((...(((((((......)))))))...)))))).)))))))))))))...)))))))) |
| NC_076629.1 | GGUUUGCUUAGUAGCUAAGUAAGUACUGUGCUGUCCUCAGCUCUUGUGGUGUGUUCCGUCACACUAACAAUCUGAUAAACAGCCUUCUCUCCUGUUCCGUCAGGGGAGUUGUGUGGAUAGUGUAUUCCGUUACACUUGAAACCAGCAACUGCCAGCUAUGAACAGCAACC<br>(((.((((..((((((..((.((....(.(((((..((((...((((((((((......)))))).)))).)))).)))...))))))..((((((((......))))))))((((.(((..((((((......)))))).....))))))))))).))))))....))))))) |

| NCBI reference genome | SL5 sequence and EternaFold predicted secondary structure |
|---|---|
| NC_003436.1 | GGUUUGCUUAAGUAGCCAUCGCAAGUGCUGUGCUGUCCUCUAGUUCCUGGUUGGCGUUCCGUCGCCUUCUACAUACUAGACAAACAGCCUUCCUCCGGUUCCGUCUGGGGGUUGUGUGGAUAACUAGUUCCGUCUAGUUUGAAACCAGUAACUGCCGGCUAUGGCUAGCAACC<br>(((.((((...((((((...((.(((((((.(((((..((((((...(((..((((......))))..)))...))))))...)))))(..(((((((......)))))))..)....(..((((((......)))))).....))))).))))).))))))....))))))) |
| NC_054004.1 | GCCAAAGUUUGGUAACAAGUGUCGUGCGUCCUAGUAGCUCUGGUUAGGUGUUCCGUCACCAUCCAUCUACUACAAACGCCCUUUGGCAGUUCCGUCUGCUAGAGUGUGUGGAAGACGAGUUCUAGCUUGUCUGAAACCGACAACUGUACCAUAUGGC<br>(((...(.((((.((.(((((((.((((..((((((...(((.(.((((......))))).))).))))))...))))(((((((((......))))))))).).((...((((((((....))))))))...)))))).))))))))).))))) |
| NC_018871.1 | GCCUAAUUUGGGUAACAAGUGCAGUGCGUCCUAGUGGCUCUGAGUGAGGGUUCCGUCCCCUCUCAUCCAUUACAAACGCCCUUUGCCAGUUCCGUCUGGUGAAGUGUGUGGAAGUCAAGUUCCGUCUUGUCUGAAACCUGUAACUGUGCCCAAUGGC<br>(((..(.((((((.((.(((((((.((((..((((((...((((..((((.......)))))))).))))))...))))(((((((((......))))))))).).((...((.((((......)))).))...)))))).))))))))))))))) |
| NC_076684.1 | GACUGUCUGUUGGCUUGCUAAGUGUCGUGGGUGUUCAGACCUCUGUUAGGGUGUUCCGUCACUCUUUCACGUUUGGAUUACUCCCCUGCGCAGUUCCGUCUGUGCAGGUGUGUGGAUAUCAAGUUCCGUCUUGGUUGAAACCGAUUACUCCAAGCCAUGUCAAGCAGUC<br>(((((.((.(((((((...((((((...((((((((((...((..((((((......))))))..)).))))))))(((.(.(((((((((......))))))))).).)))...((((((......))))))....)))))).))).)))))))...)).)))))) |
| NC_076685.1 | GAUUGUGUAUAUAAGGGCUCAAGUGCUGUGCCUCUGUUGUGUUCCUAGUGAGGUGUUCCGUCACCCACUACACAUAACCUAGGCCCCUCUGCAGUUCCGUCUGUAGAGGUGUGUGGAUAUCAAGUUCCGUCUUGGUUGAAACCAGUAACUGUUCCUUAUGGCAAACAAUC<br>(((((((.((((((((..(.(((((((.((((..(((((((...(((((.((((......)))))))))).)))))))..))))((((((((((......))))))))).).((....((((((......))))))....)))))).)))).))))))))))..)))))) |
| NC_028752.1 | GGUUGCAACAGUUUGGAAAUAAGUGCUGUGCGUCCUAGUCUAAGGGUUCUGUGUUCUGUCACGGGAUUCCAUUCUACAAACGCCUUACUCGAGGUUCUGUCUCGUGUUUGUGUGGAAGCAAAGUUCUGUCUUUGUGGAAACCAGUAACUGUUCCUAAUGGCCUGCAACC<br>(((((((.(.(((.((((.(.(((((((.(..((((......)(((((((((((......))))))))))..(((((((.......((.(((((.....))))).))))..))))))(((((((......)))))))))..))))).)))))))).))))...))))))) |
| NC_002645.1 | GGUUGCAACAGUUUGGAAGCAAGUGCUGUGUGUCCUAGUCUAAGGGUUUCGUGUUCCGUCACGAGAUUCCAUUCUACAAACGCCUUACUCGAGGUUCCGUCUCGUGUUUGUGUGGAAGCAAAGUUCUGUCUUUGUGGAAACCAGUAACUGUUCCUAAUGGCCUGCAACC<br>(((((((.(.(((.(((.((.(((((((.((.(((........((((((((((......))))))))))..(((((((.......((.(((((.....))))).))))..))))))(((((((......)))))))))).)))))).)))))))).))))...))))))) |
| NC_055953.1 | GCCGGUCUGCGUUAGGAAGGCAAGUGUAGUGCUUGCUGUAAGCUCUAGUGCAGUGUUCCGUCACUGUACUAUCUUGCAGUAAGACCCAGCCAGUUCUGUCUGGUUGGUGUGUGAAAGCAACGUUCUGUCGUUGUGGAAACUUACAACUGCUCCUAAUGGCACACGGGC<br>(((.((.((((((((((..((.(((((((..((((((((((...((((((((((......)))))))))).))))))))))((((((((((......)))))))))).))((....((((((......))))))....)))))).))))))))))))).))).)).))) |
| NC_048216.1 | UGUAUAAUUGAGGGAGCAAGUGUUGUGUGACAGCUGUCUAAGGGCUGGGUGUUCCGUCACCUAGUUCCAUUCAGUUGGAACCUAGUCGGUGUUCCGUCACUGGCUAGUGUGGAAGCAGUGUUCCGUCGCUGUGGAAACCAACAACUGUCCCUCAUGGCUUACA<br>((((....(((((((.(.(((((((.((.(((((((.....((((((((((......))))))))))....)))))))..((((((((((((......))))))))))).)).....((((((......))))))....)))))).))))))))))).....)))) |
| NC_032107.1 | GGUUCGUAAAGUGGUUAGGCAAGUGUUGUGUGCCCUGUGUUUAAGUACUGGUGGUUCCGUCCACUGGUGCACUCGCUUGGCACCUAAGCAGUGUUCCGUCAUUGCUUAGUGUGGAAACAACGUUCUGUCGUUGUGGAAACCAAUAACUGCUAACCAUGUCCUACAACC<br>((((.(((..((((((((.(.(((((((.(((((..(((.....((((((((((......))))))))))...)))..)))))(((((((((((......)))))))))))).((....((((((......))))))....)))))).)))))))))))))....))))))) |
| NC_005831.2 | UUGUAAACUGGUUAGGCAAGUGUUGUAUUUUCUGUGUCUAAGCACUGGUGAUUCUGUUCACUAGUGCAUACAUUGAUAUUUAAGUGGUGUUCCGUCACUGCUUAUUGUGGAAGCAACGUUCUGUCGUUGUGGAAACCAAUAACUGCUAACCAUGUUUUACAA<br>((((((((((((((.(.(((((((...((((((((((....((((((((((......)))))))))))..)))).......(((((((((......)))))))))........((((((......)))))))))))).)))).))))))))))).))).))))) |
| NC_034972.1 | GGUUUGCACGUCUUGGAGGAAACGGUGUGUACGUUUCUAGUGUUUACGCCGUAUCGGUUCCGGCCCGAUAGGUAUUGCAUUAGACGUCCUGGGUGGUUCUGCCUGCCCUUGUGUGAUUCGGCUGUUCCGUCAGUUUGGUCACCUCACACGUCCUUAAGACAUGGCUAACC<br>((((.(((.(((((.((((..(((.((((.(((..((((((((..(.(((.((((((.......)))))))))).).))))))))))))...((((((......))))))..(.((((((.(((((......))))).))))))).)))))))))))))))).).)).)))) |
| NC_032730.1 | GGUUCGCACGUCUUGUAGGAAACGGUGUGUACUUUUAGUGUUUACGCUUGUAUCGGUUCCGCCCGAUAGGUAUUGCACUAACAGUCUUGUGGGUUGUUCCGACCUUCCCCUUGUGUGGCACUCCUAUUCCGACUAGUGAGGUCACCACACACGUCCUUUAGACAUGGCCAACC<br>((((.(((.((((...(((..((((((((.(((.(((((((.(((..(..((((((......)))))))))))..)))))))).)))...(.(((((.....))....)))).....(((((.((((((.......))).)))))))))))).))))))..)))).).)).)))) |
| NC_025217.1 | GAUUCAUAGUACAUCCUUGUCUAGCUUUUGGUUGGCAGUGAGCCUACGGUUUCGUCCGUGUCGCUCACAAUUAUCCACACAGUAGGUUUCGUCCGCUGUGGUUGAGUUGCUAGUCCGUUGCUGUUUCGUCAGCCAUCUACAACUCGACACCAUGUGUAGCAAGGGGUAUGAACUGAAUC<br>((((((..((((.((((((.(((((...(((((((..((((((.(((((......)))))..))))))......)))((((((.((......))))))))(((((((((.(((...(..((((......)))))..)))))))))))))).)).))))))))))))....)))))) |
| NC_030292.1 | GCUUGUUGUCCAUAUUGAGUCUAGGAGUUUGGCUAGCUGUGUGGGUUCCGUCUCACAUAGUAGGUGUGUUGGCUAGUUCCGUCUAGUCUUCACGUGGAUCUCAGUUCCGUCUGGGUGAAACCUAACCCCCCUCACUAUGGUGUUCAAGC<br>(((((....(((((.((((....((.(((.((((.((((((((((......)))))))))))(((((..((((((......))))))..)))))....(((((......)))))....)).))).)).)))).)))))....))))) |
| NC_023760.1 | UUAUCUAGCUUGUUAACCAUUCUGAGGUAGGGGUUUGGCUGGCUGAGCGUGUUCCGUCACGCUUAGUAGGUGCUUCAGCUAGUUCCGUCUAGCUGUUGCGUGGAUCCUAGUUCCGUCUAGGUGAAACCUUACUCCCCUCAUUAUGGGAUCCAAGCACUUUAA<br>.......(((((....((((..(((((..(((((..((((.((((((((((......)))))))))).))(((..(((((((......)))))))..)))(....(((((......)))))....)))..)))))))))..))))....)))))...... |
| NC_004718.3 | UCGUUGACAAGAAACGAGUAACUCGUCCCUCUUCUGCAGACUGCUUACGGUUUCGUCCGUGUUGCAGUCGAUCAUCAGCAUACCUAGGUUUCGUCCGGGUGUGACCGAAAGGUAAGAUGGAGAGCCUUGUUCUUGGUGUCAACGA<br>((((((((((.((((((...(((.(((.(((((((..((((((.(((((......)))))..))))))......)))(((((((.((......))))))))(((....))))))).)))))).)))))))))...)))))))) |
| NC_045512.2 | UCGUUGACAGGACACGAGUAACUCGUCUAUCUUCUGCAGGCUGCUUACGGUUUCGUCCGUGUUGCAGCCGAUCAUCAGCACAUCUAGGUUUCGUCCGGGUGUGACCGAAAGGUAAGAUGGAGAGCCUUGUCCCUGGUUUCAACGA<br>(((((((((((((..(((...(((.(((((((((((..((((((.(((((......)))))..))))))......)))(((((((.((......))))))))(((....)))))))))))))).))))).)))...))))))) |
| NC_014470.1 | UCGUUGACAGGACACGAGUAACUCGUCUAUCUUCUGCAGGCUGCUUACGGUUUCGUCCGUGUUGCAGCCGAUCAUCAGCAUACCUAGGUUUCGUCCGGGUGUGACCGAAAGGUAAGAUGGAGAGCCUUGUCCCUGGUUUCAACGA<br>(((((((((((((..(((...(((.(((((((((((..((((((.(((((......)))))..))))))......)))(((((((.((......))))))))(((....)))))))))))))).))))).)))...))))))) |
| NC_038294.1 | UUUUCAGUUAGAGCGUCGUGUCUCUUGUACGUCUCGGUCACAAUACACGGUUUCGUCCGGUGCGUGGCAAUUCGGGGCACAUCAUGUCUUUCGUGGCUGGUGUGACCGCGCAAGGUGCGCGCGGUACGUAUCGAGCAGCGCUCAACUCUGAAAA<br>((((((((.((((((..((.(((..(((((((((((((((..(((.(((......)))))).)))))....)))).(((((((.(((......)))))))))))((((((((.......)))))))))))))).)))))))))))))..))))))) |
| NC_019843.3 | UUUUCAGUUAGAGCGUCGUGUCUCUUGUACGUCUCGGUCACAAUACACGGUUUCGUCCGGUGCGUGGCAAUUCGGGGCACAUCAUGUCUUUCGUGGCUGGUGUGACCGCGCAAGGUGCGCGCGGUACGUAUCGAGCAGCGCUCAACUCUGAAAA<br>((((((((.((((((..((.(((..(((((((((((((((..(((.(((......)))))).)))))....)))).(((((((.(((......)))))))))))((((((((.......)))))))))))))).)))))))))))))..))))))) |
| NC_009019.1 | UUUUCAUUCUGAGCUAUCGUGUCUCUUGUACUUCUCGGUCACAAUACCCUGUUCCGUCAGGUGCGUGGCAAUCCGGGGCAUACUAUGCUUUCCAAGGCUAGUGUGACCACGCAAGGUGCGCGUGGUAAGUAUCGAGCAGAGCUCUAUAAUGAAAA<br>((((((((..(((((..(.((.(((..((((((((((((((((....(((((......)))).).))))....))))).(((((((.((((....)))))))))))((((((((.......))))))))))))).))))))))))))...)))))))) |
| NC_039207.1 | UUUUCAUUUAGAGCUGCGUGUCUCAAGUGCGUCUCGGUCACAAUAUACUGUUCCGUCAGGUGCGUGACAAUCCGGGGCUCAUCAUGUCUUCCGCGACUGGUGAGGGUUCUCAAGGUGCGAGGGCCACGUAUAGAGCAGCGCUCAACAAUGAAAA<br>((((((((..((((.((.((.(((..((((((((((((((...((.(((......)))))..))))....))))).(((((((.(((......)))))))))))((((((((.......))))))))))))).))))))))))))...)))))))) |
| NC_009020.1 | UUUUCAGUUAGAGCAUCGUGUCUCAAGUGCUUCACGGUCACAAUAUACCGUUUCGUCGGGUGCGUGGCAAUUCGGUGCACAUCAUGUCUUUCGUGGCUGGUGUGGCUCCUCAAGGUGCGAGGGGCAAGUAUAGAGCAGAGCUCAACACUGAAAA<br>((((((((.((((.((.((.(((..((((((.(((((........)))))..((((((((((...)).))))))))(((((((.(((......)))))))))))((((((((.......))))))))))))).))))))))))).).)))))))) |

**Supplemental Table S3: Stem length, resolution, and modeling convergence of the cryo-EM maps of the SL5 domain of coronaviruses.**

| Species | Conformation | Estimated number of helical turns of stem in cryo-EM map | | | | Estimated length of stem in cryo-EM map (Å) | | | | Resolution (Å) | Modeling convergence (Å) |
|---|---|---|---|---|---|---|---|---|---|---|---|
| | | SL5-stem | SL5a | SL5b | SL5c | SL5-stem | SL5a | SL5b | SL5c | | |
| SARS-CoV-2 | - | 1 ¾ | 1 ½ | 1 | ½ | 52 | 54 | 36 | 18 | 4.7 | 2.47 |
| SARS-CoV-1 | - | 2 ½ | 1 ½ | ¾ | ½ | 75 | 54 | 32 | 18 | 7.1 | 3.16 |
| MERS | Conformation 1 | 1 ½ | † | 1 | ¾* | 53 | 58^ | 34 | 39* | 6.9 | 3.46 |
| | Conformation 2 | 1 ½ | 1 ½^ | 1 | ¾* | 55 | 56^ | 35 | 28 | 6.4 | 3.23 |
| | Conformation 3 | 1 ½ | † | 1 | ¾* | 55 | 58 | 33 | 32 | 6.4 | 3.35 |
| BtCoV-HKU5 | Conformation 1 | 1 ½ | 1 ½^ | 1 | ¾* | 56 | 58^ | 32 | 26* | 5.9 | 2.99 |
| | Conformation 2 | 1 ½ | 1 ½^ | 1 | ¾* | 55 | 52^ | 34 | 27* | 6.4 | 3.06 |
| | Conformation 3 | 1 ½ | † | 1 | † | 48 | 51 | 30 | 30* | 8.0 | 5.18 |
| | Conformation 4 | 1 ½ | † | 1 | ¾* | 50 | 44 | 30 | 30* | 7.3 | 2.99 |
| HCoV-229E | - | 1 ¼ | 2 ¼ | 1 | 1 ¼ | 40 | 67 | 30 | 40 | 6.5 | 2.34 |
| HCoV-NL63 | Conformation 1 | † | † | † | † | 64 | 53 | 40 | 26 | 9.0 | † |
| | Conformation 2 | † | † | † | † | 58 | 43 | 34 | 25 | 8.4 | † |

^ A bend in this helix distorts the A-form structure; the number of turns and length before and after the bend are summed.
* End of stem is involved in a tertiary interaction and thus deviates from the A-form helix.
† Helical grooves not resolved sufficiently.

**Supplemental Table S7: Cryo-EM sample preparation conditions for the SL5 domain and SL5-6 domains of coronaviruses.**

| | SARS-CoV-2 SL5 collection 1 | SARS-CoV-2 SL5 collection 2 | SARS-CoV-2 SL5 collection 3 | SARS-CoV-2 SL5 collection 4 | SARS-CoV-2 SL5-6 | SARS-CoV-2 SL5-6 SL5b extended | SARS-CoV-2 SL5-6 SL5c extended | SARS-CoV-2 SL5-6 SL6 extended | SARS-CoV-1 SL5 | MERS SL5 | BtCoV-HKU5 SL5 collection 1 | BtCoV-HKU5 SL5 collection 2 | HCoV-229E SL5 | HCoV-NL63 SL5 |
|---|---|---|---|---|---|---|---|---|---|---|---|---|---|---|
| **RNA length (nt)** | 124 | | | | 196 | 204 | 204 | 129 | 143 | 135 | 135 | | 140 | 160 |
| **Molecular weight (kDa)** | 40.03 | | | | 63.11 | 65.70 | 65.70 | 41.36 | 46.14 | 43.67 | 43.73 | | 45.10 | 51.43 |
| **Concentration (µM)** | 20 | 30 | 30 | 30 | 20 | 20 | 20 | 20 | 30 | 30 | 30 | 30 | 30 | 25 |
| **Concentration (mg/mL)** | 0.80 | 1.20 | 1.20 | 1.20 | 1.26 | 1.31 | 1.31 | 0.83 | 1.38 | 1.31 | 1.31 | 1.31 | 1.97 | 1.28 |
| **Refolding conditions** | RNA denatured at 90°C for 3 min with Na-HEPES (50mM final concentration). RNA cooled at room temperature for 10 minutes. $MgCl_2$ (10 mM final concentration) added, and RNA folded at 50°C for 20 minutes. RNA cooled at room temperature for 10 minutes. RNA is immediately blotted or stored on ice before freezing. | | | | | | | | | | | | | |
| **Grid type** | Quantifoil R 2/1 grids (Cu, 200 mesh) | Quantifoil R 1.2/1.3 grids (Cu, 300 mesh) | Quantifoil R 1.2/1.3 grids (Cu, 300 mesh) | Quantifoil R 1.2/1.3 grids (Cu, 300 mesh) | Quantifoil R 2/1 grids (Cu, 200 mesh) | Quantifoil R 1.2/1.3 grids (Cu, 300 mesh) | Quantifoil R 1.2/1.3 grids (Cu, 300 mesh) | Quantifoil R 1.2/1.3 grids (Cu, 300 mesh) | Quantifoil R 1.2/1.3 grids (Cu, 300 mesh) | Quantifoil R 1.2/1.3 grids (Cu, 300 mesh) | Quantifoil R 1.2/1.3 grids (Cu, 300 mesh) | Quantifoil R 1.2/1.3 grids (Cu, 300 mesh) | Quantifoil R 1.2/1.3 grids (Cu, 300 mesh) | Quantifoil R 1.2/1.3 grids (Cu, 300 mesh) |
| **Glow discharge** | 30 s glow discharge with air, 15 s wait | | | | | | | | | | | | | |
| **Freezing conditions** | 4 s blot, 5 s wait | 4 s blot, 5 s wait | 4 s blot, 5 s wait | 4 s blot, 5 s wait | 4 s blot, 5 s wait | 4 s blot, 1 s wait | 4 s blot, 1 s wait | 2.5 s blot, 1 s wait | 2.5 s blot, 1 s wait | 2.5 s blot, 1 s wait | 4 s blot, 1 s wait | 4 s blot, 1 s wait | 4 s blot, 1 s wait | 4 s blot, 1 s wait |

**Supplemental Table S8: Cryo-EM data collection parameters for the SL5 domain and SL5-6 domains of coronaviruses.**

| | SARS-CoV-2 SL5 collection 1 | SARS-CoV-2 SL5 collection 2 | SARS-CoV-2 SL5 collection 3 | SARS-CoV-2 SL5 collection 4 | SARS-CoV-2 SL5-6 | SARS-CoV-2 SL5-6 SL5b extended | SARS-CoV-2 SL5-6 SL5c extended | SARS-CoV-2 SL5-6 SL6 extended | SARS-CoV-1 SL5 | MERS SL5 | BtCoV-HKU5 SL5 collection 1 | BtCoV-HKU5 SL5 collection 2 | HCoV-229E SL5 | HCoV-NL63 SL5 |
|---|---|---|---|---|---|---|---|---|---|---|---|---|---|---|
| **Microscope** | Titan Krios G3i | Titan Krios G3i | Titan Krios G3i | Titan Krios G3i | Titan Krios G3i | Titan Krios G3i | Titan Krios G3i | Titan Krios G3i | Titan Krios G3i | Titan Krios G3i | Titan Krios G3i | Titan Krios G3i | Titan Krios G3i | Titan Krios G3i |
| **Voltage (kV)** | 300 | 300 | 300 | 300 | 300 | 300 | 300 | 300 | 300 | 300 | 300 | 300 | 300 | 300 |
| **Energy filter** | No | No | No | No | No | Selectris | BioQuantum | Selectris | BioQuantum | No | No | No | Selectris | Selectris |
| **Energy filter slit width (eV)** | - | - | - | - | - | 10 | 20 | 10 | 20 | - | - | - | 10 | 10 |
| **Detector** | Falcon 4 | Falcon 4 | Falcon 4 | Falcon 4 | Falcon 4 | Falcon 4 | K3 | Falcon 4 | K3 | Falcon 4 | Falcon 4 | Falcon 4 | Falcon 4 | Falcon 4 |
| **Nominal magnification** | 96,000 | 96,000 | 96,000 | 96,000 | 75,000 | 165,000 | 130,000 | 165,000 | 105,000 | 96,000 | 96,000 | 96,000 | 165,000 | 165,000 |
| **Pixel size (Å)** | 0.82 | 0.82 | 0.82 | 0.82 | 1 | 0.741 | 0.68 | 0.741 | 0.86 | 0.82 | 0.82 | 0.82 | 0.741 | 0.741 |
| **Electron exposure ($e^-$ /Å²)** | 50.00 | 50.00 | 50.00 | 49.99 | 50.00 | 50.12 | 50.00 | 49.88 | 50.00 | 49.97 | 49.97 | 49.79 | 50.12 | 60.00 |
| **Dose per frame ($e^-$ /Å²/frame)** | 1.25 | 1.25 | 1.25 | 1.25 | 1.25 | 1.25 | 1.25 | 1.25 | 1.25 | 1.25 | 1.25 | 1.24 | 1.25 | |
| **Frame duration (s/frame)** | 0.075 | 0.096 | 0.103 | 0.139 | 0.075 | 0.117 | 0.030 | 0.115 | 0.015 | 0.119 | 0.119 | 0.095 | 0.117 | 0.090 |
| **Exposure time (s)** | 6.00 | 3.84 | 4.13 | 5.56 | 8.50 | 4.66 | 1.19 | 4.60 | 0.59 | 4.74 | 4.74 | 3.81 | 4.66 | 3.61 |
| **Defocus range** | -1.2 to -2.5 | -1.2 to 2.2 | -1.0 to -2.0 | -1.0 to -2.0 | -1.2 to -2.5 | -1.0 to -2.0 | -1.0 to -2.0 | -1.0 to -2.0 | -1.0 to -2.0 | -1.0 to -2.0 | -1.0 to -2.0 | -1.2 to 2.2 | -1.0 to -2.0 | -1.2 to -2.0 |
| **Micrographs acquired** | 11,815 | 4,217 | 4,592 | 5,540 | 1,763 | 11,496 | 5,875 | 16,511 | 10,376 | 13,244 | 5,728 | 12,411 | 16,022 | 8,304 |

**Supplemental Table S9: Cryo-EM data processing details for the SL5 domain and SL5-6 domains of coronaviruses.**

| | SARS-CoV-2 SL5 | | | | SARS-CoV-2 SL5-6 | SARS-CoV-2 SL5-6 SL5b extended | SARS-CoV-2 SL5-6 SL5c extended | SARS-CoV-2 SL5-6 SL6 extended | | | | SARS-CoV-1 SL5 | MERS SL5 | | | BtCoV-HKU5 SL5 | | | | HCoV-229E | HCoV-NL63 | |
|---|---|---|---|---|---|---|---|---|---|---|---|---|---|---|---|---|---|---|---|---|---|---|
| **Micrographs used (% of acquired)** | 10,708 (91%) | 3,646 (86%) | 4,236 (92%) | 5,397 (97%) | 1,699 (96%) | 10,026 (87%) | 5,234 (89%) | 14,608 (88%) | | | | 8,929 (86%) | 9,517 (72%) | | | 14,633 (81%) | | | | 13,730 (86%) | 5,752 (69%) | |
| **Symmetry imposed** | C1 | | | | | | | | | | | | | | | | | | | | | |
| **Particle box sizes*** | 360 | | | | 420 | 512 | 256-fourier cropped 3x | 128 | | | | 380 | 360 | | | 200 | | | | 216-fourier cropped 2x | 230-fourier cropped 2x | |
| **Initial particles images** | 1,123,449 | 411,306 | 379,733 | 607,258 | 923,003 | 810,035 | 813,979 | 1,321,089 | | | | 1,734,565 | 1,207,909 | | | 5,109,283 | | | | 782,829 | 920,960 | |
| | 1,562,978 (particles combined after some initial 2D class averaging) | | | | | | | | | | | | | | | | | | | | | |
| **Particles images after 2D classification (% of initial)** | 707,462 (63%) | 172,645 (42%) | 205,122 (54%) | 477,749 (79%) | 529,912 (57%) | 535,028 (66%) | 269,712 (33%) | 495,099 (37%) | | | | 264,674 (15%) | 462,317 (38%) | | | 1,574,865 (31%) | | | | 660,081 (84%) | 376,856 (41%) | |
| | 649,371 (42%) | | | | | | | | | | | | | | | | | | | | | |
| **Final particle images (% post 2D)** | 392,855 (56%) | 72,713 (42%) | 64,842 (32%) | 157,498 (33%) | 99,785 (19%) | 199,745 (37%) | 65,937 (24%) | 79,134 (16%) | 71,615 (14%) | 84,922 (17%) | 85,939 (17%) | 220,917 (83%) | 81,341 (18%) | 97,886 (21%) | 106,867 (23%) | 163,382 (10%) | 189,981 (12%) | 127,548 (8%) | 185,489 (12%) | 160,535 (24%) | 111,416 (30%) | 106,304 (28%) |
| | 445,981 (69%) | | | | | | | | | | | | | | | | | | | | | |
| **Map resolution (Å)** | 4.9 | 6.8 | 7.4 | 6.6 | 7.8 | 7.4 | 9.1 | N/A | N/A | N/A | N/A | 7.1 | 6.9 | 6.4 | 6.4 | 5.9 | 6.4 | 8.0 | 7.3 | 6.5 | 9.0 | 8.4 |
| | 4.7 | | | | | | | | | | | | | | | | | | | | | |
| **FSC threshold** | 0.143 | | | | | | | | | | | | | | | | | | | | | |
| **Sharpening B-factor (Å²)** | -250 | | | | -400 | -600 | -600 | N/A | | | | -600 | -500 | -400 | -400 | -550 | -650 | -600 | -800 | -500 | -550 | -500 |

*Additional box sizes often used during processing.

**Supplemental Note S1: Sequence identity of SL5 from the studied coronaviruses.**

The sequence identity is defined here as after alignment of the sequences using a global alignment with no gap penalties and a score of 1 for a match, the number of matches divided by the length of alignment. The gap-excluded identity is defined here as the number of matches divided by the length of sequence.

The sequences are as follows:

- SARS-CoV-2 SL5: NC_045512.2 residues 150-294
- SARS-CoV-1 SL5: NC_004718.3 residues 149-293
- MERS SL5: NC_019843.3 residues 196-349
- BtCoV-HKU5 SL5: NC_009020.1 residues 178-331
- HCoV-229E SL5: MF542265.1 residues 139-307
- HCoV-NL63 SL5: NC_005831.2 residues 136-297

Mean pairwise gap-excluded identity across 6 coronaviruses SL5s imaged: 72.4%
Mean pairwise sequence identity across 6 coronaviruses SL5s imaged: 54.3%

SARS-CoV-2 : SARS-CoV-1, gap-excluded identity = 92.4%, sequence identity = 85.9%

```
UCGUUGACAG-GACA-CGAGUAACUCGUCUA--UCUUCUGCAGG-CUGCUUACGGUUUCGUCCGUGUUGCAGC-CGAUCAUCAGCACAU-C-UAGGUUUCGUCCGGGUGUGACCGAAAGGUAAGAUGGAGAGCCUUGUCC-CU-GGUU-UCAACGA
|||||||||  || | |||||||||||||    ||||||||| | |||||||||||||||||||||||||||  ||||||||||  ||| | ||||||||||||||||||||||||||||||||||||||||||||||   || || | |||||||
UCGUUGACA-AGA-AACGAGUAACUCGUC--CCUCUUCUGCA-GACUGCUUACGGUUUCGUCCGUGUUGCAG-UCGAUCAUCAG--CAUACCUAGGUUUCGUCCGGGUGUGACCGAAAGGUAAGAUGGAGAGCCUUGU--UCUUGG-UGUCAACGA
```

SARS-CoV-2 : MERS, gap-excluded identity = 71.0%, sequence identity = 52.6%

```
UCGUUGA-CAG---GA-CA--CGA-GUAACUCGUCUA-U-CU-UCUGCAGGCUGCU----UAC--GGUUUCGUCCGUGUUGCAGCCGAU--CA-U-CAG---CACAUCUAG-GU-UU-CGUCCGG--G-UGUGACCGA---AAGGUA---------A-G-AUG-GAG-AGCCUUG-UC--C-CUGGUUUCAACGA-
|  ||   |||   || |   ||  ||  ||| | |  | |  ||| | || | |     |||  ||||||||||| | |   | || |  || | | |   |||||| |  || || |||  ||  | ||||||||    |||||          | | ||  ||| || |  | ||  | || |    ||  |
U--UU--UCAGUUAGAGC-GUCG-UGU--CUC-U-U-GUAC-GUCU-C-GG-U-C-ACAAUACACGGUUUCGUCCG-G-U---G-CG-UGGCAAUUC-GGGGCACAUC-A-UGUCUUUCGU--GGCUGGUGUGACCG-CGCAAGGU-GCGCGCGGUACGUAU-CGAGCAG-C--GCUCAACUCU-G----AA--AA
```

SARS-CoV-2 : BtCoV-HKU5, gap-excluded identity = 69.7%, sequence identity = 51.0%

```
UCGUUGA-CAGGAC--ACGAGUAAC-UCGU--CU-A--U-CUUCUGCA-GGCUGC----U-UACG-GUUUCGUCCGUGUUGCA-GCCGAU--CA-U-CAG---CACAUCUAG-GU-UU-CGUCCGG--G-UGUGA-C-CGA---AAGGU---A-----A-G-AU-G-G-AGAGCCUUGUC--C-CUGGUUUCAACGA-
|  ||   || |    | |||   | ||||  || |  | | | | || || | |    | |||  ||||||| || |  |   | || |  || | | |   |||||| |  || || |||  ||  | ||||  | |     |||||   |     | | || | | |||| |   ||  | || |    ||  |
U--UU--UCA-G--UUA-GAG---CAUCGUGUCUCAAGUGC-U-U-CACGG-U-CACAAUAUAC-CGUUUCGU-CG-G--G--UG-CG-UGGCAAUUC-GGUGCACAUC-A-UGUCUUUCGU--GGCUGGUGUG-GCUC--CUCAAGGUGCGAGGGGCAAGUAUAGAGCAGAG-C---UCAACACU-G----AA--AA
```

SARS-CoV-2 : HCoV-229E, gap-excluded identity = 73.1%, sequence identity = 51.0%

```
UCG-UUG-A-CAG---G-ACACG-A-GUAA-CUCGUCUA-UC-UUCU-G-C-A-GGCUGCUUA-CGGU-UUC-GUC-CGU-G-UUGCAGCCGAU-CAU-CA---GCACAUCUA-G-GU---UUC-GUC-CG-G---GUGUG-ACC-G-AAAG----GUAA----GAUGGAGAG-CC--U---UGUC-CCU---GGUU--U-CAACGA-
  | ||| | |||   | | | | | ||   || |  |  |  |||| | | | ||  | ||  | || ||| ||| ||  | ||    || || | | ||   || | | || | |    ||| ||| || |   ||||| |   | ||||    ||      | |||| |  ||  |   |||  |||   ||    | ||||
--GGUUGCAACAGUUUGGA-A-GCAAGU--GCU-G--U-GU-GUUCUAGUCUAAGG--G-UU-UC-GUGUUCCGUCACG-AGAUU----CC-AUUC-UACAAACGC-C-U-UACGCG-AGGUUCUGUCUCGUGUUUGUGUGGA--AGCAAAGUUUUGU--CUUUG-UGGA-A-ACCAGUAACUGU-UCCUAAUGG--CCUGCAAC--C
```

SARS-CoV-2 : HCoV-NL63, gap-excluded identity = 69.0%, sequence identity = 48.3%

```
UCGUUGAC-AGGACAC-GAGUA-A--C---UCGUCU--AUCUUCUGCA-G-G-CU--GCUUAC-GGUUUCG-UC-CGUGUUGCAGCCG-A-U-CAU-CA--GCAC-AUCU-A-G-G-U-UUC-GUC-CGG-G-U----GUGACCGAA--A--G----GUAA-GAU-G-GAGAG--CC--U---UGUC---CC-UGGUUUCA-ACGA-
   |||   |  | || | ||  |  |   | || |  || || | |  | | ||  ||  || ||  | | |  | |||| ||  |  | | ||| ||  | |  || | | | | | ||| ||| |   | |    |||   |||  |  |    ||   | | | | ||   ||  |   || |   || | ||||   || |
---UUG--UA--A-ACUG-GU-UAGGCAAGU-GU-UGUAU-UU-U-C-UGUGUCUAAGC--ACUGG--U-GAU-UC-UGUU-CA--C-UAGUGCAUACAUUG-A-UAU-UUAAGUGGUGUUCCGUCAC--UGCUUAUUGUG---GAAGCAACGUUCUGU--CG-UUGUG-GA-AACCAAUAACUG-CUAACCAU-GUUU--UAC-AA
```

### SARS-CoV-1 : MERS, gap-excluded identity = 70.3%, sequence identity = 51.8%

```
U---C-GUU-GA-CAAGAAA-CGA-GUAACUC--GUCC-C-UCUUCUGCAG--AC--UG-CUUACGGUUUCGUCCGUGUUGCAGUCGAU-CA-U-CAG---CAUACC-U-AG-GU-UU-CGUCCGG--G-UGUGACCGA---AAGGUAAGAU-G-GA-GAGCCUU--GU-UCUUG-GU--G---UCAAC---GA---
|   | ||| || |  |     ||  ||  |||  ||    | || || |  |  ||  |  |  |||||||||||| | ||| || |    || | | |    |  | | | |  || || |||  ||  | ||||||||    |||||  |    | |  | |   |  || ||  | |    |    |||||   ||
UUUUCAGUUAGAGC--G---UCG-UGU--CUCUUGU--ACGUC-UC-G--GUCACAAU-AC--ACGGUUUCGUCCG-G-UGC-GU-G--GCAAUUC-GGGGC--A-CAUCA-UGUCUUUCGU--GGCUGGUGUGACCG-CGCAAGGU--G--CGCG-CG-G---UACGUAUC--GAG-CAGCGCUCAACUCUGAAAA
```

### SARS-CoV-1 : BtCoV-HKU5, gap-excluded identity = 67.6%, sequence identity = 48.8%

```
UCGUUGA-CAAGAA--ACGAGUAAC-UCGUCCCUCU-UCUGCAGAC-UGCUU-ACGGU-----U-U-C-GUC--CGU-GU--UGCAGUCG--A-U-CA--U-CAGCAUACCUAG-GU-UU-CGUCCGG--G-UGUGA-C-CGA---AAGGUAAG--AUGGAGAGCCUU--GUUCU-U-G-GU--G---UCAAC---GA---
|  ||   | ||    | |||   | |||       | ||| || |  ||||| |||||      | | | ||    ||| |    ||| || |  | | |    | || |||  | |  || || |||  ||  | ||||  | |      |||||  |  | || | | |     |    | | | |   |    |||||   ||
U--UU--UC-AG--UUA-GAG---CAUCG------UGUCU-CA-A-GUGCUUCACGGUCACAAUAUACCGU-UUCGUCG-GGUGC-GU-GGCAAUUC-GGUGCA-CAU--C-A-UGUCUUUCGU--GGCUGGUGUG-GCUC--CUCAAGGU--GCGA-GG-G-G-C--AAG---UAUAGAG-CAGAGCUCAACACUGAAAA
```

### SARS-CoV-1 : HCoV-229E, gap-excluded identity = 73.8%, sequence identity = 51.7%

```
UCG-UUGACAAGAAACGAGU----AA-C---U-C-GUCCC-UC-UUCUGCAGACUGCUUAC-GGU-UUCGUCCGUGUUGCA-GUC--GA--U-C-AU-C-AGCAUA-C-C-U-A-G-GU---UUC-GUC-CG-G---GUGUG-ACC-G-AAAG----GUAA----GAUGGAGAG-CC--U---UGUUC-U--UGG--UGUCAACGA-
  | ||| |     ||| |||     || |    | | ||     |  ||||  ||  | | ||  ||  ||  | |||||| |  |||  ||   | | || | | || | | | | | | |     ||| ||| || |   ||||| |    | ||||     ||        | |||| |  ||  |    ||||| |  |||  || ||||
--GGUUG-C----AAC-AGUUUGGAAGCAAGUGCUGU---GU-GUUCU--AG--U-C-UA-AGG-GUU--U-CGUGUU-C-CGUCACGAGAUUCCAUUCUA-CA-AACGCCUUACGCG-AGGUUCUGUCUCGUGUUUGUGUGGA--AGCAAAGUUUUGU--CUUUG-UGGA-A-ACCAGUAACUGUUCCUAAUGGCCUG-CAAC--C
```

### SARS-CoV-1 : HCoV-NL63, gap-excluded identity = 67.6%, sequence identity = 46.9%

```
UCGUUG-ACAA--G---A---AACGA-GUAACUCGUCCC-UCUU-CUGCA-GA-CU--GCUUAC-GGU--UUC-GU-C-CGU-GUUGCAGUCGAUCAUCA-GCAUACCU--A-G-G-U-UUC-GUC-CGG-G-U----GUGACCGAA--A--G----GUAA-GAU-G-GAGAG--CC--U---UGUUCU-----UGGUGUCA--ACGA-
   ||| | ||  |    |    || |  ||    | ||     | || |||    |  ||  ||  || |||  ||| || | | | | |||| |  | |||    | |||   |   | | | | ||| ||| |    | |     |||    |||  | |     ||    | | | | ||   ||  |    ||  ||       | || |     || |
---UUGUA-AACUGGUUAGGCAA-G-UGU---U-GU---AU-UUUCUG--UG-UCUAAGC--ACUGGUGAUUCUGUUCAC-UAG-UGCA-U--A-CAU--UG-AUA--UUUAAGUGGUGUUCCGUCAC--UGCUUAUUGUG---GAAGCAACGUUCUGU--CG-UUGUG-GA-AACCAAUAACUG--CUAACCAU-GU-U--UUAC-AA
```

### MERS : BtCoV-HKU5, gap-excluded identity = 87.0%, sequence identity = 77.0%

```
UUUUCAGUUAGAGCG-UCGUGUCUCUUGUAC-GU-CU-C--GGUCACAAUAC-ACG-GUUUCGUCCGG-UGCGUGGCAAUUCGGG-GCACAUCAUGUCUUUCGUGGCUGGUGUGA-C-CGCG-CAAGGUGCGC-GCGGUA-C--GUAUC-GAGCAGC-GCUCAACU-CUGAAAA
||||||||||||||  |||||||||    |  || || |  ||||||||||  ||  ||||||| ||| |||||||||||| || |||||||||||||||||||||||||||||||  | | |  |||||||||  | ||    |  ||||  |||||| ||||||| |||||||
UUUUCAGUUAGAGC-AUCGUGUCUC----A-AGUGCUUCACGGUCACAAUA-UAC-CGUUUCGU-CGGGUGCGUGGCAAUUC-GGUGCACAUCAUGUCUUUCGUGGCUGGUGUG-GCUC-C-UCAAGGUGCG-AG-GG--GCAAGUAU-AGAGCAG-AGCUCAAC-ACUGAAAA
```

### MERS : HCoV-229E, gap-excluded identity = 67.5%, sequence identity = 47.5%

```
UU--UU-CA---GUUA-G-A-GC--GU-C-GUGUC-UCU-U-GUAC---G--UCU-CG-GU-CACAA-UACACG-GUU-UCGUCCGG-UG-CGUGG-CAA-----UU-CG-G-GGCACAU-CAUGUCUUUCGUGGCUG--GUGUGACCGC--GCAAG-GUGC---G-C---GC-GGUA--C--GUAUCGAGCA-G--CGCUCAACU--C-UGAA-AA--
    || ||   |||  | | ||  || | ||||  | | | | |   |  | | || || | |  | ||| | |  | |||   | | |   |||      || || | ||   | | |||| |||| | |   |||||   |   |||| ||       | |   |  || |  | |  |||   | |   |  | || || |  | ||   ||
--GGUUGCAACAGUU-UGGAAGCAAGUGCUGUGU-GU-UCUAGU-CUAAGGGU-UUCGUGUUC-C--GU-CACGAG--AU--UCC--AU-UC-U--ACAAACGCCUUACGCGAGG----UUC-UGUC--UCGU-G-U-UUGUGUG---G-AAGCAA-AGU--UUUGUCUUUG-UGG-AAACCAGUA---A-C-UGUUC-CU-AA-UGGCCUG--CAACC
```

### MERS : HCoV-NL63, gap-excluded identity = 67.5%, sequence identity = 49.1%

```
UUU-UCA----G-UUAGAGC--GUCGUGUC-UC-UUGUACGUCUCG-GUCACAAUA--CAC-GGU--UUC-GU-C-CGGUGC-GUGGCAAUUCGGGGCACAUCAUG-UC-UUUC--GUGGCUGGUGUGACCG-C---GCA--AG--GUGCGC--GC---G----GUACGUAUCGA-G----C-AG----C-GCUCAACUC-UGA----A-AA
 || | |    | |||| ||  || || |  | | |  ||| | ||  | | ||  |||  |||  ||| || | |  | | |    |||  || | |       ||||  || |  |||   |||| | || |  ||| |   ||     |   | ||| |   ||   |  | | |||  ||   |    | ||| ||| | ||      | ||
-UUGU-AAACUGGUUAG-GCAAGU-GU-U-GU-AUU-U---UCU-GUGU--C--UAAGCACUGGUGAUUCUGUUCAC--U--AGU-GC-A-U------ACAU--UGAU-AUUU-AAGUGG-U-GU-U--CCGUCACUGC-UUA-UUGUG-G-AAGCAACGUUCUGU-CGU-U-G-UGGAAACCA-AUAACUGCU-AAC-CAUG-UUUUACAA
```

### BtCoV-HKU5 : HCoV-229E, gap-excluded identity = 67.5%, sequence identity = 47.5%

```
UU--UU-CA---GUUA-G-A-GCA--U-C-GUGU---CU---C-AAGUG----C-U-U-CACGGUCAC-A-AU---AU---AC---CGU--UU-CGUCG-GGUG-C-GU--G-GCAAUUC-G-GUGC-ACAU-CAU--GUCUUUCGUGGCUGGU-GUGGCU---CC--UCAAGG-UG--CGAGGGGCAAGUAUA-GAGCAGAGCU-CAACACUGAAAA
    || ||   |||  | | |||  | | ||||   ||    | | ||| |    | | | | ||||| | ||   ||   ||   ||  || || || ||| | ||  | |    ||   | | |||  | |  ||   || ||| ||  || | ||||      ||  | ||   || |        |   || | | ||    || |||| |
--GGUUGCAACAGUU-UGGAAGCAAGUGCUGUGUGUUCUAGUCUAAG-GGUUUCGUGUUC-C-GUCACGAGAUUCCAUUCUACAAACG-CCUUACG-CGAGGU-UCUGUCUCGUG---UU-UGUGUG-GA-A-GCA-AAGU-UUU-GU--CU--UUGUGG--AAACCAGU-AA--CUGUUC------C---UA-AUG-GC----CUGCAAC-C------
```

### BtCoV-HKU5 : HCoV-NL63, gap-excluded identity = 65.6%, sequence identity = 47.0%

```
UUUUCAGUUAGAGCAUCGUG-UCU---CAAGUGCUUC--A----C-G-GUCACAAUAUA-C-C-G-U--UUCGUCGGGUGCGUGGCAAUUCG--GUGCACAU-CAUGUC--U-UUCGUG--GCUGGUGUGGCUCC-UCAAGG-UGCG--A--G-GG--GCAA-GUAUA--GA--G---------C-AGAG---CU-C-AACAC-UGA----A-AA
  ||  | || |  | | || | |   |||||| ||   |     | | || | | | | |||| |   ||   |  | | |   |||  ||| ||| |   | ||  |   | ||||||   ||| || |   |||    |  | ||  |||| || |   |    |          | | |    || | ||| | ||      | ||
--UU--G-UA-A--A-C-UGGU-UAGGCAAGUG-UU-GUAUUUUCUGUGU--C--UA-AGCACUGGUGAUUC-U---GU---U--C-A--C-UAGUG--CAUACAU-U-GAUAUU--U-AAG-UGGUGU---UCCGUC-A--CUGC-UUAUUGUGGAAGCAACGU-U-CUG-UCGUUGUGGAAACCA-A-UAACUGCUAAC-CAUG-UUUUACAA
```

### HCoV-229E : HCoV-NL63, gap-excluded identity = 75.9%, sequence identity = 59.1%

```
GGUUGC-AACAGUU-UGGA--AG-CAAGUGCUGUGUG-UU--CUAGU--CUAAG----GGU--UUCGUGUUC-C--GU-CA--CGA--GAUUCCAUUCUACAAACGCCUUACGCGAG-GUUCU-GUCU-CGUG-UU-UGUGUGGAAGCAAA-GUUU-UGUCU-UUGUGGAAACCAG-UAACUGUUCCUAAUGGCC-UG-----CAACC
  |||  || |    |||    || |||||| | |||  ||  || ||  |||||     |||  ||| ||||| |  || ||  | |   || |    ||| |    || |   |     | | ||||  |||   | || || | |||||||| || | || ||||  |||||||||||  ||||||   ||||   || ||      |||
--UUG-UAA-A---CUGG-UUAGGCAAGUG-U-UGU-AUUUUCU-GUGUCUAAGCACUGGUGAUUC-UGUUCACUAGUGCAUAC-AUUGA-U--AUU-U---AA-G---U----G-GUGUUC-CGUC-AC-UGCUUAU-UGUGGAAGC-AACG-UUCUGUC-GUUGUGGAAACCA-AUAACUG---CUAA---CCAUGUUUUACAA--
```

**Supplemental Note S2: Explanation of data processing flowcharts**.

All data processing was conducted in CryoSPARC 3.2.0 including 3D variability analysis and non-uniform refinement (13–15).

Titles for each dataset are at the top of each workflow and are designated by a black box with white text. Workflows that result in multiple conformations, rather than a single dominant conformation, are also labeled with a black box with white text, at the end of the workflow.

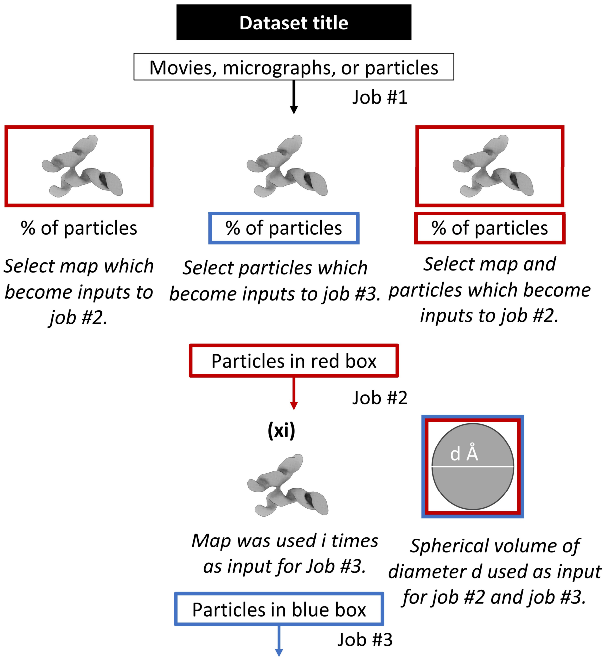

Quantities of movies, micrographs, or particles are boxed in black and connected by arrows, with the job name below the inputs. Jobs may generate one or multiple maps, with the percentage of input particles making up each output map labeled under each map.

Four types of items may be in a thicker, colored box: maps, spherical volumes, or particles. All of these serve as inputs to the job type with a similarly colored arrow. For some heterogeneous refinement jobs, spherical volumes are represented by a grey circle with the diameter labeled in Å. Any of these items described may have multiple thicker, colored boxes if they are used multiple times in the data processing workflow. For jobs that have multiple copies of the same map as input, the number of map copies is indicated above the map in bold.

See left for visuals of all workflow items described.
